# INSPIRE: interpretable, flexible and spatially-aware integration of multiple spatial transcriptomics datasets from diverse sources

**DOI:** 10.1101/2024.09.23.614539

**Authors:** Jia Zhao, Xiangyu Zhang, Gefei Wang, Yingxin Lin, Tianyu Liu, Rui B. Chang, Hongyu Zhao

## Abstract

Recent advances in spatial transcriptomics technologies have led to a growing number of diverse datasets, offering unprecedented opportunities to explore tissue organizations and functions within spatial contexts. However, it remains a significant challenge to effectively integrate and interpret these data, often originating from different samples, technologies, and developmental stages. In this paper, we present INSPIRE, a deep learning method for integrative analyses of multiple spatial transcriptomics datasets to address this challenge. With designs of graph neural networks and an adversarial learning mechanism, INSPIRE enables spatially informed and adaptable integration of data from varying sources. By incorporating non-negative matrix factorization, INSPIRE uncovers interpretable spatial factors with corresponding gene programs, revealing tissue architectures, cell type distributions and biological processes. We demonstrate the capabilities of INSPIRE by applying it to human cortex slices from different samples, mouse brain slices with complementary views, mouse hippocampus and embryo slices generated through different technologies, and spatiotemporal organogenesis atlases containing half a million spatial spots. INSPIRE shows superior performance in identifying detailed biological signals, effectively borrowing information across distinct profiling technologies, and elucidating dynamical changes during embryonic development. Furthermore, we utilize INSPIRE to build 3D models of tissues and whole organisms from multiple slices, demonstrating its power and versatility.

## Introduction

Spatial transcriptomic (ST) technologies enable spatially resolved transcriptomic studies by profiling gene expressions with spatial information in intact tissues [1, 2]. Recently, various ST technologies have been developed with complementary strengths [3, 4]. For example, widely used next-generation sequencing-based methods, such as 10x Visium [5], Slide-seq [6, 7] and Stereo-seq [8], allow for transcriptome-wide gene expression profiling. While technologies based on in situ hybridization (e.g., seqFISH [9, 10] and MERFISH [11]) and in situ sequencing (e.g., STARmap [12, 13]) require panel designs for target genes with prior knowledge, they offer single-cell and subcellular resolution that is essential in characterizing cellular communications. These diverse ST approaches provide great opportunities for deciphering complex tissue architecture [14, 15], understanding how cells interact with each other [16, 17], and identifying spatial developmental trajectories in tissues [18, 19].

Non-negative matrix factorization (NMF) has proven an appealing approach for analyzing transcriptomic count matrices [20, 21, 22]. For instance, in the context of single-cell RNA-sequencing (scRNA-seq) data analyses, NMF-based methods have the ability to decompose gene expression in individual cells into a set of interpretable gene programs associated with cell-type identities and cellular activities [23, 24]. These methods offer valuable insights, such as unraveling cell states that arise in various perturbations [25, 26]. Most recently, two NMF-based dimension reduction methods, SpiceMix [27] and NSFH [28], were proposed for analyzing complex ST data by capturing spatial dependence of cells. Owing to the decomposition nature of NMFs [29], SpiceMix and NSFH are powerful in deciphering signals within ST data by decomposing them into a collection of interpretable spatial factors, each encoding a unique spatial pattern [27, 28]. These methods excel at uncovering spatial organization of cell identities,identifying spatially variable features, and revealing important biological processes. However, these methods are designed only for interpreting a single ST dataset. The development of interpretable and spatially-aware analytical methods that can effectively integrate multiple diverse ST datasets is in great need and remains a challenge.

With advancements in ST technologies, numerous ST studies utilizing different technologies have been conducted, each often generating multiple slices [3, 4]. For instance, ST profiles have been characterized in multiple sagittal and coronal sections from mouse brains [30, 31, 32], and in multiple parallel slices along the left-right axis from a late-stage mouse embryo [8]. Multiple ST slices have also been created from mouse embryos at varying developmental time points [8, 33]. Effectively interpreting these diverse ST datasets, both within and across studies, is crucial for establishing a comprehensive understanding of tissue architectures and their developmental dynamics. However, unwanted variations across samples, batches, ST technologies and developmental time points can introduce confounding factors that hinder the discovery of biologically meaningful spatial signals [34, 35]. Consequently, while methods like SpiceMix and NSFH can unveil meaningful spatial factors in a single ST dataset, they face difficulties in distinguishing shared biological signals among datasets from the heterogeneous unwanted variation when applied to multiple ST datasets. This task can become even more challenging if certain datasets contain unique biologically meaningful spatial factors that need to be accurately identified and separated from the unwanted variation [36]. Therefore, there remains a need for computational methods that are specifically designed for joint analyses of multiple diverse ST datasets.

Here, we develop INSPIRE, a deep learning-based method that unifies NMF and adversarial learning [37] to achieve interpretable, flexible and spatially-aware integration of ST datasets. INSPIRE leverages graph neural networks [38, 39] to perform spatially informed analyses of ST slices, by accounting for local microenvironments of cells or spatial spots. For joint analyses of multiple datasets, INSPIRE incorporates a tailored adversarial learning mechanism to adaptively distinguish complex unwanted variations across multiple batches, samples, technical platforms and developmental stages from intrinsic biological variations, even when certain datasets present unique biological signals. Hence, INSPIRE can reliably eliminate heterogeneous unwanted variations in its analyses. By seamlessly integrating this adversarial learning mechanism with NMF, INSPIRE enables a harmonized NMF for multiple diverse ST datasets. It allows for the discovery of spatial factors among multiple datasets without confounded by unwanted variations, deciphering detailed spatial organizations in diverse datasets. For these spatial factors, INSPIRE also explicitly models their gene signatures, enabling the interpretation of their biological meanings and the identification of gene programs associated with them.

Through the application of INSPIRE to various ST datasets, including human cortex slices from different samples, mouse brain slices prepared at different orientations and resolutions, as well as a collection of whole mouse embryo slices, we demonstrate that INSPIRE can flexibly integrate diverse ST datasets from multiple samples, created by different ST technologies and at varying developmental time points. In these diverse integrative analyses, INSPIRE shows its power to decipher fine-grained spatial architecture with biological meanings, elucidate spatial cell-population distributions, and uncover biological processes organized in complex tissues. Through these applications, we show that INSPIRE is a versatile analytical approach that allows for various downstream analyses. For instance, it enables pathway enrichment analysis, identification of spatially variable genes, detection of spatial trajectories, imputation of spatial gene expressions as well as 3D reconstruction of tissue structures using multiple parallel slices along an axis. Of note, INSPIRE is also scalable to handle large-scale datasets. As a demonstration, we applied INSPIRE to spatiotemporal atlases of mouse organogenesis, comprising half a million high-resolution spatial spots. INSPIRE effectively modeled these atlases, deciphering dynamical changes during mouse embryonic development. INSPIRE is publicly available as a Python package (https://github.com/jiazhao97/INSPIRE), offering an efficient and reliable tool for ST data analyses.

## Results

### Method overview

INSPIRE takes gene expression count matrices and spatial coordinates from multiple ST slices as inputs. It effectively integrates information across slices in a shared latent space. In this space, meaningful biological variations from the input slices are preserved, while complex unwanted variations are eliminated (Fig. 1**a** and panel **a1**). Utilizing this shared latent space, INSPIRE achieves an integrated NMF on gene expressions across slices, decomposing biological signals in different slices into consistent and interpretable spatial factors with associated gene programs (Fig. 1**a** and panel **a2**). The non-negative spatial factors of cells or spatial spots are inferred from their latent representations, where unwanted variations are largely eliminated. Hence, these spatial factors are free from complex unwanted variations, enabling unified discoveries of fine-grained spatial patterns among slices. The gene signatures associated with these spatial factors are captured through shared non-negative gene loadings among slices, enhancing the interpretability of multi-slice integrative analyses.

**Figure 1:**
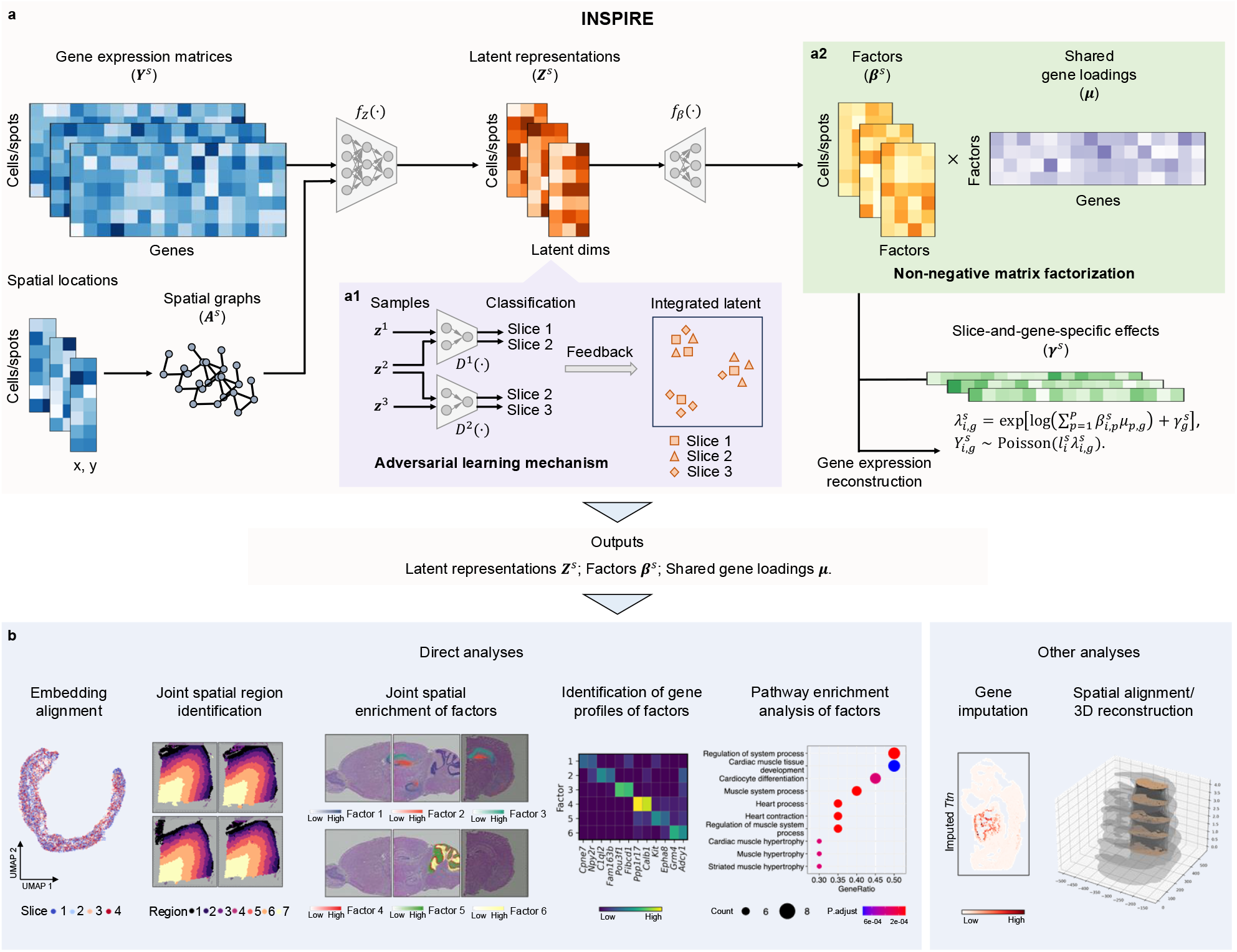
Overview of INSPIRE. **a.** INSPIRE is a unified deep learning method that incorporates adversarial learning and non-negative matrix factorization (NMF) for interpretable integrations of ST datasets. Raw gene expressions and spatial locations from multiple slices are taken as the inputs. (**a1**). INSPIRE embeds the biological variations from ST slices into a shared latent space. By incorporating a tailored adversarial learning mechanism, INSPIRE effectively eliminates unwanted variations in this latent space, providing harmonized representations of cells or spatial spots among slices. (**a2**). The latent space enables INSPIRE to achieve an integrated NMF for multiple slices, further decomposing biological signals into a set of consistent and interpretable factors among slices. Unconfounded by unwanted variations, these spatial factors reveal detailed spatial organizations in multiple ST slices. The gene signatures of these spatial factors are explicitly characterized by the shared gene loading matrix, elucidating their biological meanings. After training, INSPIRE simultaneously outputs integrated latent representations, interpretable spatial factors, and corresponding gene loadings. **b**. INSPIRE’s outputs enable multiple downstream analyses, including spatial trajectory inference, identification of fine-grained spatial regions and tissue structures, detection of spatially variable genes, and pathway enrichment analysis for deciphering biological processes in tissues. INSPIRE can also be applied to tasks including gene imputation and 3D reconstruction of tissues with multiple parallel 2D slices.

INSPIRE seamlessly incorporates the above designs in a unified deep-learning framework (Fig. 1**a**). Specifically, it uses a spatially-informed encoder, *f*_*Z*_(·), to map cells or spatial spots from ST slices *s* = 1, 2, *…, S* into the shared latent space. This encoder is a graph neural network [38, 39] that takes gene expressions and spatial neighborhood graphs of ST slices as inputs. For any cell or spatial spot *i* from slice *s*, the encoder is designed to output its latent representation 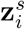, unaffected by unwanted variations. INSPIRE achieves this by incorporating a tailored adversarial learning mechanism [34, 37] (Fig. 1**a**, panel **a1**). To align 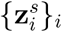 from slice *s* with 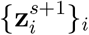 from slice *s* + 1, an auxiliary discriminator network, *D*^*s*^(·), is deployed in the latent space to detect where poor mixing between 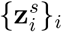 and 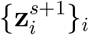 occurs. Its feedback then guides encoder *f*_*Z*_(·) to improve the alignment. The discriminator can adaptively preserve slice-specific signals by only guiding *f*_*Z*_(·) to integrate shared biological variations between slices. By introducing *S* − 1 discriminators, including *D*^*s*^(·), *s* = 1, 2, *…, S* − 1, INSPIRE effectively harmonizes all *S* slices in the shared latent space.

Next, INSPIRE adopts an integrated NMF across slices to further decompose the biological signals in the shared latent space into a set of interpretable spatial patterns with gene programs (Fig. 1**a**, panel **a2**). This provides characterizations of tissue structures at a finer-grained level and with enhenced interpretability. The integrated NMF includes non-negative spatial factor matrices {***β***^*s*^}_*s*_ for the multiple slices, and a shared non-negative gene loading matrix, ***μ***, among slices. For any cell *i* in slice *s*, 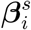 presents the set of non-negative weights across the hidden spatial factors it contains. The sum of the non-negative weights of spatial factors for any given cell equals to one. The contributions of different genes to diverse hidden spatial factors are explicitly encoded by the non-negative weights in ***μ***, revealing gene programs associated each detailed spatial pattern. Two unique designs in INSPIRE enable its integrated NMF across slices. First, INSPIRE uses a decoder network, *f*_*β*_(·), to generate spatial factors 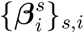 directly from integrated representations 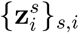 in the latent space. This ensures that 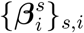 only decompose meaningful biological signals among slices, as the unwanted variations are removed in 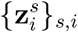. Second, INSPIRE introduces additional slice *s*-and-gene *g*-specific effects 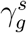 to help explicitly model confounding signals. Therefore, 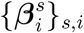 and ***μ*** can efficiently fit the biological signals, decomposing these key signals into a set of detailed spatial factors with their associated gene modules that are consistent among slices.

INSPIRE formulates the learning of encoder network *f*_*Z*_(·), decoder network *f*_*β*_(·), and gene loading matrix ***μ*** into a unified optimization problem. After training, INSPIRE simultaneously outputs integrated latent representations {**Z**^*s*^}_*s*_, spatial factors {***β***^*s*^}_*s*_, and interpretable gene loadings ***μ***. The outputs enable comprehensive characterizations of tissue structures through various downstream analyses (Fig. 1**b**). These include the identification of spatial trajectories and major spatial regions using {**Z**^*s*^}_*s*_; the discovery of detailed tissue architectures, spatial distributions of cell types, and the organization of biological processes using {***β***^*s*^}_*s*_; and the detection of spatial variable genes, the identification of gene modules, along with pathway enrichment analysis using ***μ***. Details are included in the Methods section.

### INSPIRE offers superior accuracy and interpretability for the integrative analysis of multiple ST datasets

In this section, we first demonstrate that by integrating adversarial learning with NMF for joint modeling of multiple ST slices, INSPIRE can achieve superior accuracy in capturing biological signals across slices. This advantage allows INSPIRE to produce improved results for the identification of spatial regions, which is critical in ST data analysis. Additionally, we highlight that INSPIRE can model gene programs that characterize detailed spatial organization patterns in tissues, enhancing the interpretability of multi-slice integrative analysis.

We applied INSPIRE to a human dorsolateral prefrontal cortex (DLPFC) dataset run on the Visium platform [40]. This dataset contains four DLPFC tissue slices, indexed 151673-151676, from a neurotypical adult donor. Researchers have manually annotated six DLPFC layers (L1-L6) and white matter (WM) for each slice based on cytoarchitecture and gene markers [40]. We first focused on analyzing the spot representations obtained in INSPIRE’s latent space.

In this space, information across slices was effectively integrated, while different cortical layers were still well separated (Fig. 2**a**). Importantly, the spot representations revealed a clear trajectory from L1 to L6 and WM that is shared among slices. The identified trajectory aligns with corticogenesis, during which cortical neurons are born in a successive order from outer to inner layers [41], showing INSPIRE’s ability to distill meaningful biological variation among slices. The effective preservation of biological signals in the latent space enables INSPIRE to reliably identify spatial regions in tissues. Using spot representations, INSPIRE effectively recovered the layer structures among all DLPFC slices (Fig. 2**c**). This result shows a consistent pattern compared to manual annotation, indicating INSPIRE’s high reliability in spatial domain identification (Fig. 2**b, c**).

**Figure 2:**
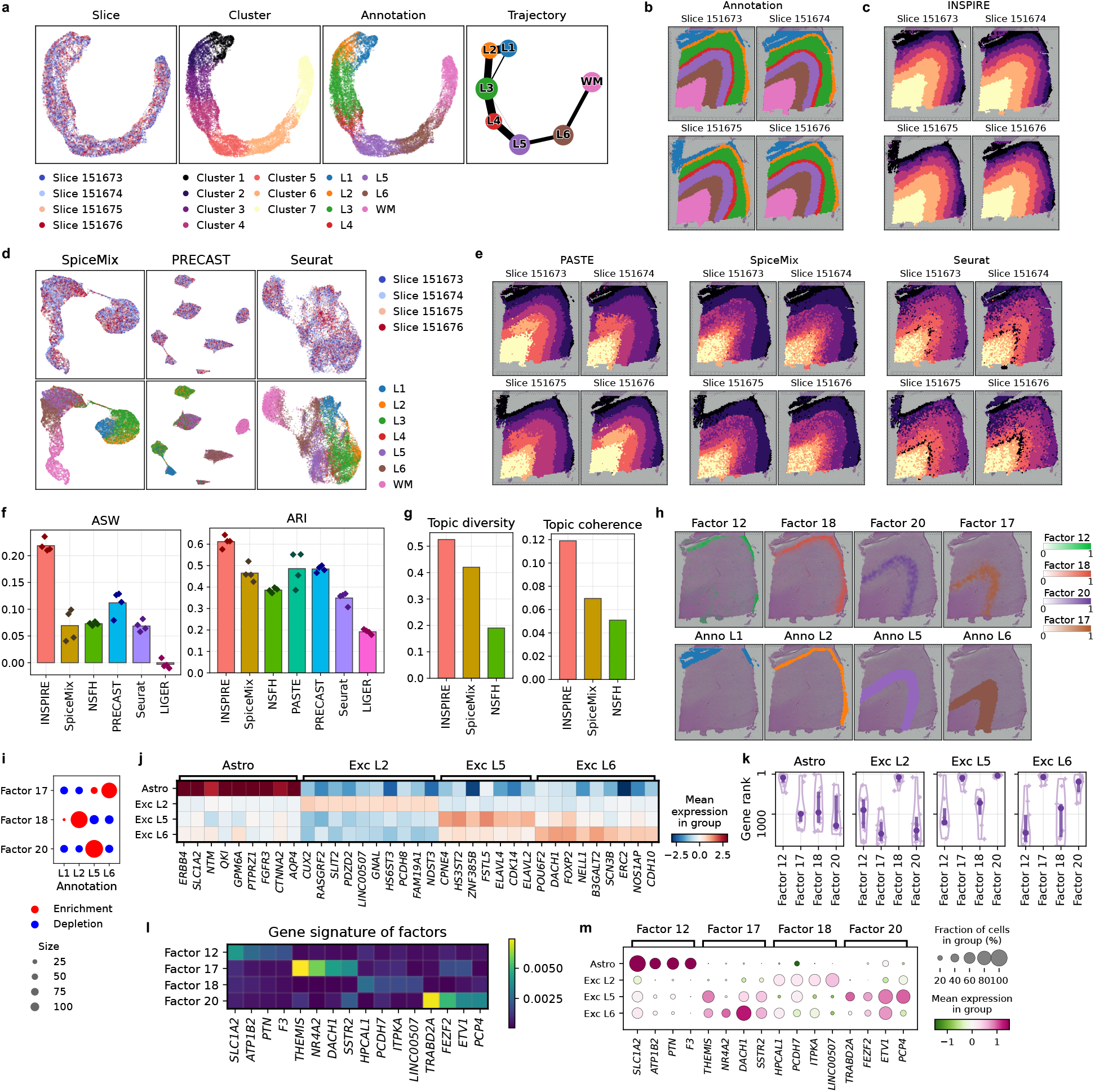
Benchmarking of INSPIRE and state-of-the-art methods based on the human DLPFC dataset. **a.** UMAP plots of spot representations from INSPIRE, colored by slice indices, INSPIRE’s assigned spatial domain labels, and manual annotations. The PAGA algorithm was applied to the spot representations for spatial trajectory inference. **b**. Manual annotations on slices. **c, e**. Spatial domain identification results from INSPIRE (**c**), PASTE, SpiceMix, and Seurat (**e**). **d**. UMAP plots of spot representations from SpiceMix, PRECAST and Seurat, colored by slice indices and manual annotations. **f**. ASW and ARI scores of the benchmarked methods. **g**. Factor diversity and factor coherence scores of the benchmarked methods. **h**. Spatial distributions of factors 12, 18, 20 and 17 identified by INSPIRE on slice 151673. **i**. Enrichment or depletion of different spatial factors in cortical layers. **j, k**. We identified marker genes for four cell types using a scRNA-seq atlas (**j**), and visualized the rank distribution of them in different spatial factors (**k**). **l**. Spatial factor-specific genes identified by INSPIRE. **m**. Expression levels of the factor-specific genes among cell types.

For a quantitative evaluation of INSPIRE’s spot representation and spatial domain identification, we used the manual annotations as ground truth and adopted three metrics: average silhouette width (ASW), adjusted rand index (ARI), and normalized mutual information (NMI). ASW measures the conservation of different annotated layers in spot representations, while ARI and NMI assess the accuracy of spatial domain identification by comparing it to the manual annotations. Higher scores on these metrics indicate better performance. To benchmark INSPIRE against existing tools, we compared it to representative state-of-the-art methods applicable for producing spot representations or identifying spatial domains in this analysis, including Seurat [42], LIGER [22], SpiceMix [27], NSFH [28], PRECAST [35], and PASTE [43]. INSPIRE achieved superior performance compared to all the other methods, reflected by the highest scores for all three metrics (Fig. 2**f** and Supplementary Fig. 2). In contrast, scRNA-seq data integration methods Seurat and LIGER showed less satisfactory results across all three scores due to their lack of consideration for spatial information (Fig. 2**e, f** and Supplementary Fig. 3). Two spatially-informed data integration methods, PASTE and PRECAST, had better performance compared to Seurat and LIGER, illustrating the importance of spatial coordinate modeling. However, PASTE could not provide spot representations across slices, and its spatial domain identification results were inconsistent across slices (Fig. 2**e**). PRECAST incorrectly mixed spots from layers L4 and L5, and showed limited ability to preserve the continuous trajectory among cortical layers (Fig. 2**d**). SpiceMix and NSFH are two spatially-aware NMF-based methods for ST data analysis, designed to handle one ST slice at a time. We manually concatenated the four slices to form one ST slice to apply these methods. Both methods successfully uncovered the spatial trajectory among layers, indicating their ability to capture biological signals (Fig. 2**d** and Supplementary Fig. 1). However, compared to INSPIRE, they both showed less satisfactory performance across all three metrics, suggesting their limited ability to leverage information across slices for achieving improved results (Fig. 2**f**, Supplementary Figs. 2 and 3).

**Figure 3:**
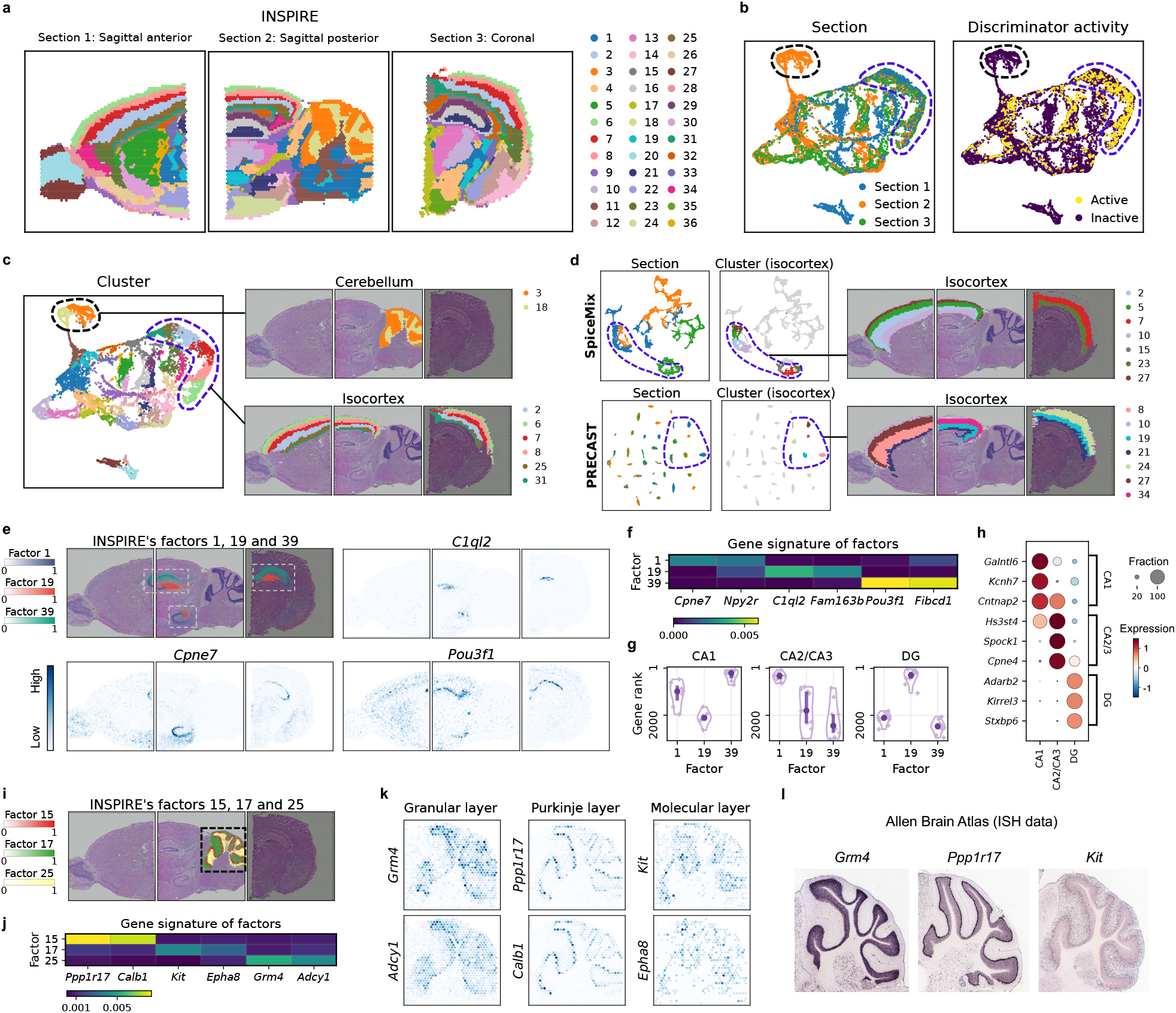
Analysis of multiple mouse brain ST slices with only partially shared spatial tissue organizations. INSPIRE integrated sagittal anterior, sagittal posterior and coronal sections of mouse brains. **a**. Spatial clusters visualized on the three ST slices. **b**. UMAP plots of spot representations from INSPIRE, colored by slice indices and discriminator activities. INSPIRE’s discriminators were active on the spots describing the shared spatial structures across slice, while they were inactive on spots related to the slice-unique structures. Thereby, they adaptively guided INSPIRE to align shared variations among slices, while preserving slice-unique signals. **c**. Spatial regions 3 and 18 from INSPIRE characterized the cerebellum. Spatial regions 2, 6, 7, 8, 25 and 31 characterized layers in the isocortex. **d**. UMAP plots of spot representations from SpiceMix and PRECAST, and visualizations of their spatial clusters in the isocortex. **e, f**. Spatial distributions of factors 1, 19 and 39 identified by INSPIRE (**e**). Based on the learned gene signatures (**f**), INSPIRE identified spatially variable genes associated with the three factors respectively (**e**). **g, h**. We identified marker genes of CA1, CA2/3 and DG using a scRNA-seq atlas (**h**), and visualized the rank distribution of them in gene loadings of the three factors (**g**). **i, j**. Spatial distributions (**j**) and gene signatures (**j**) of factors 5, 17 and 25 respectively. **k**. Top spatially variable genes associated with each of the three factors identified by INSPIRE. **i**. ISH images of genes *Grm4, Ppp1r17* and *Kit* from Allen Brain Atlas.

So far, we have demonstrated the superior performance of INSPIRE in learning spot representations. We note that the gained accuracy in the latent space is contributed by the NMF component in INSPIRE. For illustration, we manually removed the NMF component from INSPIRE, and denoted this version as “INSPIRE (w/o NMF)”. Compared to INSPIRE (w/o NMF), INSPIRE’s spot representation consistently showed higher scores for all three metrics ASW, ARI, and NMI (Supplementary Fig. 4). This demonstrates the effectiveness of INSPIRE’s integration of the shared latent space with the NMF model.

**Figure 4:**
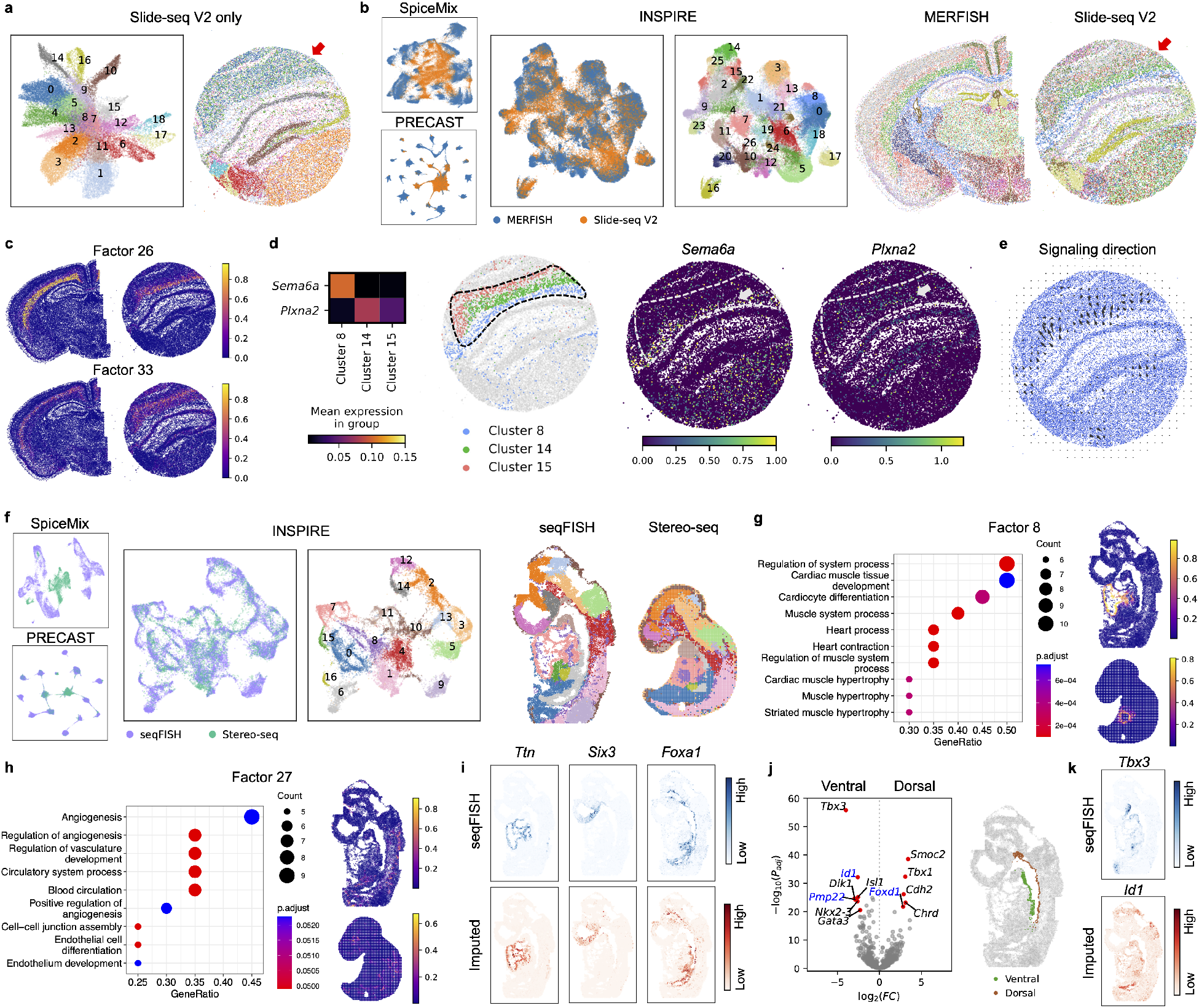
Integrative analyses of data from different ST technologies to enhance biological insights. **a.** Spatial domain identification from the analysis of Slide-seq V2 data alone. **b**. Cell representations aligned between the MERFISH and Slide-seq V2 datasets using SpiceMix, PRECAST, INSPIRE, and spatial domain identification result from INSPIRE. **c**. Spatial distributions of INSPIRE’s factors 26 and 33. **d**. Expressions of genes *Sema6a, Plxna2* among INSPIRE’s identified spatial regions 8, 14 and 15. **e**. Signaling direction from ligand *Sema6a* to receptor *Plxna2* across INSPIRE’s annotated regions inferred by the COMMOT method. **f**. Cell representations aligned between the seqFISH and Stereo-seq embryo datasets using SpiceMix, PRECAST, INSPIRE, and spatial domain identification result from INSPIRE. **g, h**. Significant gene ontology terms and spatial distributions of INSPIRE’s factors 8 (**g**) and 27 (**h**). **i**. We manually held out genes from the seqFISH dataset and imputed their expressions from the Stereo-seq dataset to show INSPIRE’s gene imputation capability. We visualized the expressions of *Ttn, Six3* and *Foxa1* measured by seqFISH (top panel), and compared them with the imputed expression levels (bottom panel). **j**. Differentially expressed gene analysis between the ventral and dorsal sides of the developing gut tube in the seqFISH dataset. Top differentially expressed genes with imputed expression levels from INSPIRE are annotated in blue, and top genes with expression levels quantified by seqFISH are annotated in black. **k**. *Tbx3* expression levels measured by seqFISH, and *Id1* expression levels imputed by INSPIRE.

Next, we focused on analyzing the spatial factors and the associated gene loading in the NMF across slices from INSPIRE. To quantitatively evaluate the quality of spatial factors, we employed two metrics: factor diversity and factor coherence [44]. Factor diversity measures the percentage of unique genes associated with each factor, with a higher diversity score indicating more varied factors. Factor coherence evaluates the interpretability of factors by assessing the co-expression of genes associated with the same factor across spots. A higher coherence score indicates better interpretability of spatial factors. Using these two metrics, we compared the performance of INSPIRE to NMF-based methods SpiceMix and NSFH. As shown by the highest scores for both metrics, INSPIRE achieved superior factor quality compared to other methods (Fig. 2**g**).

Besides quantitative evaluation, we also visualized INSPIRE’s spatial factors to illustrate their ability to decipher detailed spatial organization patterns among slices. For instance, three factors (factors 18, 20, and 17) showed clear enrichment in cortical layers L2, L5, and L6, respectively (Fig. 2**h, i**). By exploring their gene loadings from INSPIRE, we found their correspondence to excitatory neuronal subtypes specific to L2, L5, and L6, respectively. To be specific, we identified marker genes for these neuronal subtypes using an external scRNA-seq atlas (Fig. 2**j**). These marker genes showed top rankings in the gene loadings of these factors (Fig. 2**k**; the Methods section), confirming the biological meanings of the factors. Additionally, with the interpretable gene loadings, INSPIRE is able to unveil variable genes that are associated with the spatial factors, offering the biological insights. For example, genes specific to the three factors were identified (Fig. 2**l**; the Methods section) and their reliability was validated (Fig. 2**m**). Unlike INSPIRE, which uncovered neuronal subtypes in detailed cortical layers, SpiceMix only captured broad layer structures with its factors (Supplementary Figs. 5 and 6). For example, its factor 14 described a mixture of multiple layers, including L4, L5, and L6. Consistent with its relatively low factor quality scores, NSFH’s factors did not present clear spatial structures in the tissue (Supplementary Fig. 7). Notably, beyond cortical layers, INSPIRE also depicted other detailed spatial structures. For instance, INSPIRE revealed the spatial distribution of astrocytes, with the gene signature of factor 12 characterizing the gene expression profile of astrocytes (Fig. 2**j**-**m**). This demonstrated that INSPIRE identified the spatial organization in the tissue that is not covered by manual layer annotation, further highlighting INSPIRE’s ability to decipher fine-grained and interpretable spatial structures using spatial factors and gene loadings.

**Figure 5:**
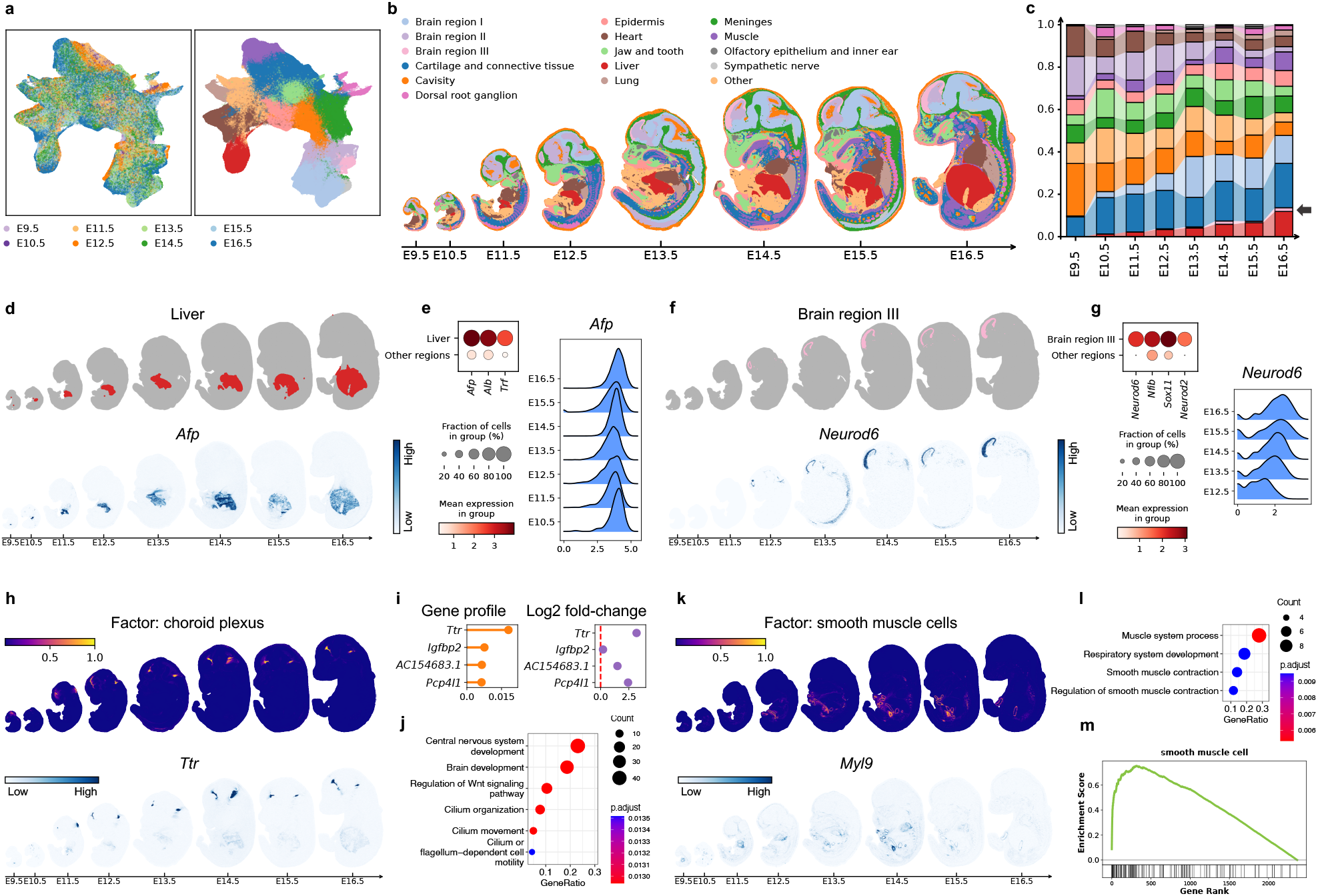
Construction of the spatiotemporal mouse organogenesis atlas by integrating whole-embryo ST slices across various developmental stages. **a.** UMAP plots of spot representations, colored by slice indices and spatial domain labels assigned by INSPIRE. **b**. Visualization of the identified spatial domains across slices. **c**. For each developmental stage, we visualized the proportions of the embryo occupied by different spatial regions. **d, e**. Visualization of the liver region (**d**) and the expression patterns of liver marker genes (**e**). **f, g**. Visualization of a brain region (**f**) and the expression pattern of its differentially expressed genes (**g**). **h, i**. Spatial distribution of the factor representing the choroid plexus (**h**) and its gene signature (**i**). The gene profile indicates the expression levels of genes for this factor. The log2-fold change measures the difference between gene expressions in this factor and other factors. **j**. GO analysis for the genes specific to the factor in **h. k**. Spatial distribution of the factor related to smooth muscle cells and the spatial expression pattern of a smooth muscle cell marker gene. **l**. GO analysis for the genes specific to the factor in **k. m**. Gene set enrichment analysis comparing genes related to the factor in **k** with the marker gene set for smooth muscle cells.

**Figure 6:**
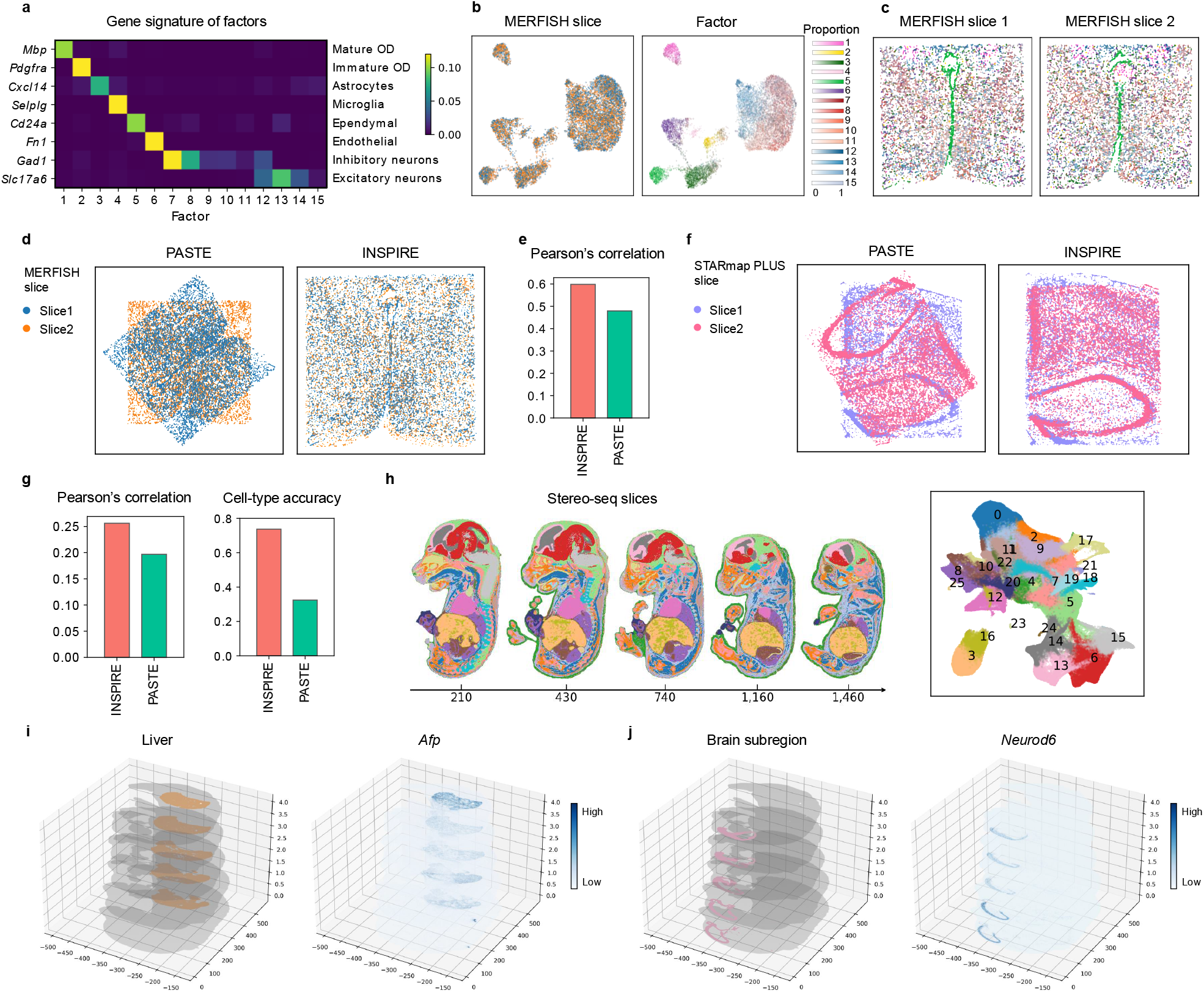
3D reconstruction of tissues and whole organisms by INSPIRE. We utilized INSPIRE to spatially align two MERFISH slices from the mouse hypothalamic preoptic region (**a** - **e**), two STARmap PLUS slices from the mouse hippocampus region (**f, g**), and five whole-embryo slices produced by Stereo-seq (**h** - **j**). **a**. Gene signatures of spatial factors, with each row indicating a marker gene specific to a cell type. **b**. UMAP plots of cell representations from INSPIRE, colored by MERFISH slice indices and the inferred factor proportions among cells. **c**. The MERFISH slices colored by the proportions of factors. **d**. The MERFISH slices after PASTE alignment and INSPIRE alignment, respectively. **e**. Pearson correlation scores of INSPIRE and PASTE evaluated using the MERFISH dataset. **f**. The STARmap PLUS slices after PASTE alignment and INSPIRE alignment, respectively. **g**. Pearson correlation and cell-type accuracy scores of INSPIRE and PASTE evaluated using the STARmap PLUS dataset. **h**. Visualization of the Stereo-seq whole-embryo slices and UMAP plot of spot representations, colored by the identified spatial domains across slices. **i**. Visualization of liver structure and distribution of *Afp* expression on the reconstructed 3D model of the embryo. **j**. Visualization of the 3D structure of a brain subregion and the distribution of *Neurod6* expression on the reconstructed 3D embryo.

### Precise stitching of multiple sagittal and coronal mouse brain slices with partially shared spatial structures

In this section, we evaluate INSPIRE’s performance in a more challenging scenario compared to our previous benchmarking study: integrating multiple slices from different samples, where the spatial structures only partially overlap. This situation presents a unique challenge in learning spot representations and spatial factors, as it requires methods to adaptively identify and align shared biological variations among the slices, while preserving signals unique to each slice and accounting for batch effects.

To explore the complex architecture of the brain, 10x Genomics created Visum slices in both sagittal and coronal planes. Due to the size restriction on the captured area, the sagittal plane was further dissected into two sections, each profiled in an individual ST slice. In total, there are three ST brain slices: the sagittal anterior [30], sagittal posterior [31], and coronal slices [32]. Brain structures captured in these ST slices are only partially shared. For instance, while all the three slices contain the isocortex, the main olfactory bulb is unique to the sagittal anterior slice, and the cerebellum is unique to the sagittal posterior slice. We used INSPIRE to jointly model the three slices, merging data collected from distinct views of the brain.

When applied to this task, INSPIRE effectively depicted the mouse brain architecture. After INSPIRE’s integration, the representations of spatial spots were correctly aligned across the three slices in the latent space (Fig. 3**b**). By clustering the spot representations, INSPIRE was able to partition the mouse brain into 36 distinct and well-organized spatial regions (Fig.3**a**). Among these clusters, six layer-structured spatial domains with labels 2, 6, 7, 8, 25, and 31 together formed the isocortex region shared across the three brain slices (Fig. 3**b, c**). Importantly, INSPIRE also successfully preserved slice-unique tissue structures such as the cerebellum, characterized by spatial regions 3 and 18 in the sagittal posterior slice, and the main olfactory bulb, characterized by spatial regions 20 and 27 in the sagittal anterior slice (Fig. 3**b, c**). In contrast, SpiceMix, NSFH, and PRECAST, which performed relatively well in the human DLPFC-based benchmarking study, produced results that were confounded by strong batch effects and intrinsic differences among the slices (Fig. 3**d** and Supplementary Fig. 12), highlighting their limited applicability in creating a comprehensive tissue ST atlas through the integration of slices with partial overlap. For instance, none of these methods successfully described the shared cortical layers among the multiple mouse brain slices (Fig. 3**d**, Supplementary Figs. 9, 10, and 11).

In this example, we also confirmed that INSPIRE’s designed discriminators (the Methods section) indeed adaptively distinguished between shared biological signals among slices and slice-unique signals, facilitating INSPIRE’s adaptive data integration. In the latent space, discriminators were active for spot populations shared among slices, such as spots in the isocortex (Fig. 3**b, c**), encouraging their alignment across slices. Conversely, discriminators were found inactive for slice-specific spot populations, such as spots in the cerebellum (Fig. 3**b, c**), helping preserve their identities.

Next, we investigated the spatial factors inferred by INSPIRE. Each of them provided a unique and detailed description of the spatial organization in the brain (Supplementary Fig. 13). For instance, the spatial distributions of different hippocampal neuron types in the brain, which are concentrated mainly in the curve-shaped CA1, CA2/CA3, and DG subregions of the hippocampus, were depicted by spatial factors 1, 19, and 39. This observation aligns well with the reference from the Allen Reference Atlas – Mouse Brain [45] (Fig. 3**e**, Supplementary Figs. 16, 17). The gene loadings for spatial factors 1, 19, and 39 also corresponded with marker genes for these hippocampal neuron types identified in an external scRNA-seq atlas [46] (Fig. 3**h**). Using these factors, we were able to identify specific regional markers, such as *C1ql2* and *Fam163b* for CA1; *Cpne7* and *Npy2r* for CA2/CA3; and *Pou3f1* and *Fibcd1* for DG (Fig. 3**e, f**). Furthermore, we showed that INSPIRE excels at capturing detailed spatial organization unique to individual sections by spatial factors. One example is that, non-negative weights (proportions) of spatial factors 15, 17, and 25 on spots revealed spatial distributions of neurons specific to the molecular layer, the Purkinje layer, and the granular layer in the cerebellum, respectively. Using the associated gene signatures, spatially variable genes specific to each of these fine-grained layers were identified, such as *Grm4, Adcy1* for the granular layer; *Ppp1r17, Calb1* for the Purkinje layer; and *Kit, Epha8* for the molecular layer (Fig. 3**j, k**). The spatial specificity of these genes was confirmed by in situ hybridization (ISH) data from the Allen Brain Atlas (Fig. 3**l**), providing additional support for the high quality and interpretability of INSPIRE’s results.

Unlike INSPIRE, both SpiceMix and NSFH produced results that were less satisfactory due to the confounding of their inferred spatial factors by strong batch effects across sections (Supplementary Figs. 14 and 15). Additionally, NSFH showed limited capability in characterizing detailed spatial structures, as evidenced by the lack of clear spatial patterns in its learned factors (Supplementary Fig. 15). This challenging task demonstrates the superior performance and broad applicability of INSPIRE for integrating ST slices with batch effects and a very low degree of spatial overlap, outperforming all other tools.

### INSPIRE integrates ST data from different ST technologies, facilitating multiple downstream analyses

Different ST technologies and platforms produce data with varying spatial resolutions and sequencing depths [3, 4]. In this section, we demonstrate INSPIRE’s performance in integrating data from different ST technologies, leveraging the strengths of each technology to enhance biological insights.

We first collected mouse brain ST datasets from Slide-seq V2 [7] and MERFISH [47]. The Slide-seq V2 dataset contains expression measurements of over 20,000 genes, with a near-cellular spatial resolution of 10 μm, where each barcoded bead typically captures one to two cells. In contrast, the MERFISH dataset measures gene expression levels in individual cells but covers only 1,122 genes. Before integration, we applied the standard Scanpy workflow [48] to each of these datasets. The Slide-seq V2 data failed to recover known layer structure in the isocortex (Fig. 4**a**). Conversely, the MERFISH data successfully revealed this structure, indicating its higher data quality for annotating spatial regions (Supplementary Fig. 18). Aligning these two datasets helps to combine the better spatial organization characterization from MERFISH with the transcriptome-wide measurement from Slide-seq V2. However, prior to integration, the two datasets showed strong discrepancies, making their integration a challenging task (Supplementary Fig. 19).

Despite the challenge, INSPIRE effectively eliminated confounding technical noises in its cell representations, enabling the identification of spatial regions consistent across the two datasets (Fig. 4**b**). For example, it identified three different hippocampal regions (clusters 9, 16, and 23), and three broad cortical layers (clusters 2, 14 and 15). INSPIRE’s spatial factors revealed additional spatial structures at a finer granularity. Specifically, spatial factors 11, 15, 26, 27, 29, and 33 delineated six highly detailed cortical layers in both datasets, unaffected by the strong technical effects (Fig. 4**c** and Supplementary Fig. 23). A notable achievement was INSPIRE’s ability to call out the hidden cortical layer structure within the Slide-seq V2 data by leveraging information from the MERFISH data (Fig. 4**a**-**c**). To demonstrate how this facilitates downstream biological analyses, we utilized the COMMOT method [16] to screen for cell-cell communications among INSPIRE’s identified cortical layers and nearby regions in the Slide-seq V2 data. Such analysis revealed interactions that were undetectable using the MERFISH data alone, due to its limited number of profiled genes. For example, MERFISH did not capture gene *Plxna2*. However, the integrative analysis enabled COMMOT to identify a signaling direction from ligand Sema6a to receptor Plxna2 cross INSPIRE’s annotated regions (Fig. 4**d, e**). Among them, *Sema6a* was enriched in fiber tracts (cluster 8), while *Plxna2* was highly expressed in the inner cortical layer (cluster 14) with decreasing expression in the adjacent cortical layer (cluster 15). This Sema6a-Plxna2 interaction suggests potential cell migration and neuronal connectivity around the inner cortical layer area of the brain [49].

As a comparison, other methods including SpiceMix, NSFH and PRECAST performed less satisfactorily in this challenging task. Their results including representation of cells, identification of spatial domains and/or spatial factors were still severely confounded by the unwanted technical effects (Fig. 4**b**, Supplementary Figs. 20, 21, 22 and 25). Additionally, SpiceMix’s factors did not show clear spatial distributions in any dataset, lacking the ability to uncover spatial organization patterns in this analysis (Supplementary Fig. 24).

Next, we collected two mouse whole-embryo slices generated by seqFISH [33] and Stereo-seq [8] respectively, and applied INSPIRE to jointly analyze them. Similar to previous crosstechnology data example, these datasets exhibited clear discrepancies due to strong technical effects prior to integration (Supplementary Fig. 26). Nevertheless, INSPIRE successfully aligned cell embeddings between the two datasets after data integration (Fig. 4**f**). This alignment facilitated the detection of 16 well-organized and biologically meaningful spatial domains consistent between the two mouse embryos (Fig. 5**f**). For instance, cluster 1 corresponded to developing hearts, while cluster 2 described the midbrains in both embryo datasets. Additionally, INSPIRE’s spatial factors allowed for more detailed spatial characterization of the embryos (Supplementary Fig. 30). For example, spatial factors 6 and 8 accurately delineated subregions within the embryonic hearts, including the atria and ventricles (Fig. 4**g** and Supplementary Fig. 33). Spatial factor 27 was ubiquitously distributed in both embryos. Through gene ontology (GO) analysis of genes related to factor 27, we found it depicted meaningful biological processes such as angiogenesis, vasculature development, and circulatory system processes in both datasets (Fig. 4**h**).

Through this example, we show that INSPIRE’s effective data integrative analysis (Supplementary Fig. 34) enables accurate information transfer across datasets, facilitating deeper biological understanding. To validate this, we manually held out six genes from the seqFISH dataset, which includes only 351 genes in total. These held-out genes included cardiomyocyte markers *Ttn* and *Popdc2* [50, 51], brain markers *Six3* and *Lhx2* [52, 53], and gut endoderm markers *Foxa1* and *Cldn4* [54, 55]. Using INSPIRE, we imputed spatially-resolved expression levels of these genes from the Stereo-seq dataset. The imputed gene expressions showed consistent patterns with the measured expression levels, highlighting INSPIRE’s effective integrative analysis even with a limited number of genes (Fig. 4**i** and Supplementary Fig. 35). After validating INSPIRE’s gene imputation capability, we extended this approach to impute all genes from the Stereo-seq dataset for cells in the seqFISH dataset, increasing the number of genes in the seqFISH dataset from 351 to over 20,000. This comprehensive gene imputation enabled a detailed exploration of gene expression differences between the dorsal and ventral parts of the developing gut tube, which were clearly separated only in the seqFISH dataset (Fig. 4**j**). Based on *t*-test results, the top differentially expressed genes on the dorsal side include *Smoc2, Tbx1, Cdh2, Chrd*, and *Foxd1*, while the top genes enriched on the ventral side include *Tbx3, Id1, Isl1, Dlk1, Pmp22, Nkx2-3*, and *Gata3* (Fig. 4**j**). Dorsal-ventral patterning of the gut tube is essential for the separation of the dorsal esophagus and the ventral trachea in mouse embryos [56, 57]. Consistent with a previous study of mouse embryo [58], genes known to be upregulated in the esophagus, such as *Foxd1*, were detected to be highly expressed on the dorsal side of the gut tube. Similarly, genes upregulated in the trachea, including *Tbx3, Id1*, and *Isl1*, were identified as enriched on the ventral side. This supports the reliability of our analysis of dorsal-ventral differential gene expressions. Notably, among the identified genes, the expression levels of *Id1, Pmp22*, and *Foxd1* were imputed using INSPIRE (Fig. 4**j, k**), underscoring INSPIRE’s ability to facilitate biological discoveries.

To compare the performance of INSPIRE to other methods, we also applied SpiceMix, NSFH, and PRECAST to this cross-technology data analysis. However, consistent with the results from previous cross-technology data example, all these methods had difficulties to account for strong unwanted discrepancies between datasets caused by technical effects, leading to less satisfactory integrative analyses (Fig. 4**f**, Supplementary Figs. 27, 28, 29, 31 and 32). The comparison between INSPIRE and these methods further demonstrated the superior performance and wide applicability of the INSPIRE method.

### INSPIRE creates a comprehensive mouse organogenesis atlas, enabling spatiotemporal analysis of embryonic development

Understanding how a complex organism develops over time and space is a fundamental problem in developmental biology. Recently, mouse whole-embryo slices were collected at eight developmental stages, spanning from embryonic day 9.5 (E9.5) to day 16.5 (E16.5) [8]. To investigate the spatiotemporal dynamics of organogenesis, we applied INSPIRE to jointly analyze this collection of ST slices across different developmental time points.

These eight ST slices were profiled using high-resolution technology Stereo-seq, with each slice containing numerous spatial spots measuring 25 *μm* in diameter. For example, the ST slices sampled at E14.5, E15.5, and E16.5 each included over 100,000 spatial spots. In total, the eight whole-embryo slices encompass more than half a million spatial spots. Analyzing this spatiotemporal dataset presents new challenges for computational methods: it requires not only precise alignment of eight complex slices across various developmental stages, but also demands high efficiency and scalability to manage over half a millions spatial spots.

INSPIRE is scalable to handle this challenging task, and completed the analysis in 80 minutes. It successfully aligned the eight ST slices in its latent space, accommodating the extreme large number of spatial spots and varying embryo sizes (Fig. 5**a**). This alignment enabled INSPIRE to identify biologically meaningful spatial regions that consistently localized across all eight embryonic stages. These regions corresponded to major organs or tissues, such as heart, liver, lung, and brain (Fig. 5**b**). Their spatial locations aligned with the known anatomical structure of the mouse embryo [59]. The identities of these regions were further validated by analyzing the expression levels of specific marker genes, such as *Myl7* for heart, *Afp* for liver, *Sftpc* for lung [8], and *Six3, Lhx2, Otx2*, and *Pou3f1* for brain [52, 60], across all developmental time points (Fig. 5**b, d**, and Supplementary Fig. 36).

Using the well-annotated spatial regions identified by INSPIRE, we visualized the changes in proportions of spatial spots covered by the corresponding organs or tissues within embryos across developmental stages, providing insights into embryo developmental dynamics (Fig. 5**c**). The liver, initially occupying a very small area at the earliest stage E9.5, rapidly increased its proportion within the embryo as development progressed (Fig. 5**c, d, e**). In contrast, the heart was already well-structured at early stages E9.5 and E10.5, with its size remaining relatively stable in subsequent stages (Supplementary Figs. 36 and 37), consistent with its role as the first functional organ to form in the embryo [61]. Importantly, this analysis also highlighted organs or tissues that were barely formed at earlier stages but became well developed in later embryonic stages. For example, the shape of the lung became clear after stage E12.5, aligning with the observed increase in the expression of the lung marker gene after E12.5 (Supplementary Figs. 36 and 38). This finding is consistent with the fact that although lung buds begin to form during the early embryonic stages from E9.5 to E12.5, the process of branching morphogenesis occurs after E12.5 [62]. Similarly, INSPIRE identified a brain region characterized by the specific expression of genes *Neurod6* and *Neurod2*, which only became clearly localized in the forebrain after stage E12.5 (Fig. 5**c, f, g**). The expressions of these two genes suggest active neuronal differentiation and development in the forebrain during these stages. These results demonstrate INSPIRE’s capability to deepen the understanding of developmental dynamics through its effective integrative data analysis.

Using NMF modeling, INSPIRE enabled cross-developmental stage comparisons that went beyond the resolution of major organs or tissues. Instead of concentrating solely on broader structures, it identified intricate tissue architectures and cell type distributions through its spatial factors, allowing for detailed comparisons among embryos at various stages. For instance, two distinct spatial factors emerged in the forebrain region by stage E11.5, gradually forming a layered structure that became clearly visible by stage E14.5 (Supplementary Fig. 39). Similarly, three spatial factors captured the evolving structure of the embryonic mouth and jaw, with their spatial organization becoming increasingly complex by stages E12.5 and E13.5 (Supplementary Fig. 40). Furthermore, analyzing the spatial distributions of INSPIRE’s spatial factors alongside their gene signatures offered more in-depth biological insights into the developmental dynamics.

As an example, we identified a factor annotated as the choroid plexus based on its spatial location in the embryo at E16.5 (Fig. 5**h**). GO analysis of its gene profile, as reported by INSPIRE, confirmed its involvement in central nervous system and brain development. It also linked this factor to biological processes like cilium organization and movement, providing insights into its specific functions (Fig. 5**i, j**). This factor, along with its associated genes *Ttr* and *Pcp4l1*, exhibited clear spatial localization after stage E11.5, illustrating its developmental trajectory over time. Another example is the identification of a factor with detailed spatial concentrations within the embryos, specifically surrounding many tiny tube-like structures (Fig. 5**k**). Gene set enrichment analysis revealed its correspondence with smooth muscle cells (Fig. 5**l, m**), showing alignment between its specific gene programs and a set of smooth muscle cell marker genes such as *Acta2* and *Myl9* [63]. Given that this spatial factor also colocalized with the expression of *Nkx2-3* (Supplementary Fig. 41), we inferred that it described the nuanced distributions of gastrointestinal smooth muscle cells [64]. Spatial visualizations indicated that gastrointestinal smooth muscle cells and tracts became clearly presented after stage E12.5, uncovering their spatiotemporal dynamics. To summarize, the above results highlight the power of INSPIRE to facilitate the study of embryonic development by effectively identifying nuanced differences with biological meanings across developmental stages using spatial factors and the associated gene profiles.

### INSPIRE enables precise 3D reconstruction of tissues and whole organisms

Recently, there has been a growing number of ST datasets composed of multiple parallel 2D slices (x-y-axis) along the z-axis within tissues. Each slice captures a 2D spatial transcriptomic landscape, and when these slices are spatially aligned and jointly analyzed, they offer valuable opportunities to construct a comprehensive 3D view of spatial structures and cell type distributions within tissues or organs, deepening our understanding of complex biological systems. Here, we demonstrate INSPIRE’s capability to reliably register adjacent 2D ST slices, enabling accurate 3D reconstruction of tissues. For a comprehensive illustration, we applied INSPIRE for two different scenarios, utilizing slices that vary in scale from tissue regions to whole organisms.

In the first scenario, we highlighted INSPIRE’s enhanced performance in slice registration by using two sets of mouse brain region slices. Specifically, we first tested INSPIRE on a dataset comprising two adjacent MERFISH slices from the mouse hypothalamic preoptic region [65]. The application of INSPIRE revealed spatial factors that corresponded distinctly to different cell types or subtypes within the hypothalamic preoptic region, aiding in accurate cell-type identification in the brain (Fig. 6**a, b**). Moreover, the two slices displayed consistent spatial factor distributions both in the UMAP plot and in spatial locations, providing valuable information for establishing spatial correspondence between slices (Fig. 6**b, c**). Utilizing this integration result, we identified mutual nearest neighbor (MNN) cells between the two slices within the integrated embedding, serving as anchor pairs for guiding the registration process. These anchors enabled the calculation of the optimal rigid transformation, effectively aligning one slice with the other and achieving precise registration (Fig. 6**d** and Supplementary Fig. 42). However, PASTE, a state-of-the-art method for spatial registration, could not align these two slices as INSPIRE did. To quantitatively compare INSPIRE and PASTE, we utilized the Pearson correlation metric, which measures the similarity of gene expression levels between spatially proximal spots from the two slices after spatial registration, with a higher score indicating better performance. As shown in Fig. 6**e**, INSPIRE demonstrated superior registration performance compared to PASTE. We also compared INSPIRE and PASTE on another dataset, consisting of two STARmap PLUS slices from the mouse hippocampus region [13]. Consistent with the previous results, INSPIRE achieved precise registration between the two slices, while PASTE did not (Fig. 6**f**, Supplementary Figs. 43 and 44). For the quantitative evaluation, in addition to the Pearson correlation metric, we evaluated cell-type accuracy score as the original dataset provided cell type annotations. The cell-type accuracy score measures the similarity of cell type annotation between spatially proximal spots from the two slices after spatial registration, with a higher score indicating better spatial alignment performance. INSPIRE achieved higher scores for both metrics compared to PASTE (Fig. 6**g**), highlighting its superior performance in the slice registration task.

In the second scenario, we tested INSPIRE’s ability to register multiple slices at the whole-organism level. We applied INSPIRE to a Stereo-seq dataset consisting of five adjacent 2D slices taken along the left-right axis of a mouse embryo at developmental stage E16.5. Spatial registration and integrative analysis of these slices are crucial for constructing a comprehensive 3D model of the mouse embryo. In this application, INSPIRE effectively and efficiently integrated 567,381 spatial spots across multiple slices (Supplementary Fig. 45). Its consistent identification of tissues across the slices, such as heart, liver and brain, further showed the reliablity of the integration result (Fig. 6**h**). Guided by MNN cells from each pair of adjacent slices, INSPIRE sequentially aligned all adjacent slices using rigid transformation. The resulting 3D model of mouse embryo successfully reconstructed major organs such as liver and heart, and characterized 3D expression pattern of their marker genes (Fig. 6**i** and Supplementary Fig. 46). INSPIRE was also able to reconstruct detailed spatial structures such as tissue subregions for accelerating comprehensive understandings of tissue 3D organizations. For instance, it described the 3D structures of subregions in the forehead which vary along the left-right axis (Fig. 6**j** and Supplementary Fig. 47).

To summarize, the results from these two scenarios illustrate that INSPIRE could be applied to build 3D architectures of tissues or even the whole organisms with high reliability. This capability of INSPIRE makes it a powerful tool for conducting 3D analyses in a wide range of biological systems, enhancing our understanding beyond traditional 2D analyses.

## Discussion

In this paper, we have presented INSPIRE, an effective and versatile tool powered by advanced deep learning technologies for integrating and interpreting multiple ST slices from diverse sources. The results demonstrate that INSPIRE effectively addresses the challenges posed by the heterogeneity in ST data, such as variations in samples, technologies, and developmental stages. By combining an adversarial learning mechanism and NMF, INSPIRE not only integrated these diverse datasets, but also deciphered fine-grained spatial tissue architectures through spatial factors and interpreted their biological meanings based on their associated gene signatures.

Although several computational methods have also been developed and have greatly facilitated transcriptomics data analysis, direct application of them did not sufficiently address challenges in ST data integrative analysis, as shown in our examples. Methods including Seurat and LIGER, designed to remove batch effects among scRNA-seq datasets, lack the ability to model spatial dependencies among spots or cells in ST data. PASTE and PRECAST, although capable of performing spatially-informed analysis for multiple ST tissue slices, have shown less satisfactory results in managing heterogeneous unwanted variation across ST datasets. PASTE is primarily designed for joint analysis of ST slices from biological replicate samples, whereas PRECAST relies on Gaussian mixture model shared among slices to correct for unwanted variations, which can be less powerful when dealing with strong slice-specific effects. Methods SpiceMix and NSFH use spatially-informed NMF to extract spatial signals from tissue slices, successfully revealing fine-grained spatial organizations with biological interpretations. However, they are limited to handling one slice at a time and are not designed for integrative analysis of ST data. In contrast, INSPIRE addresses all these challenges and offers advantages over existing methods in ST data integrative analysis by the innovations on its model.

The first major advantage of INSPIRE is its utilization of a tailored adversarial learning mechanism. The adversarial learning mechanism adaptively detects unwanted variations among datasets, even when certain slices present their unique biological signals, providing accurate guidance for neural networks to distill meaningful biological variations across slices in the shared latent space. Meanwhile, the nonlinearity of neural networks offers great flexibility for INSPIRE to adjust for heterogeneous unwanted effects originating from diverse sources.

Second, the seamless integration of the adversarial learning mechanism with NMF in INSPIRE can decipher biological signals across multiple slices into detailed and interpretable spatial factors, unconfounded by unwanted variations. The capability to learn NMF consistently among slices is particularly essential to reveal fine-grained spatial structural patterns in multislice integrative analysis. Meanwhile, downstream analyses, including GO analysis and GSEA, provide interpretation of biological meanings of spatial factors, leading to the discovery of spatial cell type distributions and biological processes. Additionally, as shown in the DLPFC example, the learning of spatial factors aids in eliminating redundant signals that are not related to the inferred gene programs, thus enhancing the discovery of biologically meaningful spot representations in the latent space and facilitating improved result in analyses including spatial trajectory inference and spatial domain identification.

Lastly, INSPIRE incorporates GNNs to perform spatially informed analyses. The GNNs take into account the microenvironments of cells or spatial spots within the tissue, enhancing the ability of INSPIRE to understand tissue organizations. Furthermore, with the utilization of lightweight GNNs that allow for mini-batch optimization, INSPIRE is scalable to analyze large-scale ST datasets, as demonstrated by the construction of mouse organogenesis atlas and the 3D reconstruction of mouse embryo tasks, each encompassing over half a million spots.

Through a comprehensive benchmarking study, INSPIRE has shown superior performance in integrating information across multiple tissue slices, significantly enhancing the characterization of spatial architecture compared to existing methods. We also demonstrated INSPIRE’s effectiveness and wide applicability in a range of challenging applications, including stitching together tissue slices with only partially overlapping structures, integrating data from different ST technologies, aligning slices collected at a series of embryonic developmental stages, and reconstructing 3D tissue models. Each of these applications highlighted a distinct advantage of INSPIRE: constructing a comprehensive atlas by merging data from different tissue views, enabling downstream analyses that leverage the strengths of distinct ST technologies, advancing the study of developmental dynamics, and deepening our understanding of 3D tissue organization.

One potential limitation of INSPIRE, despite its numerous strengths, is its dependence on shared genes across datasets for data integration and interpretation. This reliance might result in the exclusion of important gene signals that are unique to specific datasets. Extending this method to incorporate and align non-shared genes among ST datasets could further enhance biological analyses.

The interpretable and scalable integration of diverse ST datasets across different experimental designs is invaluable in advancing biological discoveries. As the field of spatial transcriptomics continues to grow rapidly, the need for comprehensive integrative analysis of ST datasets will only increase. We expect that INSPIRE, with its exceptional performance, interpretability, and versatility, will be a powerful addition to the modern life scientist’s ST analysis toolkit.

## Methods

### The model of INSPIRE

INSPIRE offers interpretable and spatially-informed integration of ST data from multiple tissue slices. Let *s* = 1, 2, *…, S* be the index of ST slices. For slice *s*, we observe gene expression count matrix 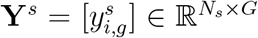 and 2D spatial coordinates of cells or spatial spots 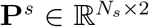, where *i* = 1, 2, *…, N*_*s*_ is the index for cells or spatial spots, and *g* = 1, 2, *…, G* is the index for genes. Using all the information as input, INSPIRE learns to decipher spatial structures across all the *S* slices and infers the associated gene programs.

To integrate both gene expressions and spatial locations across all ST slices, INSPIRE encodes the gene expression information of all cells or spatial spots into a shared latent space *Z*, using a neural network that accounts for spatial dependencies among cells or spots. Specifically, INSPIRE builds a 2D neighborhood graph for each slice using the spatial location matrix. We denote the neighborhood graph for slice *s* as 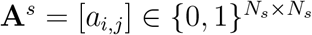, where *a*_*i,j*_ = 1 if cells or spots *i* and *j* are spatial neighbors, and *a*_*i,j*_ = 0 otherwise. Using both gene expressions {**Y**^*s*^}_*s*=1,2,*…,S*_ and spatial graphs {**A**^*s*^}_*s*=1,2,*…,S*_, the latent representations of cells or spatial spots are generated by:

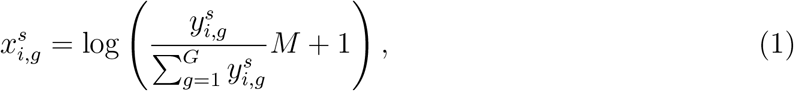

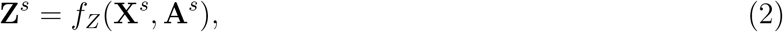

where we perform log-normalization on count matrices {**Y**^*s*^}_*s*=1,2,*…, S*_ for algorithm stability. In the data normalization, we set *M* = 10^4^ for sequencing-based ST data, and set *M* = 10^3^ for imaging-based ST data. The log-normalized data 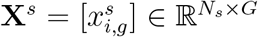 are then encoded to latent representations 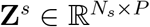 through a graph neural network *f*_*Z*_(·) with parameters shared among all slices, where *P* is the dimensionality of the shared latent space. For slice *s*, the latent representation **Z**^*s*^ embeds information from both gene expressions **Y**^*s*^ and spatial neighbors encoded in graph **A**^*s*^, describing spatially-informed biological variations in slice *s*. Importantly, in addition to capturing spatially-aware biological signals in each slice, the shared latent space is designed for achieving the integration across all input slices. For harmonizing latent representations **Z**^1^, **Z**^2^, *…*, **Z**^*S*^ from different ST slices, INSPIRE adopts a tailored adversarial mechanism in latent space *Ƶ*. Let 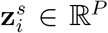 be the latent representation of cell or spot *i* in slice *s*. An auxiliary discriminator network, *D*^*s*^(·) : *Ƶ* → (0, 1), is deployed to identify where the poor mixing between representations 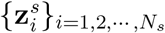 from slice *s* and 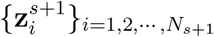 from slice *s* + 1 occurs. Encoder network *f*_*Z*_(·) is trained to compete against discriminator *D*^*s*^(·), aiming to mix 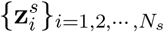 from slice *s* and 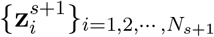 from slice *s* + 1. Through this competition, discriminator *D*^*s*^(·) provides feedback to improve encoder *f*_*Z*_(·) until representations 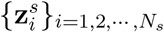 and 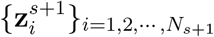 from slices *s* and *s* + 1 are well integrated. INSPIRE introduces *S* − 1 discriminators {*D*^*s*^(·)}_*i*=1,2,*…, S*−1_ for aligning all the *S* tissue slices. Guided by the *S* − 1 discriminators, encoder *f*_*Z*_(·) learns to generate integrated representations of cells or spatial spots across all *S* tissue slices.

Based on the shared latent space, INSPIRE then achieves a harmonized non-negative matrix factorization (NMF) for gene expressions across all input slices, that are not confounded by complex unwanted variations. The hidden spatial factors identified by this integrated NMF across slices provide a unified characterization of fine-grained tissue structures across all slices. Meanwhile, the gene loading matrix describes gene modules associated with each spatial factor, interpreting the biological meanings of the detailed spatial organization patterns discovered by the spatial factors. We assume there are *K* hidden spatial factors in input slices. Each spatial factor characterizes a fine-grained spatial structure in the tissue. Let 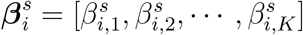 denote the set of non-negative weights among the *K* hidden spatial factors for cell or spot *i* in slice *s*, with 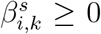 and 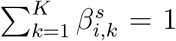. INSPIRE generates 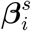 from integrated latent representation 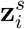 of cells or spots across slices:

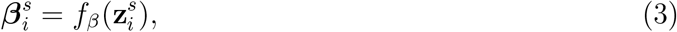

where network *f*_*β*_(·) contains a simple linear layer, followed by a softmax function. Notably, in the shared latent space, representations {**Z**^*s*^}_*s*=1,2,*…, S*_ are spatially-aware and free of unwanted variations. Hence, generated from **Z**^*s*^, the obtained 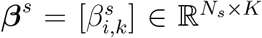 for different slices are also spatially-informed and well integrated across ST slices. The integrated {***β***^*s*^}_*s*=1,2,*…, S*_ together with shared gene loading matrix ***μ*** = [*μ*_*k,g*_] ∈ ℝ_*K×G*_ across slices form a harmonized NMF model for all input slices. In the model, {***β***^*s*^***μ***}_*s*=1,2,*…, S*_ focus on capturing shared biological signal in all slices and further decomposing it into a set of *K* interpretable spatial factors. To account for confounding factors including batch effects and technical effects across ST slices, INSPIRE also introduces slice- and gene-specific effects 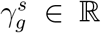 to the integrated NMF model. Combining non-negative weights for spatial factors in cells or spots {***β*** }_*s*=1,2,*…, S*_, non-negative gene loadings ***μ*** shared across slices, and slice- and gene-specific effects 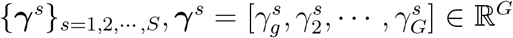 for modeling unwanted variations, INSPIRE reconstructs the observed gene expression counts using the following integrated NMF-based model across all ST slices:

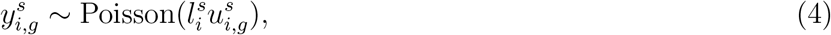

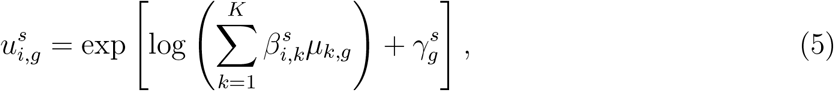

where 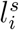 is the observed total transcript count in cell or spot *i* from slice *s*, gene loadings satisfy *μ*_*k,g*_ ≥ 0 and 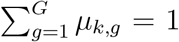. For gene loadings ***μ*** and slice- and gene-specific effects {***γ***^*s*^}_*s*=1,2,*…, S*_, INSPIRE models them as learnable parameters. After training, INSPIRE’s learned ***μ***_*k*_ = [*μ*_*k*,1_, *μ*_*k*,2_, *…, μ*_*k,G*_] ∈ ℝ^*G*^ reveals the gene signature corresponding to hidden spatial factor *k*, where a high value of *μ*_*k,g*_ indicates a greater impact of gene *g* on spatial factor *k*. Through learning and analyzing gene loadings ***μ***, INSPIRE is able to find gene programs that are associated with different hidden spatial factors. Meanwhile, in the integrated NMF across slices, non-negative weights 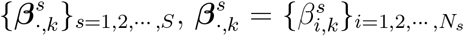 with spatial coordinates of cells or spots describe a spatial enrichment pattern of spatial factor *k* across all the *S* ST slices. Using {***β***^***s***^}_*s*=1,2,*…, S*_, INSPIRE is capable of depicting fine-grained spatial organization structures across all ST slices, without being confounded by unwanted variations.

INSPIRE is a unified method that incorporates the adversarial learning mechanism for data integration with the NMF model for jointly depicting interpretable spatial structures in multiple tissue slices. We propose to train INSPIRE under the following optimization framework:

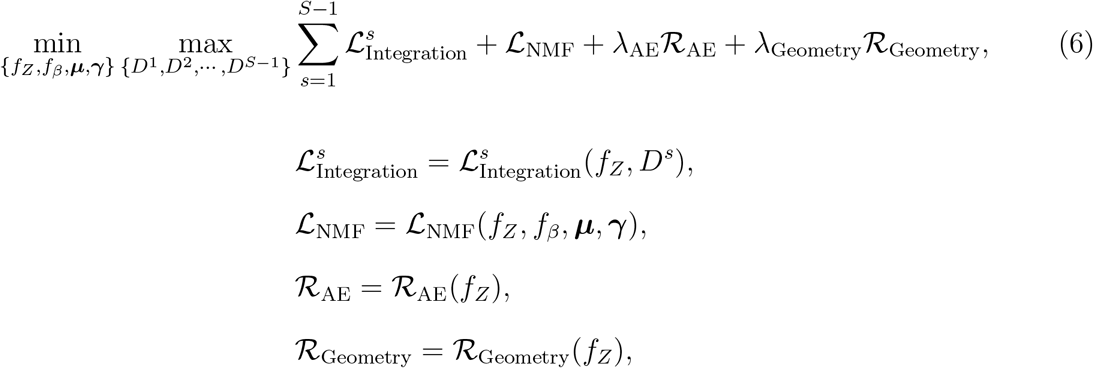

where 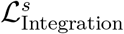 is the objective function of adversarial learning for integrating data from slice *s* and slice *s* + 1; ℒ _NMF_ is the objective function of the joint NMF for reconstructing gene expression counts across all ST slices; ℛ_AE_ and ℛ_Geometry_ are regularizers to encourage the preservation of biological signals across slices in the shared latent space; *λ*_AE_ and *λ*_Geometry_ are coefficients for the two regularizers respectively, and are set to *λ*_AE_ = 1.0 and *λ*_Geometry_ = 0.02. We explain each component in optimization problem (6) in details in the next sections. The parameters in INSPIRE include: parameters in network *f*_*Z*_(·) that encodes latent representations {**Z**^*s*^}_*s*=1,2,*…, S*_; parameters in network *f*_*β*_(·) that generates spatial factors for cells or spots among slices {***β***^*s*^}_*s*=1,2,*…, S*_; gene loading matrix ***μ*** shared across slices; parameters in *S*−1 discriminators {*D*^*s*^(·)}_*s*=1,2,*…, S*−1_ that assist data integration across slices; as well as slice- and gene-specific effects ***γ*** = {***γ***^*s*^}_*s*=1,2,*…, S*_ that account for unwanted variations. After training, INSPIRE simultaneously outputs latent representations {**Z**^*s*^}_*s*=1,2,*…, S*_, spatial factors {***β***^*s*^}_*s*=1,2,*…, S*_ and gene loadings ***μ***. The latent representations of cells or spatial spots are utilized for identifying major spatial domains in tissues and detecting spatial trajectories. Detailed spatial factors {***β***^*s*^}_*s*=1,2,*…, S*_ are used for the discovery of fine-grained tissue sub-regions and spatial distributions of cell types, providing a characterization of spatial patterns in tissues at a higher resolution. Gene loading matrix ***μ*** characterizes gene signatures associated with the detailed spatial structures discovered by spatial factors. It deciphers the biological meaning of spatial factors through factor-specific gene program detection and pathway enrichment analysis.

### Adversarial learning mechanism for data integration across slices

The adversarial training between the discriminators and the encoder is formulated as a min-max optimization problem, 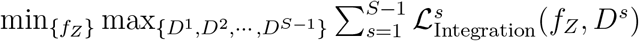, contained in (6), where

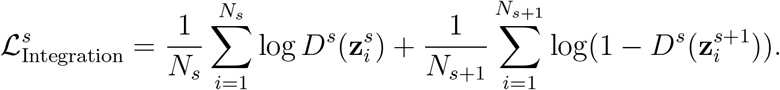

The latent representations are obtained using Eqs. (1) and (2). Given latent codes 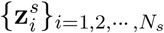 and 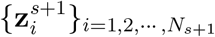 generated by *f*_*Z*_(·), discriminator *D*^*s*^(·) : *Ƶ* → (0, 1) is trained to distinguish between 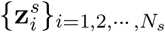 from slice *s* and 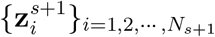 from slice *s* + 1. Here, *D*^*s*^(·) is trained to output a high score (close to one) for representations in slice *s*, while it learns to assign a low score (close to zero) for representations in slice *s* + 1. This is achieved by maximizing 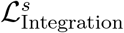 with respect to *D*^*s*^(·). Given discriminators {*D*^*s*^(·)}_*s*=1,2,*…, S*−1_, encoder *f*_*Z*_(·) is trained to mix latent representations across all slices, such that any discriminator cannot distinguish latent codes between slices. This is achieved by minimizing 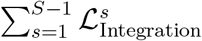 with respect to *f*_*Z*_(·). Through the competition between encoder *f*_*Z*_(·) and discriminators {*D*^*s*^(·)}_*s*=1,2,*…, S*−1_, the discriminators will guide the improvement of the encoder until the encoder generates integrated latent representations for cells or spatial spots across all the *S* slices.

With the above design, Discriminator *D*^*s*^(·) will guide *f*_*Z*_(·) to mix 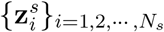 from slice *s* with 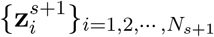 from slice *s* + 1. However, a slice-specific cell or spot population should not be mixed with cells or spots from another slice. To preserve slice-specific cell populations from being incorrectly mixed with other cells, we follow our previous work [34] to adopt a thresholding for discriminator scores. Consider a cell population that is unique in slice *s*. Discriminator *D*^*s*^(·) can easily recognize cells in this population as cells from slice *s*, and assign extremely high scores to them. By similar reasoning, *D*^*s*^(·) will assign extremely low scores to slice *s* + 1-unique cell populations. Therefore, as slice-unique cell populations are prone to be assigned with extreme discriminator scores, we set boundaries for discriminator scores to make discriminators inactive on them. Specifically, the outputs of standard discriminators are transformed into (0, 1) through the sigmoid function. For any applicable latent code **z**, *D*^*s*^(**z**) = sigmoid(*d*^*s*^(**z**)) = 1*/*(1 + exp(−*d*^*s*^(**z**))), where *d*^*s*^(**z**) ∈ R is the logit value of discriminator score *D*^*s*^(**z**). We bound discriminator score *D*^*s*^(**z**) by thresholding its logit *d*^*s*^(**z**) to a reasonable range [−*m, m*], where *m* is set to 50.0:

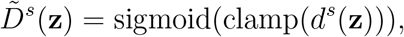

where clamp(·) = max(min(·, *m*), −*m*), *m >* 0. By clamping *d*^*s*^(**z**), 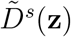 becomes a constant when *d*^*s*^(**z**) *<* −*m* or *d*^*s*^(**z**) *> m*, providing zero gradients for updating parameters in encoder network *f*_*Z*_(·). With this design, the slice-unique cell populations with extreme *d*^*s*^(**z**) scores will be left in the inactive region of discriminators. Consequently, discriminators will not force encoder *f*_*Z*_(·) to mix slice-unique cell populations with other cells, avoiding incorrect integration. Meanwhile, 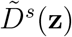 remains the same as *D*^*s*^(**z**) when 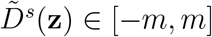, effectively guiding encoder *f*_*Z*_(·) to align cells that are likely to belong to the shared cell populations among slices for data integration. For clarity, we still use the notation *D*^*s*^(·) to denote discriminator 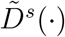 with the score thresholding design hereinafter.

### Joint NMF for reconstructing gene expressions in multiple ST slices

The major objective function in optimization problem (6) of INSPIRE, ℒ_NMF_, corresponds to the reconstruction of gene expression counts in all *S* input slices through a joint NMF model. INSPIRE uses encoder *f*_*Z*_(·) to generate integrated data {**Z**^*s*^}_*s*=1,2,*…, S*_ across slices, guided by discriminators {*D*^*s*^(·)}_*s*=1,2,*…, S*−1_. By Eq. (3), it then leverages network *f*_*β*_(·) to decompose the signal captured in {**Z**^*s*^}_*s*=1,2,*…, S*_ into a set of *K* interpretable spatial factors 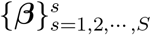 that characterize spatial structures of tissues at a fine-grained level. Combining the obtained spatial factors with addtional parameters, including gene loadings ***μ*** as well as slice- and gene-specific effects ***γ***, INSPIRE reconstructs the gene expression counts from all inputs slices through an integrated NMF-based model described by Eqs. (4) and (5). Based on this model, the corresponding objective function is given by

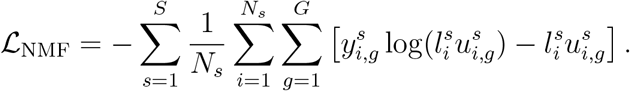

By minimizing ℒ_NMF_ with respect to *f*_*Z*_(·), *f*_*β*_(·), ***μ*** and ***γ***, INSPIRE deciphers interpretable hidden spatial factors that are unified across slices with {***β***^*s*^ ***μ***}. Here, {***β***^*s*^}_*s*=1,2,*…, S*_ characterizes detailed spatial organizations in tissues, while ***μ*** describes gene signatures associated with the tissue organization patterns identified by spatial factors for interpretability.

### Regularization for encouraging the preservation of biological variations across slices

INSPIRE uses two regularizers, ℛ_AE_ and ℛ_Geometry_, to help preserve biological signals across slices in the shared latent space. We design regularizer ℛ_AE_ as

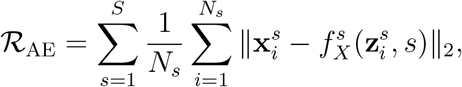

where slice-specific neural network 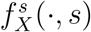 is introduced to reconstruct log-normalized gene expressions in cells or spots 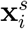 based on latent codes 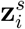 and slice label *s*. Encoder *f*_*Z*_(·) and network 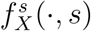 together form an auto-encoder structure between log-normalized data {**X**^*s*^}_*s*=1,2,*…, S*_ and latent codes {**Z**^*s*^}_*s*=1,2,*…, S*_. In regularizer ℛ_AE_, slice-specific network 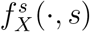 is designed to recover gene expressions from the latent space while accounting for slice-specific effects using slice labels. Therefore, encoder *f*_*Z*_(·) is encouraged to distill all biological information into the latent space without slice-specific effects. To preserve a good geometric structure in the latent space for revealing biological signals, e.g., continued developmental trajectories of cells, we propose regularizer ℛ_Geometry_:

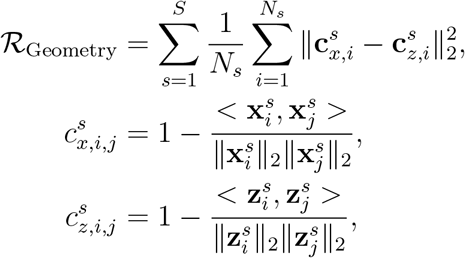

where for cells or spots *i* and *j* from slice *s*, 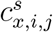 is the cosine similarity between their log-normalized gene expressions 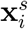 and 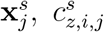 is the cosine similarity between their latent representations 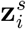 and 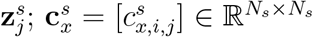 and 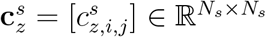 are corresponding cosine similarity matrices; 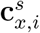 and 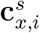 are the *i*-th rows of 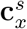 and 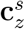 respectively. In regularizer ℛ_Geometry_, cells or spots with high similarities in gene expressions are encouraged to be close in the latent space, while cells or spots with dissimilar gene expressions are encouraged to remain distinct in the latent space. Hence, by using ℛ_Geometry_, INSPIRE encourages *f*_*Z*_(·) to preserve biological meaningful structures in the shared latent space.

### Selection of informative genes

When all input ST slices provide the whole transcriptome profiling, INSPIRE selects the informative genes and uses them as features. Following the Scanpy pipeline [48], INSPIRE selects the top *M* highly variable genes for each slice. It then takes the intersection of these highly variable genes across all ST slices to ensure that the features of cells or spatial spots are shared across all slices. By default, we set *M* = 6, 000. If the number of selected features is less than 2, 000 with *M* = 6, 000, a larger value of *M* can be adopted. When some ST slices to be analyzed are based on ST technologies that measure the expressions of a limited number of genes, such as MERFISH, INSPIRE uses all the shared genes across the input slices for the integrative analysis.

### Network structures

Encoder *f*_*Z*_(·) contains a graph neural network (GNN) layer and a dense layer. The GNN layer takes log-normalized gene expressions 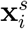 and spatial graph **A**^*s*^ as input. It outputs 512-dimensional hidden vectors. Then the dense layer in *f*_*Z*_(*·*) maps the 512-dimensional hidden vectors to the *P* -dimensional latent representations 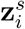 of cells or spatial spots. We set *P* = 32 throughout all analyses. Inspired by previous works [15, 66, 67], INSPIRE provides two options for the GNN layer: the graph attention layer and the lightweight graph-convolutional layer.

The graph attention layer is formulated as:

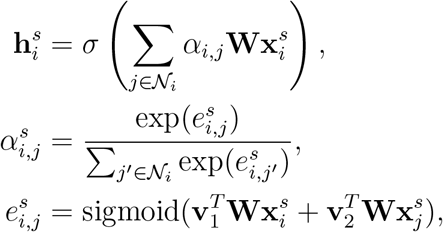

where 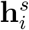 represents the output of the graph attention layer; 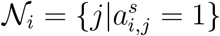 represents the neighbor set of cell or spot *i* encoded in spatial graph **A**^*s*^; **W, v**_1_ and **v**_2_ are parameters in the graph attention layer; *σ*(·) is the activation function. Based on the graph attention mechanism, parameters **v**_1_ and **v**_2_ are used to learn edge weights 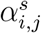 between neighboring cells or spots, helping to adaptively borrow information from neighboring cells or spots. INSPIRE adopts the graph attention layer in *f*_*Z*_(·) when handling the integration task for ST datasets with moderate numbers of cells or spatial spots. To integrate large atlas-scale ST datasets which contain hundreds of thousands or even millions of cells or spots, INSPIRE uses the lightweight graph-convolutional layer in *f*_*Z*_(·), which is formulated as:

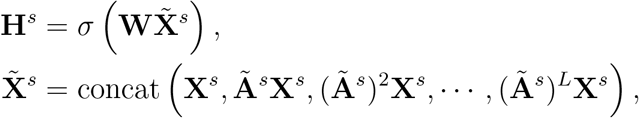

where **H**^*s*^ represents the output of the lightweight graph-convolutional layer; **W** denotes the parameters; 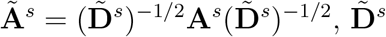 is the diagonal degree matrix of **A**^*s*^; and *L* is the number of steps in the concatenation. We set *L* = 1 by default. The graph attention layer has the advantage of providing an inference of the edge importance between neighborhood cells or spots for adaptively aggregating information in microenviroments of cells or spots. By preparing 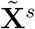 as a preprocessing step, the lightweight graph-convolutional layer enables an efficient training with mini-batch samples from 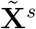, serving as a scalable approach to account for spatial dependencies in datasets with large numbers of cells or spots.

For network *f*_*β*_(·) which produces *K*-dimensional spatial factors 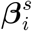^*s*^ from latent codes 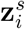 of cells or spatial spots, INSPIRE adopts a one-layer dense network with the softmax activation. The choice of *K* depends on the scale of the tissues to be analyzed. For example, INSPIRE set *K* = 20 for analyzing the cortex region of the brain, while it adopts *K* = 40 for analyzing the whole-brain slices.

For discriminators {*D*^*s*^(·)}_*s*=1,2,*…, S*−1_ that guide encoder *f*_*Z*_(·) to achieve the data integration, INSPIRE uses two-layer dense networks. *D*^*s*^(·) works as a binary classifier to distinguish between 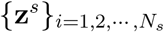 from slice *s* and 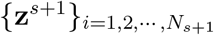 from slice *s*. Then, encoder *f*_*Z*_(·) competes against *D*^*s*^(·) to integrate 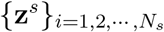 with 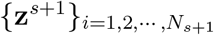. Here, *D*^*s*^(*·*) takes latent codes **z** of cells or spots from slices *s* or *s* + 1 as input. It first uses a dense layer with an activation function to map **z** to a 512-dimensional hidden state. It then uses another dense layer to produce a score belonging to (0, 1) from the 512-dimensional hidden state.

INSPIRE introduces slice-specific network 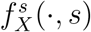 in its regularizer ℛ_AE_ to help preserve biological variations across slices in the shared latent space. 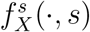 takes latent representations 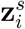 and slice label *s* as inputs. It adopts a graph attention layer or a lightweight graph-convolutional layer, followed by a dense layer, to recover log-normalized gene expressions 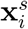 while accounting for slice-specific effects and spatial dependencies among cells or spatial spots. The dimensionality of the hidden state in 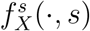 is set to be 512.

### Model training details

INSPIRE employs Adamax, which is a variant of the Adam algorithm [68], for stochastic optimization during model training. By default, the number of optimization steps in INSPIRE is set to 10,000 with learning rate lr = 0.0005, coefficients for computing running averages *β*_1_ = 0.9, *β*_2_ = 0.999 and weight decay parameter *λ* = 0.0001. We conducted all experiments on a single graphics processing unit. The computation times for all experiments are detailed in Supplementary Table 1.

### Evaluation metrics

We evaluated spot or cell representations using ASW and assessed spatial domain identification results, inferred from these representations, with ARI and NMI metrics. The quality of spatial factors was measured by factor diversity and factor coherence.

*ASW*. ASW calculates the silhouette width of cells or spatial spots with respect to spatial region annotation labels. A higher score indicates that cells or spots within the same spatial region are closely grouped, while those from different spatial regions are well separated.

*ARI*. ARI measures the alignment between spatial domain identification result and expert manual annotation. A lower score suggests that the two sets of labels for cells or spots are independent, while a higher score indicates that the labels are identical, except for a possible permutation.

*NMI*. NMI calculates the normalized mutual information between spatial domain identification result and expert manual annotation. A low NMI value indicates minimal shared information between the label sets, while a high value suggests a strong correlation between them.

*Factor diversity*. Topic diversity is defined as the percentage of unique genes among the top 10 genes across all factors. A higher score reflects greater diversity among factors, while a lower score indicates more redundancy.

*Factor coherence*. Topic coherence measures the interpretability of factors by calculating the average pointwise mutual information of top genes associated with each factor, then averaging these values across all factors. Specifically,

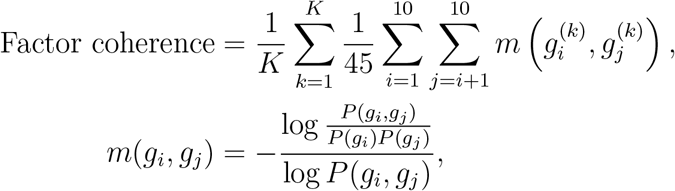

where 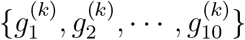 denotes the top 10 genes associated with factor *k, m*(·, *·*) is the normalized pointwise mutual information, *P* (*g*_*i*_, *g*_*j*_) is the probability of genes *g*_*i*_ and *g*_*j*_ coexpressing in a spot or cell, and *P* (*g*_*i*_) is the marginal probability of gene *g*_*i*_.

### Rankings of genes within a spatial factor

The gene loading associated with spatial factor *k* is represented as ***μ***_*k,·*_ = [*μ*_*k*,1_, *μ*_*k*,2_, *…, μ*_*k,G*_], where ***μ*** is the gene loading matrix derived from integrated NMF across all input slices, and *G* is the total number of genes analyzed. The non-negative values in ***μ***_*k,·*_ indicate the relative expression levels among genes within a factor, with a higher *μ*_*k,g*_ corresponding to greater enrichment of gene *g* expressioin in factor *k*. Therefore, for each spatial factor *k*, we can rank genes according to the non-negative values in ***μ***_*k,·*_. The gene with the highest value is ranked first, and the gene with the lowest value is ranked last.

### Identification of genes specific to a spatial factor

To identify highly expressed genes specific to spatial factor *k*, we first select the top *G*_0_ ranked genes based on the gene loading associated with spatial factor *k*, with *G*_0_ set to 50 by default. For each selected gene *g*, we then calculate the fold change between its estimated weight on spatial factor *k* and all other spatial factors. A gene *g* is considered highly expressed and specific to spatial factor *k*, if 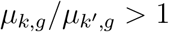 for any *k*^*′*^ ≠ *k*.

### Number of spatial factors

In the human DLPFC example, we demonstrated that the NMF component in INSPIRE enhances the accuracy of spot representations in the latent space. Additionally, we investigated how varying the number of spatial factors in INSPIRE affects the quality of these spot representations. The results indicate that the quality remained stable across different numbers of spatial factors, with a slight improvement in accuracy as the number of spatial factors increased (Supplementary Fig. 48 and Supplementary Note 1).

Next, we examined the relationship between the number of spatial factors and both the quality of the spatial factors and the model fitting accuracy using the human DLPFC and mouse brain data. The results suggest that increasing the number of spatial factors improves model fit to the ST datasets but also reduces the diversity of the spatial factors. These examples illustrate that the optimal number of spatial factors depends on the scale of the ST slices. Specifically, for the human DLPFC slices, representing only a subregion of the brain, the factor diversity score dropped below 50% when the number of spatial factors exceeded 20. In contrast, for the mouse brain data, which includes multiple complementary views of the brain, the factor diversity score remained above 50% even with 40 spatial factors (Supplementary Figs. 49, 50, and Supplementary Note 2). Based on empirical observations, we recommend using number of spatial factors *K* = 20 for analyzing a subregion of an organ, *K* = 40 for a complex organ, and *K* = 60 for a whole organism. Alternatively, the INSPIRE model can be run with different values of spatial factor number *K*, and we recommend selecting the largest *K* such that factor diversity exceeds a specified threshold. By default, we suggest a threshold of 50%.

## Data availability

All data used in this work are publicly available through online sources.

- Human dorsolateral prefrontal cortex dataset profiled by Visium platform [40] (https://research.libd.org/spatialLIBD/).
- Mouse brain sagittal anterior, sagittal posterior, and coronal sections profiled by Visium [30, 31, 32] (https://www.10xgenomics.com/datasets).
- Mouse brain slice profiled by Slide-seq V2 [7] (https://singlecell.broadinstitute.org/single_cell).
- Mouse brain slice profiled by MERFISH [47] (https://doi.brainimagelibrary.org/doi/10.35077/act-bag).
- Mouse whole-embryo slice profiled by seqFISH [33] (https://crukci.shinyapps.io/SpatialMouseAtlas/).
- Mouse whole-embryo datasets across different developmental time points profiled by Stereo-seq [8] (https://db.cngb.org/stomics/mosta/).
- Mouse hypothalamic preoptic region slices profiled by MERFISH [65] (https://doi.org/10.5061/dryad.8t8s248).
- Mouse hippocampus region slices profiled by SRARmap PLUS [13] (https://doi.org/10.5281/zenodo.7458952).

## Code availability

The INSPIRE software is available at https://github.com/jiazhao97/INSPIRE.

## Acknowledgements

This work was supported in part by NIH grants R01 GM134005, U24 HG 012108, and P50 CA196530 to H.Z.; NCCIH grant R01 AT012041, the Allen Discovery Center program, a Paul G. Allen Frontiers Group advised program of the Paul G. Allen Family Foundation to R.B.C.

