## Supplementary Material for "INSPIRE: interpretable, flexible and spatially-aware integration of multiple spatial transcriptomics datasets from diverse sources"

#### Contents

**Supplementary Figures** **2**

**Supplementary Tables** **33**

**Supplementary Notes** **34**

### Supplementary Figures

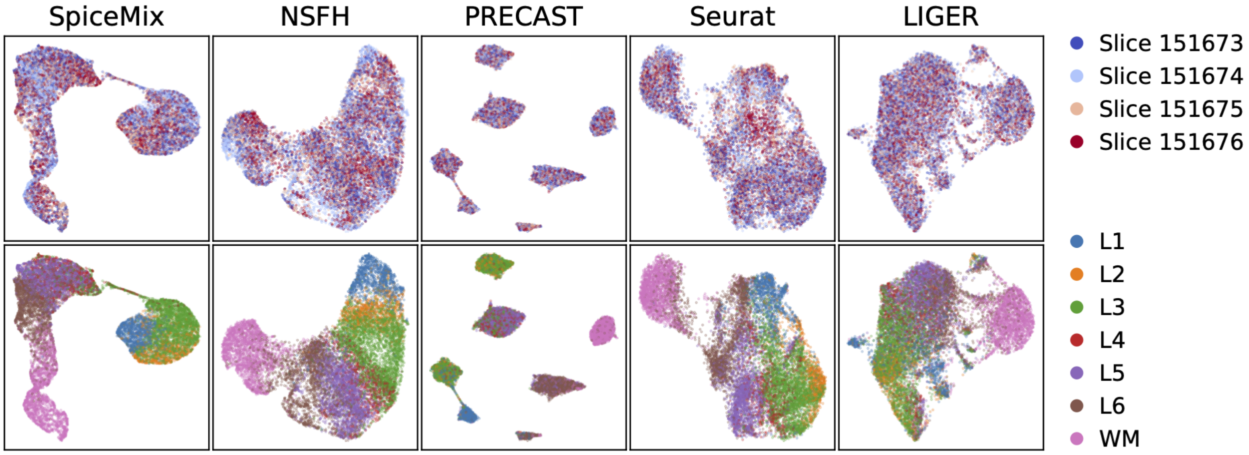


Supplementary Fig. 1: **Comparison of spot representations among methods in the analysis of the human dorsolateral prefrontal cortex (DLPFC) dataset [1].** We visualized spot representations obtained from the compared methods with UMAP plots, colored by slice indices and manual annotations.

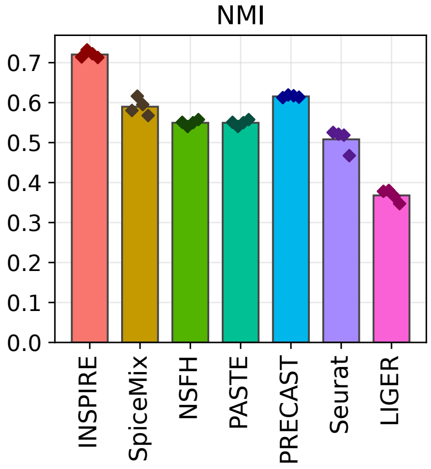


Supplementary Fig. 2: **NMI scores of methods evaluated based on the human DLPFC dataset.** We used the NMI score to evaluate the reliability of spatial region identification result by comparing it to the manual annotation. A higher NMI score indicates a better performance.

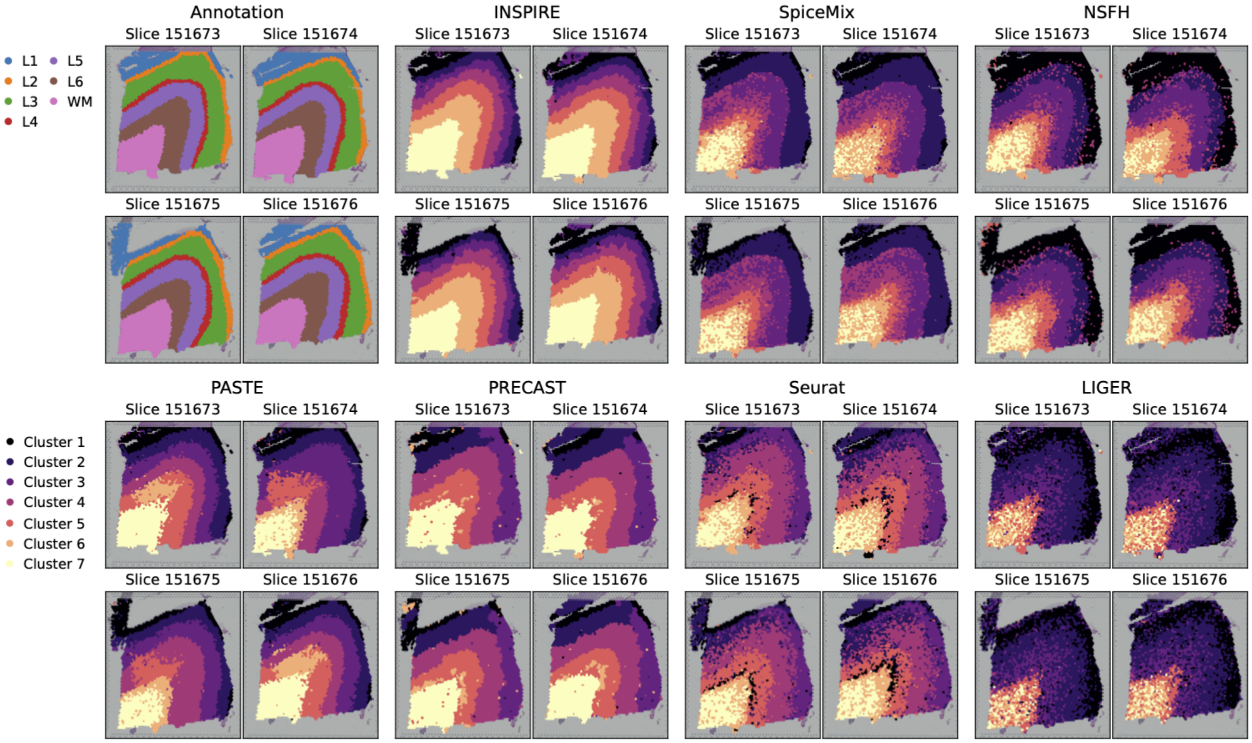


Supplementary Fig. 3: **Comparison of spatial domain identification methods based on the human DLPFC dataset.** We visualized spatial domain detection results of all compared methods, and ground truth layer annotation.

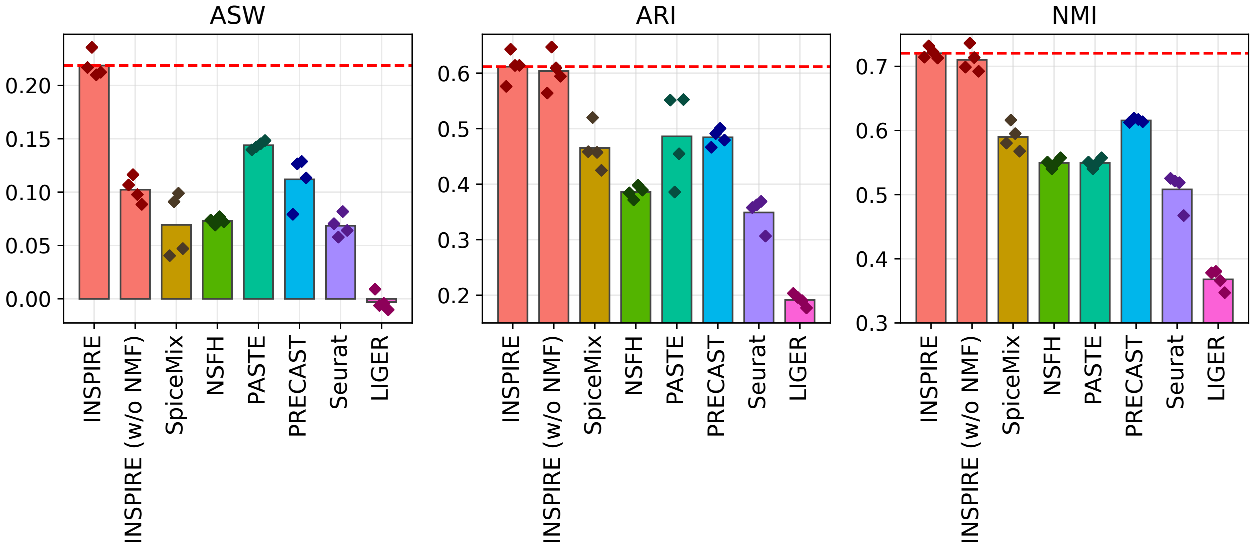


Supplementary Fig. 4: **Comparison of ASW, ARI and NMI scores among INSPIRE, INSPIRE (w/o NMF) and other state-of-the-art methods.** We manually removed the NMF component from INSPIRE, and denoted this version as “INSPIRE (w/o NMF)”. The red dashed lines correspond to the scores of INSPIRE.

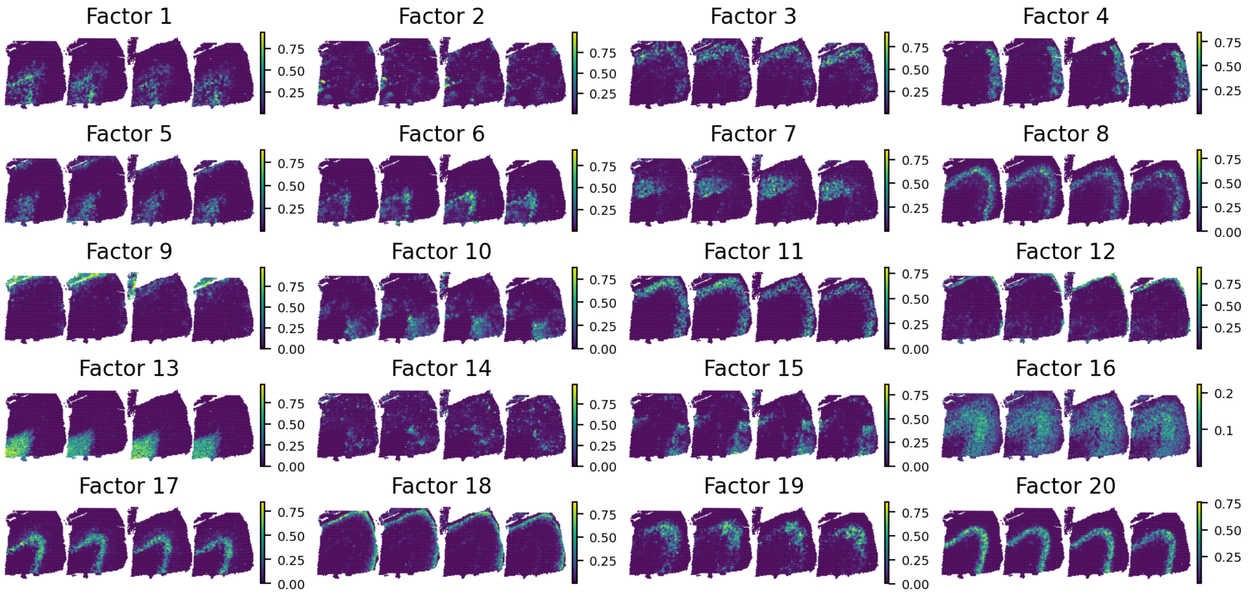


Supplementary Fig. 5: **INSPIRE's identified spatial factors in the DLPFC dataset.** The spatial distribution of each spatial factor is visualized on four slices, with slices indexed 151673-151676 from left to right, respectively.

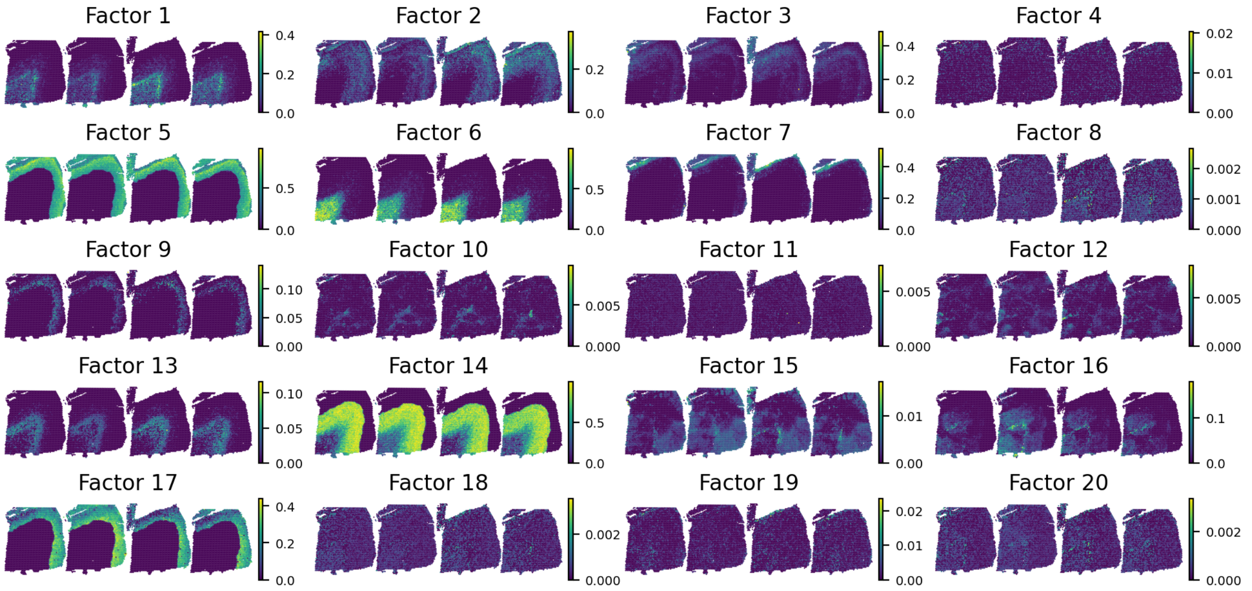


Supplementary Fig. 6: **SpiceMix's identified spatial factors in the DLPFC dataset.** The spatial distribution of each spatial factor is visualized on four slices, with slices indexed 151673-151676 from left to right, respectively.

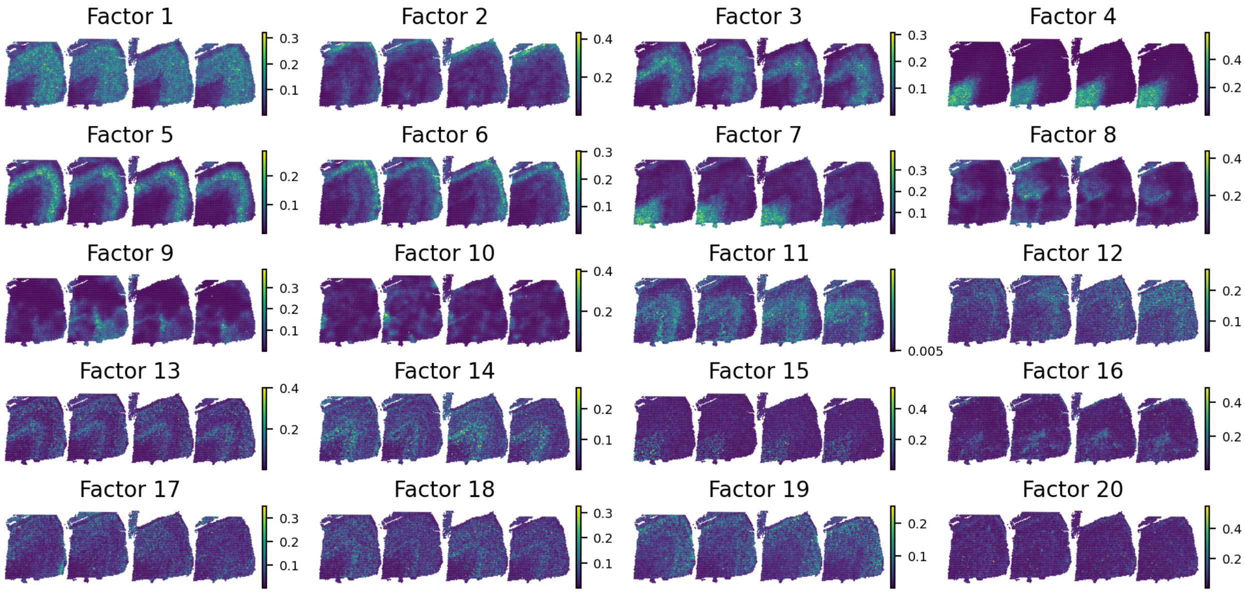


Supplementary Fig. 7: **NSFH's identified spatial factors in the DLPFC dataset.** The spatial distribution of each spatial factor is visualized on four slices, with slices indexed 151673-151676 from left to right, respectively.

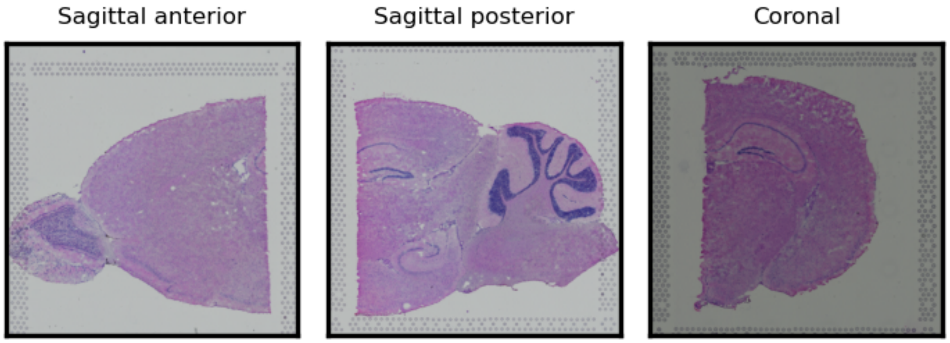


Supplementary Fig. 8: **H&E-stained histological images of the sagittal anterior, sagittal posterior and coronal slices of mouse brains.**

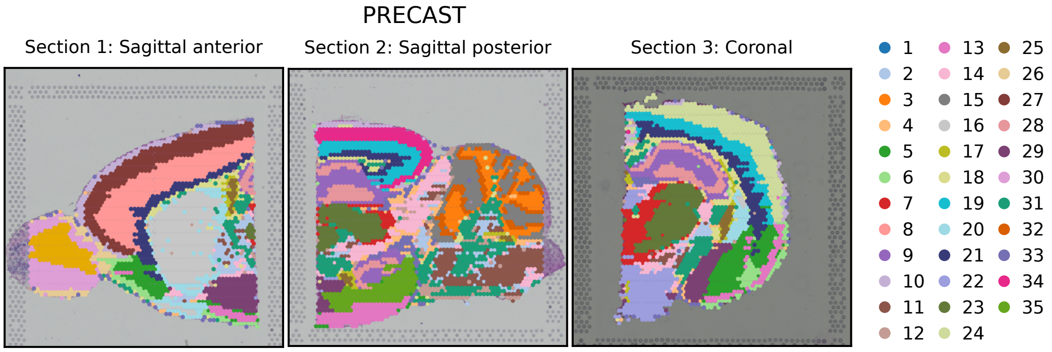


Supplementary Fig. 9: **Spatial domain identification result from PRECAST.** We applied PRECAST to integrate the sagittal anterior, sagittal posterior, and coronal slices from mouse brains. The integrated representations of spots from PRECAST were used to perform the spatial region identification task. We visualized PRECAST's identified spatial domains on H&E-stained histological images of the three slices, respectively.

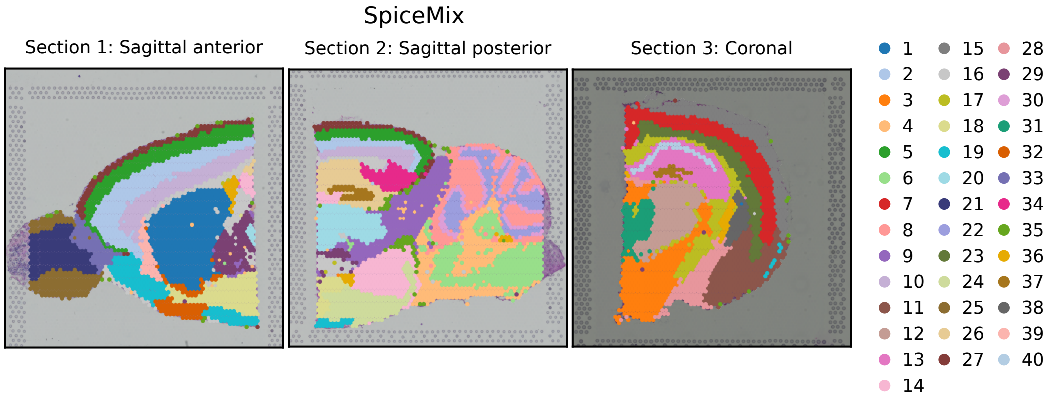


Supplementary Fig. 10: **Spatial domain identification result from SpiceMix, visualized on H&E-stained histological images of mouse brain sagittal anterior, sagittal posterior, and coronal slices, respectively.**

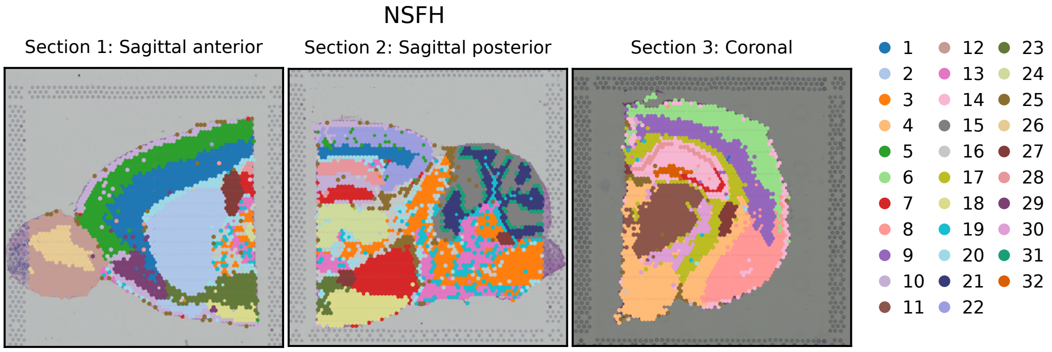


Supplementary Fig. 11: **Spatial domain identification result from NSFH, visualized on H&E-stained histological images of mouse brain sagittal anterior, sagittal posterior, and coronal slices, respectively.**

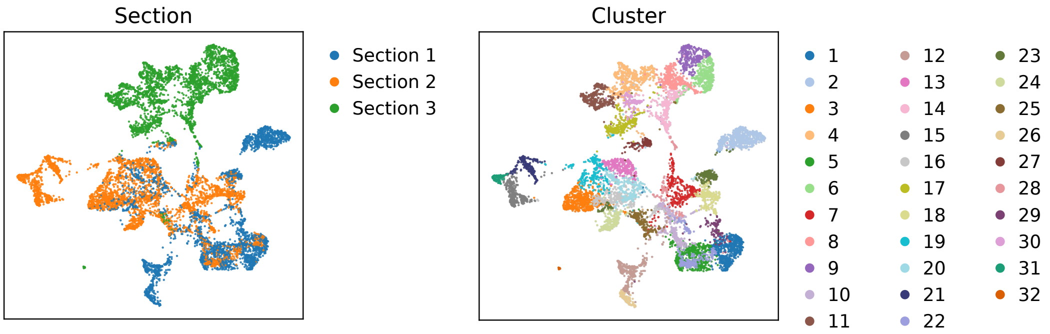


Supplementary Fig. 12: **UMAP plots of spot representations inferred by NSFH.** The UMAP plots are colored by ST section indices and NSFH's identified spatial domain labels. Sections 1, 2, 3 correspond to the sagittal anterior slice, the sagittal posterior slice, and the coronal slices of mouse brains, respectively.

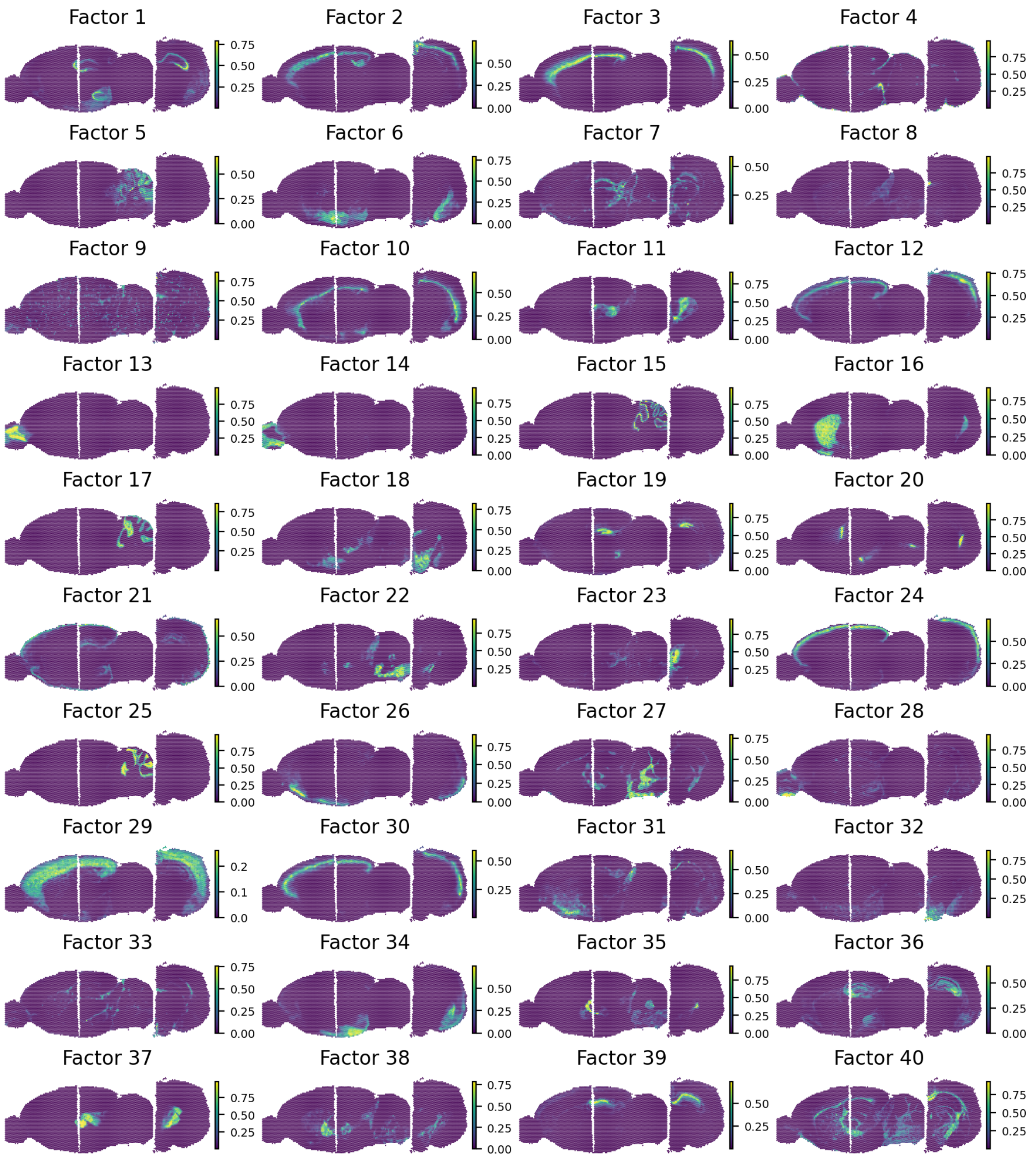


Supplementary Fig. 13: **INSPIRE's identified spatial factors in the mouse brain dataset.** The spatial distribution of each factor is visualized on three mouse brain slices: sagittal anterior, sagittal posterior, and coronal slices, displayed from left to right, respectively.

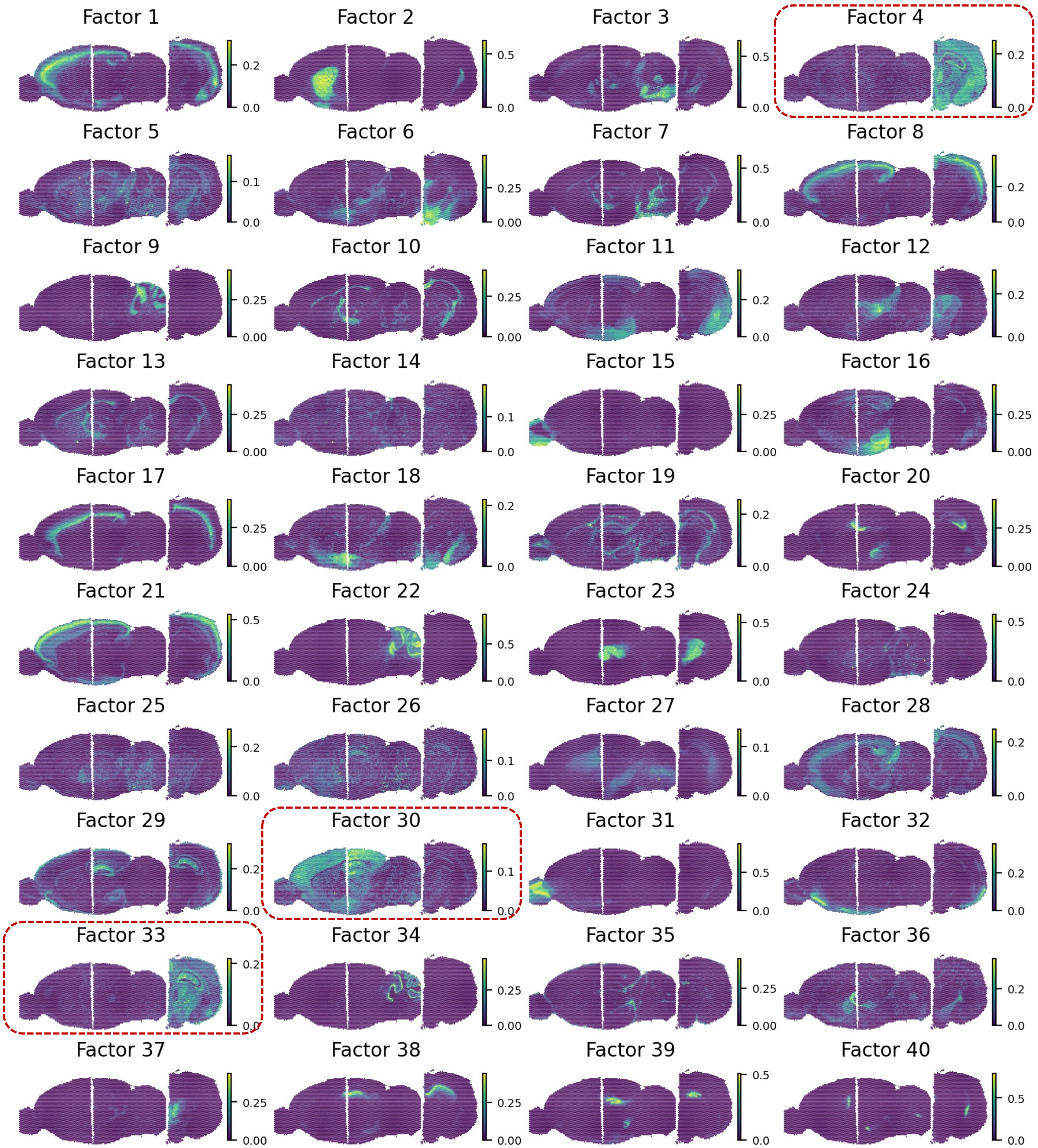


Supplementary Fig. 14: **SpiceMix's identified spatial factors in the mouse brain dataset.** The spatial distribution of each factor is visualized on three mouse brain slices: sagittal anterior, sagittal posterior, and coronal slices, displayed from left to right, respectively. We marked SpiceMix's factors that are affected by batch effects among slices, including factors 4, 30 and 33.

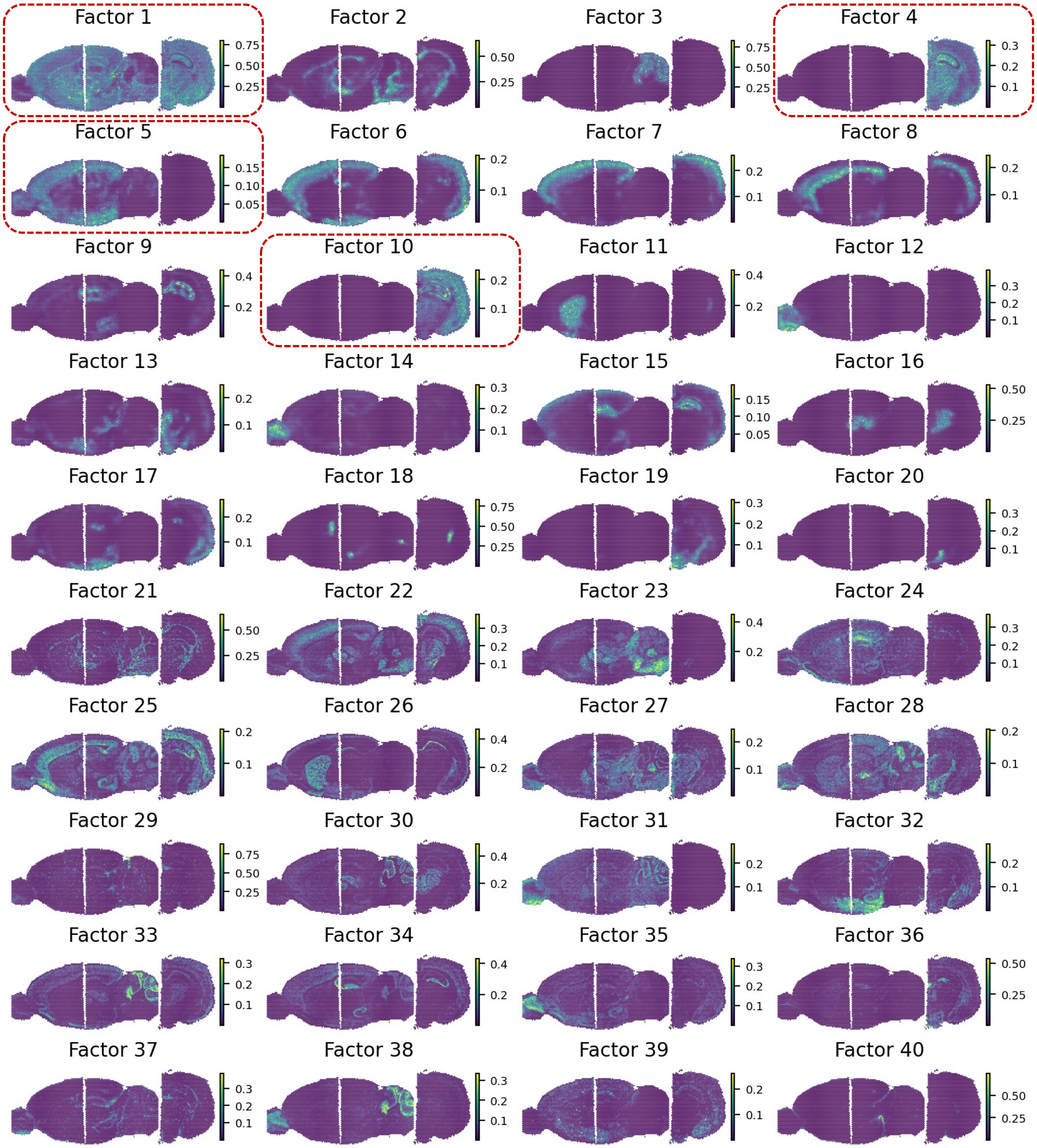


Supplementary Fig. 15: **NSFH's identified spatial factors in the mouse brain dataset.** The spatial distribution of each factor is visualized on three mouse brain slices: sagittal anterior, sagittal posterior, and coronal slices, displayed from left to right, respectively. We marked NSFH's factors that are affected by batch effects among slices, including factors 4, 5 and 10. Additionally, NSFH's factors showed less satisfactory ability to characterize detailed spatial structures in the tissue. For example, factor 1 described a very broad area in mouse brains.

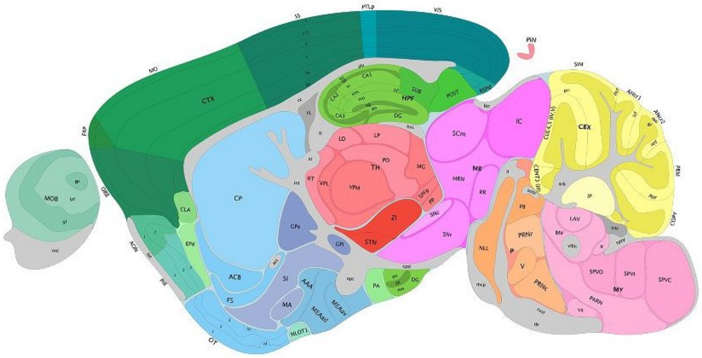

Supplementary Fig. 16: **Mouse brain sagittal anatomical reference from the Allen Reference Atlas - Mouse Brain [2].**

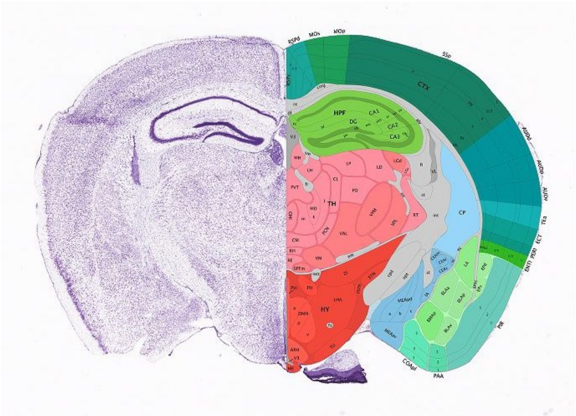

Supplementary Fig. 17: **Mouse brain coronal anatomical reference from the Allen Reference Atlas - Mouse Brain [2].**

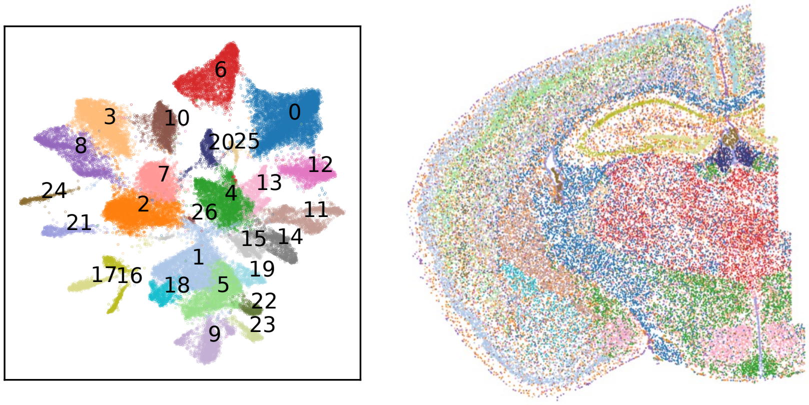


Supplementary Fig. 18: **Spatial domain identification result for the MERFISH mouse brain dataset.** We followed the Scanpy workflow to detect spatial regions for the MERFISH dataset. Both the UMAP plot and the scatter plot are colored by the detected spatial regions.

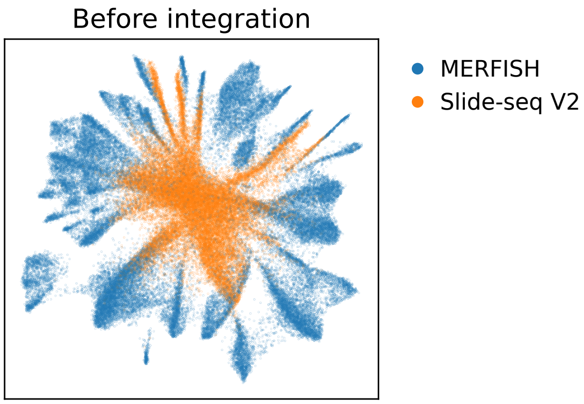


Supplementary Fig. 19: **UMAP visualization of the combined raw data from the MERFISH and Slide-seq V2 datasets prior to integration.** The UMAP plot is colored by the profiling ST technologies of the datasets.

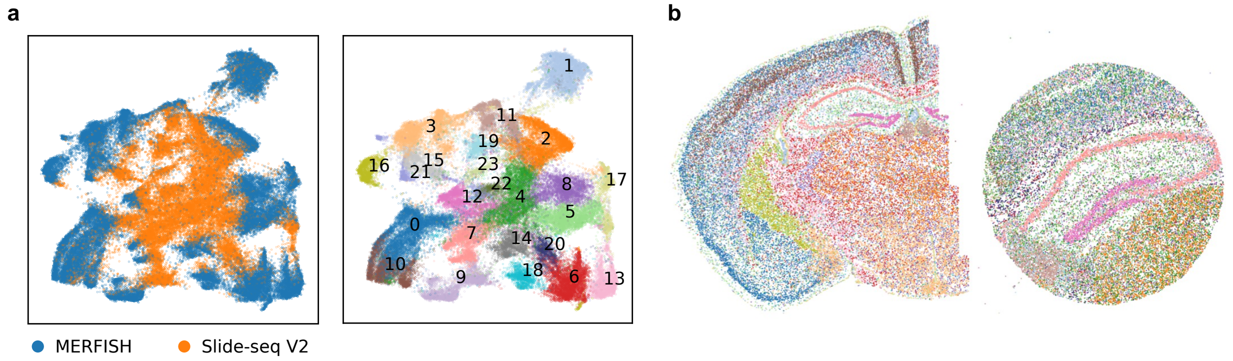


Supplementary Fig. 20: **Cell representations and spatial domain identification result in SpiceMix's joint analysis of the MERFISH and Slide-seq V2 datasets. a**. UMAP plot of cell representations colored by profiling technologies and SpiceMix's identified spatial domain labels, respectively. **b**. Visualization of the identified spatial regions within the two datasets, respectively.

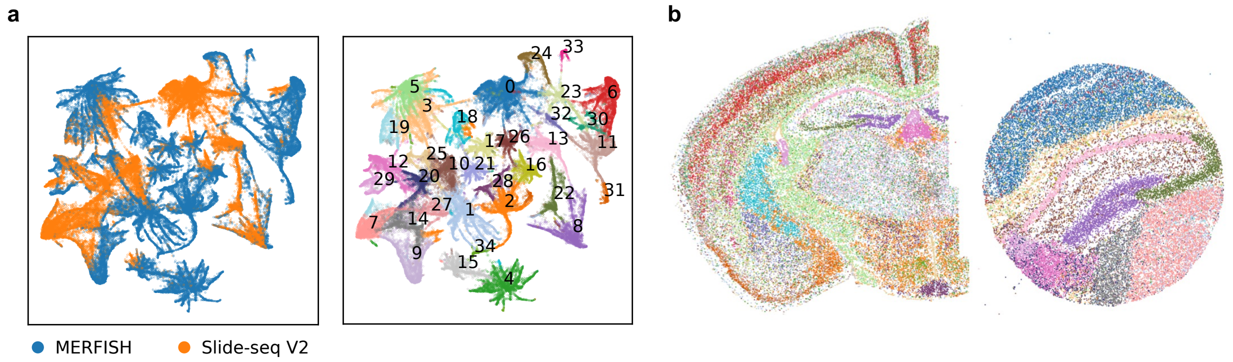


Supplementary Fig. 21: **Cell representations and spatial domain identification result in NSFH's joint analysis of the MERFISH and Slide-seq V2 datasets. a**. UMAP plot of cell representations colored by profiling technologies and NSFH's identified spatial domain labels, respectively. **b**. Visualization of the identified spatial regions within the two datasets, respectively.

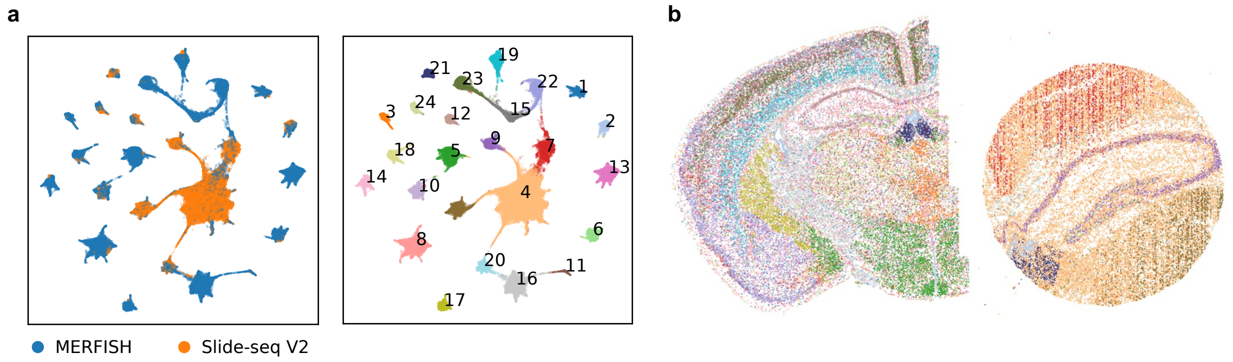


Supplementary Fig. 22: **Cell representations and spatial domain identification result in PRECAST's joint analysis of the MERFISH and Slide-seq V2 datasets. a**. UMAP plot of cell representations colored by profiling technologies and PRECAST's identified spatial domain labels, respectively. **b**. Visualization of the identified spatial regions within the two datasets, respectively.

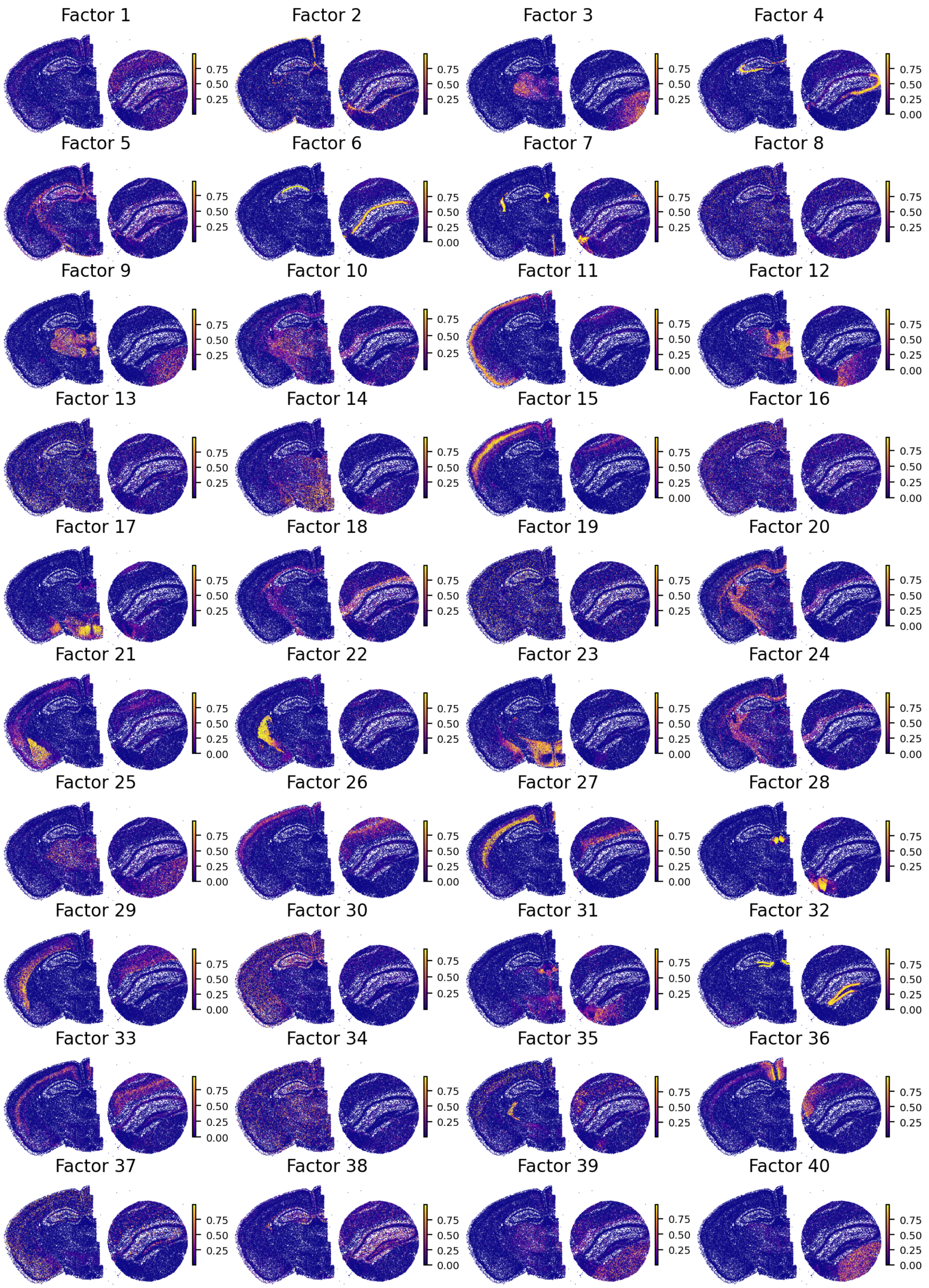


Supplementary Fig. 23: **INSPIRE's identified spatial factors in the joint analysis of the MERFISH and Slide-seq V2 brain datasets.** The spatial distribution of each factor is visualized on the MERFISH and Slide-seq V2 slices, respectively.

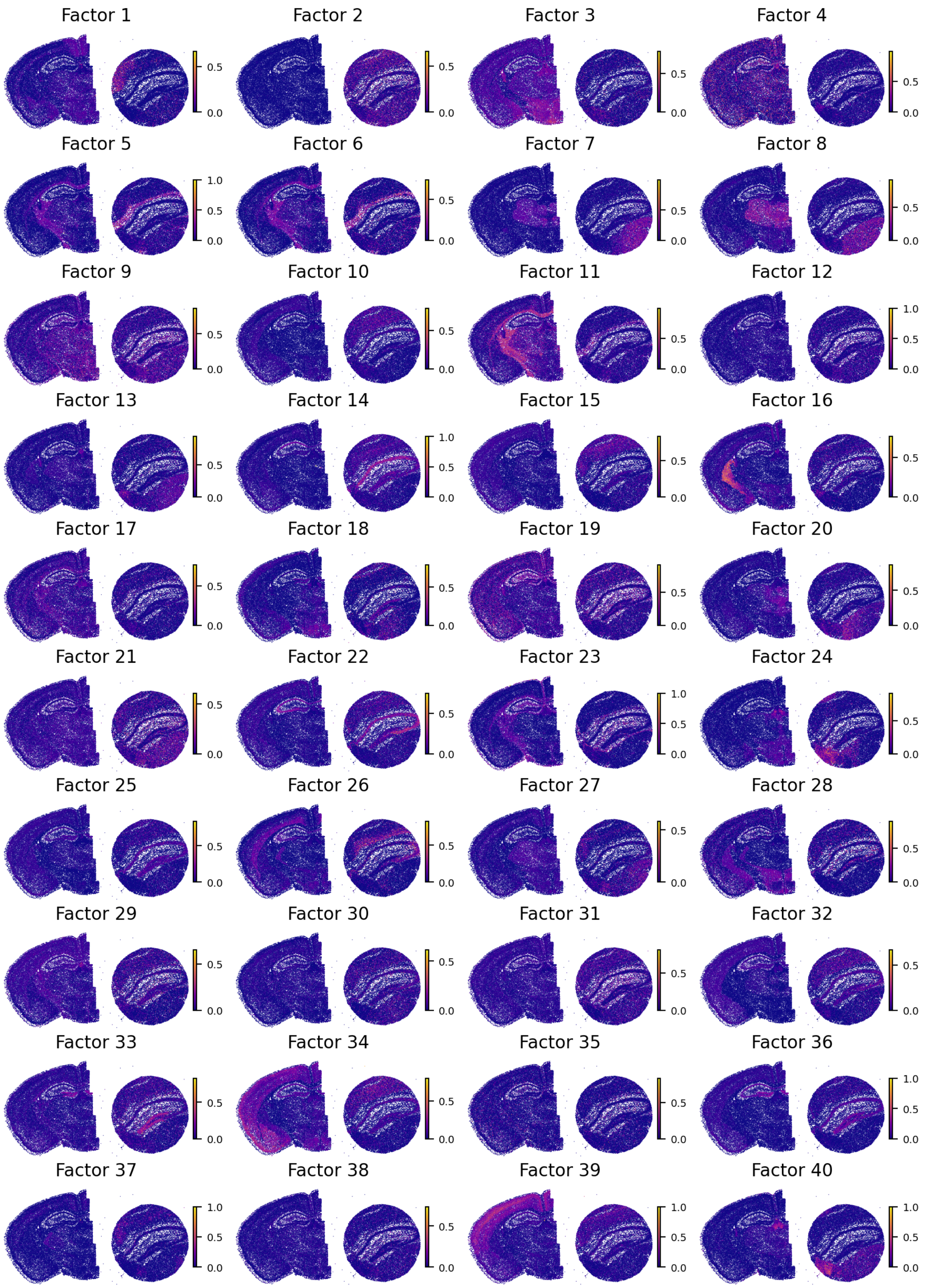


Supplementary Fig. 24: **SpiceMix's identified spatial factors in the joint analysis of the MERFISH and Slide-seq V2 brain datasets.** The spatial distribution of each factor is visualized on the MERFISH and Slide-seq V2 slices, respectively.

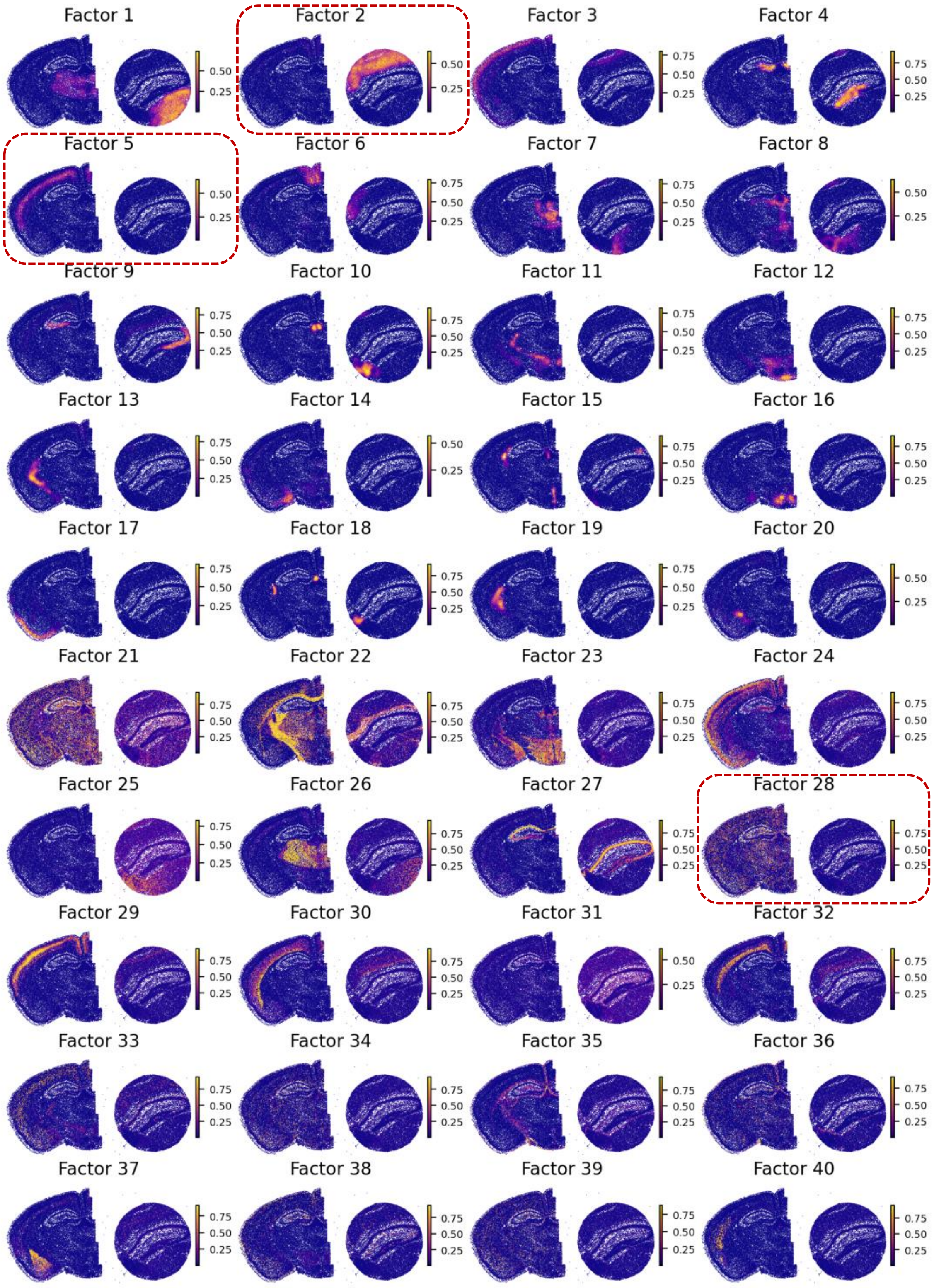


Supplementary Fig. 25: **NSFH's identified spatial factors in the joint analysis of the MERFISH and Slide-seq V2 brain datasets.** The spatial distribution of each factor is visualized on the MERFISH and Slide-seq V2 slices, respectively. We marked NSFH's factors that are affected by technical effects between datasets, including factors 2, 5 and 28.

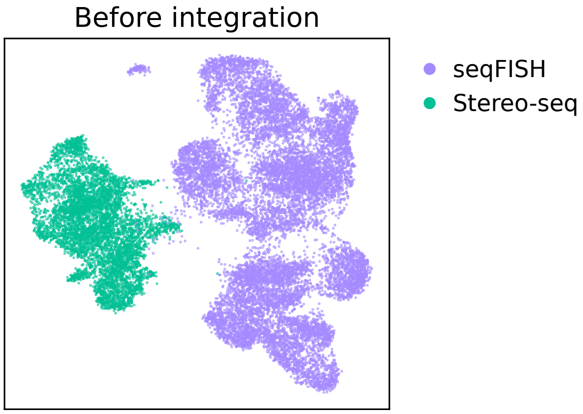


Supplementary Fig. 26: **UMAP visualization of the combined raw data from the seqFISH and Stereo-seq mouse whole-embryo datasets prior to integration.** The UMAP plot is colored by the profiling ST technologies of the datasets.

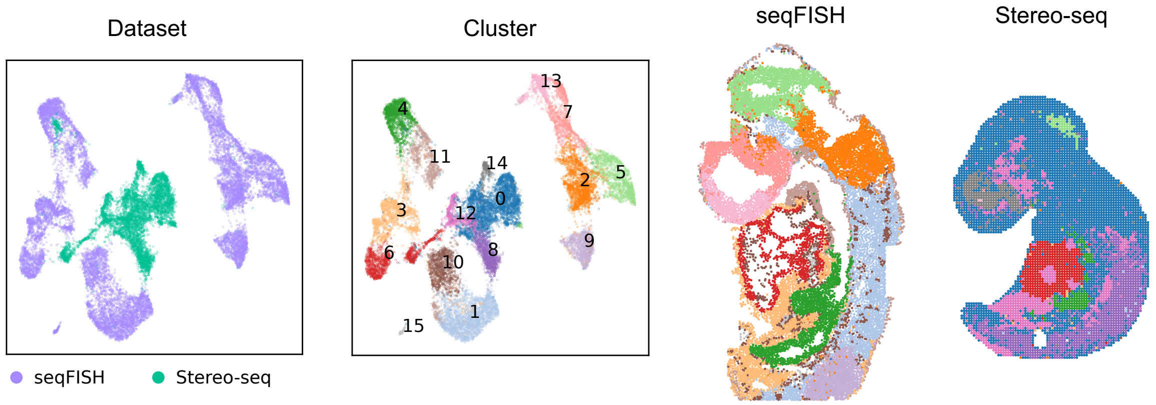


Supplementary Fig. 27: **Cell representations and spatial domain identification result in SpiceMix's joint analysis of the seqFISH and Stereo-seq mouse embryo datasets. a**. UMAP plot of cell representations colored by profiling technologies and SpiceMix's identified spatial domain labels, respectively. **b**. Visualization of the identified spatial regions within the two datasets, respectively.

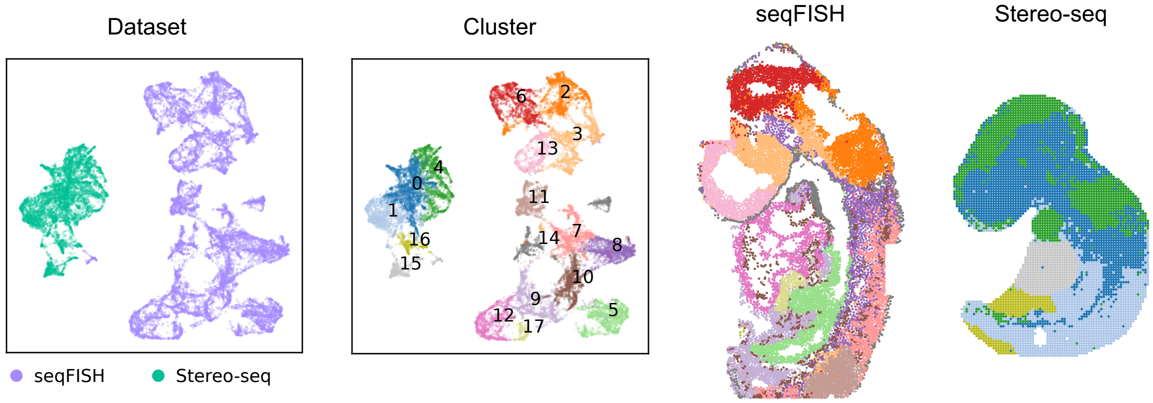


Supplementary Fig. 28: **Cell representations and spatial domain identification result in NSFH's joint analysis of the seqFISH and Stereo-seq mouse embryo datasets. a**. UMAP plot of cell representations colored by profiling technologies and NSFH's identified spatial domain labels, respectively. **b**. Visualization of the identified spatial regions within the two datasets, respectively.

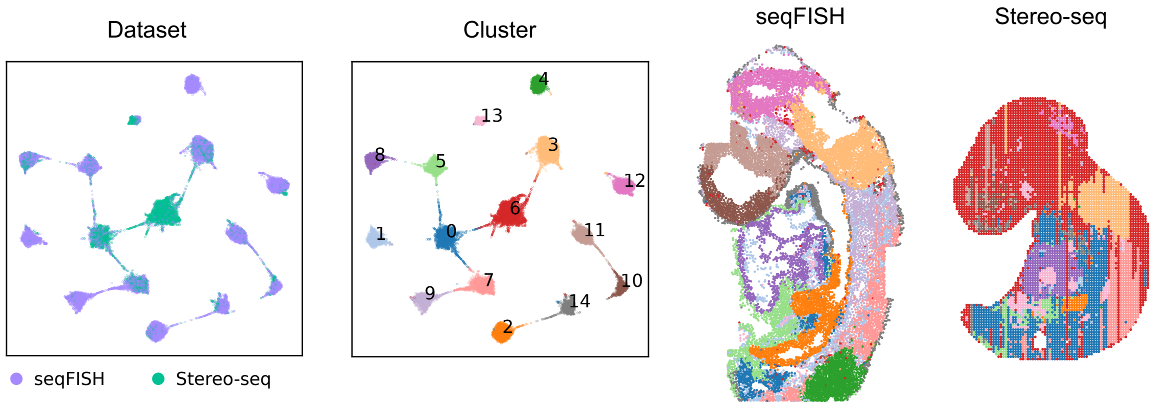


Supplementary Fig. 29: **Cell representations and spatial domain identification result in PRECAST's joint analysis of the seqFISH and Stereo-seq mouse embryo datasets. a**. UMAP plot of cell representations colored by profiling technologies and PRECAST's identified spatial domain labels, respectively. **b**. Visualization of the identified spatial regions within the two datasets, respectively.

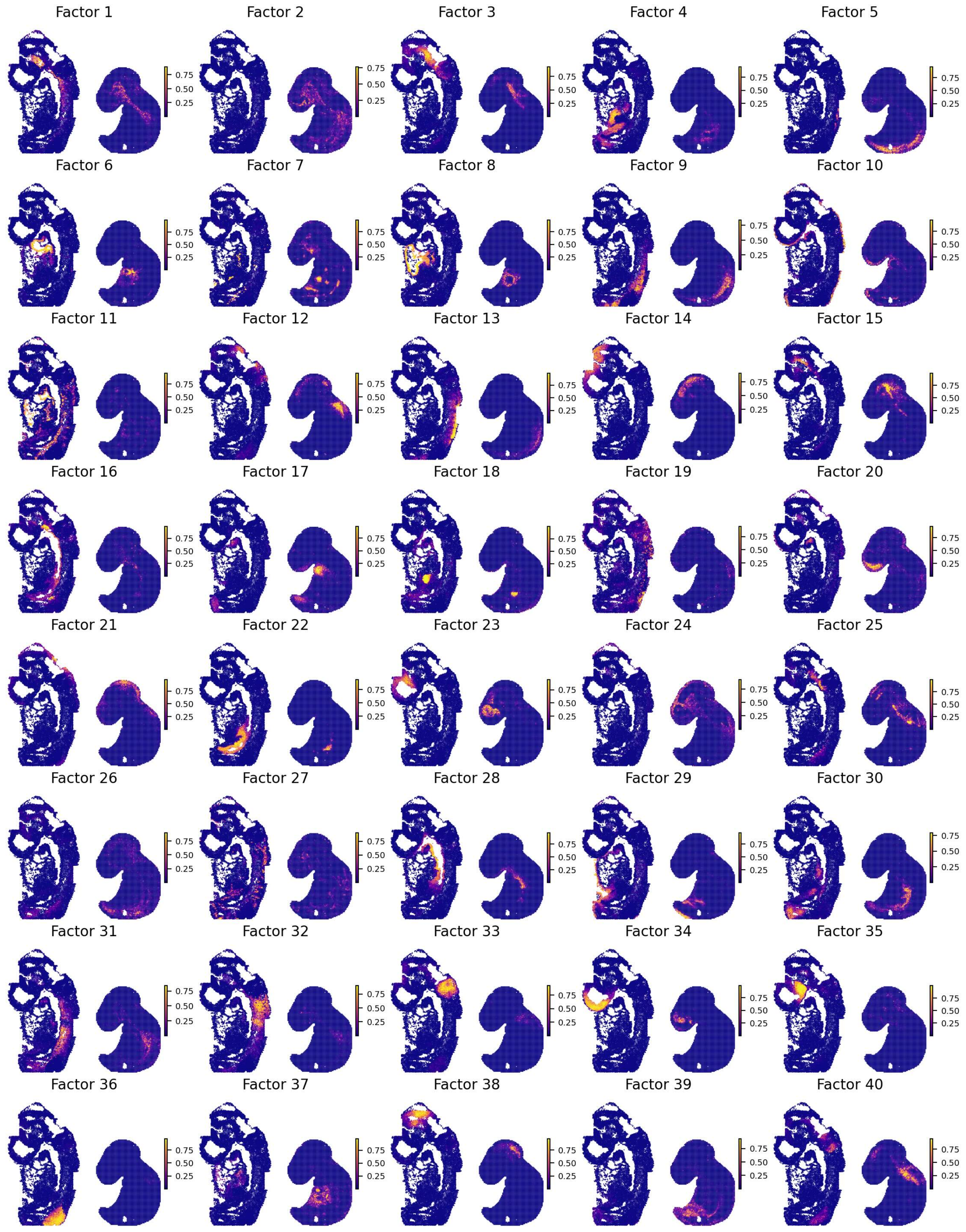


Supplementary Fig. 30: **INSPIRE's identified spatial factors in the joint analysis of the seqFISH and Stereo-seq mouse whole-embryo datasets.** The spatial distribution of each factor is visualized on the seqFISH and Stereo-seq slices, respectively.

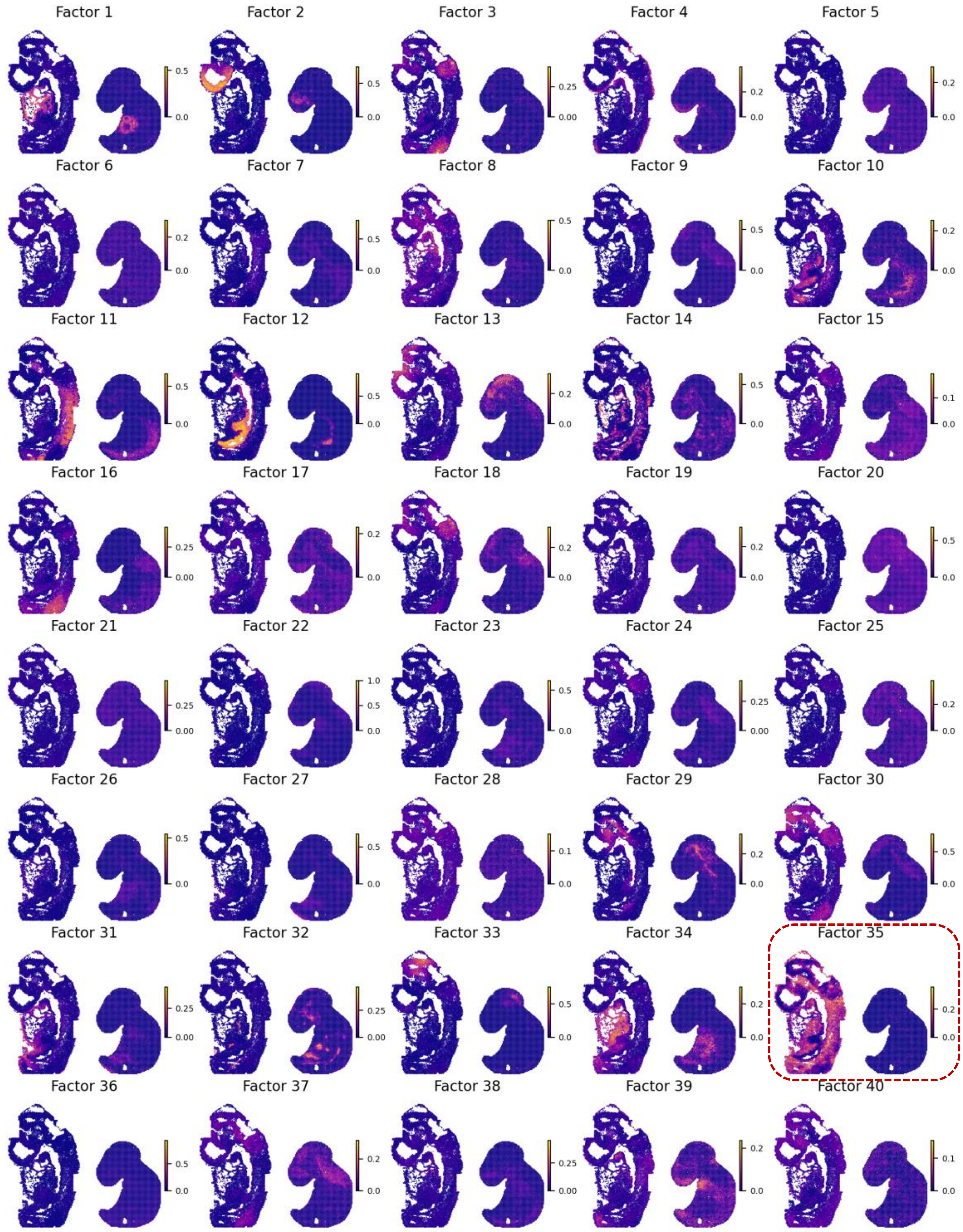


Supplementary Fig. 31: **SpiceMix's identified spatial factors in the joint analysis of the seqFISH and Stereo-seq mouse whole-embryo datasets.** The spatial distribution of each factor is visualized on the seqFISH and Stereo-seq slices, respectively. We marked SpiceMix's factor 35 which is affected by strong technical effects between slices. Additionally, SpiceMix's spatial factors showed limited ability to depict spatial structures in this data analysis.

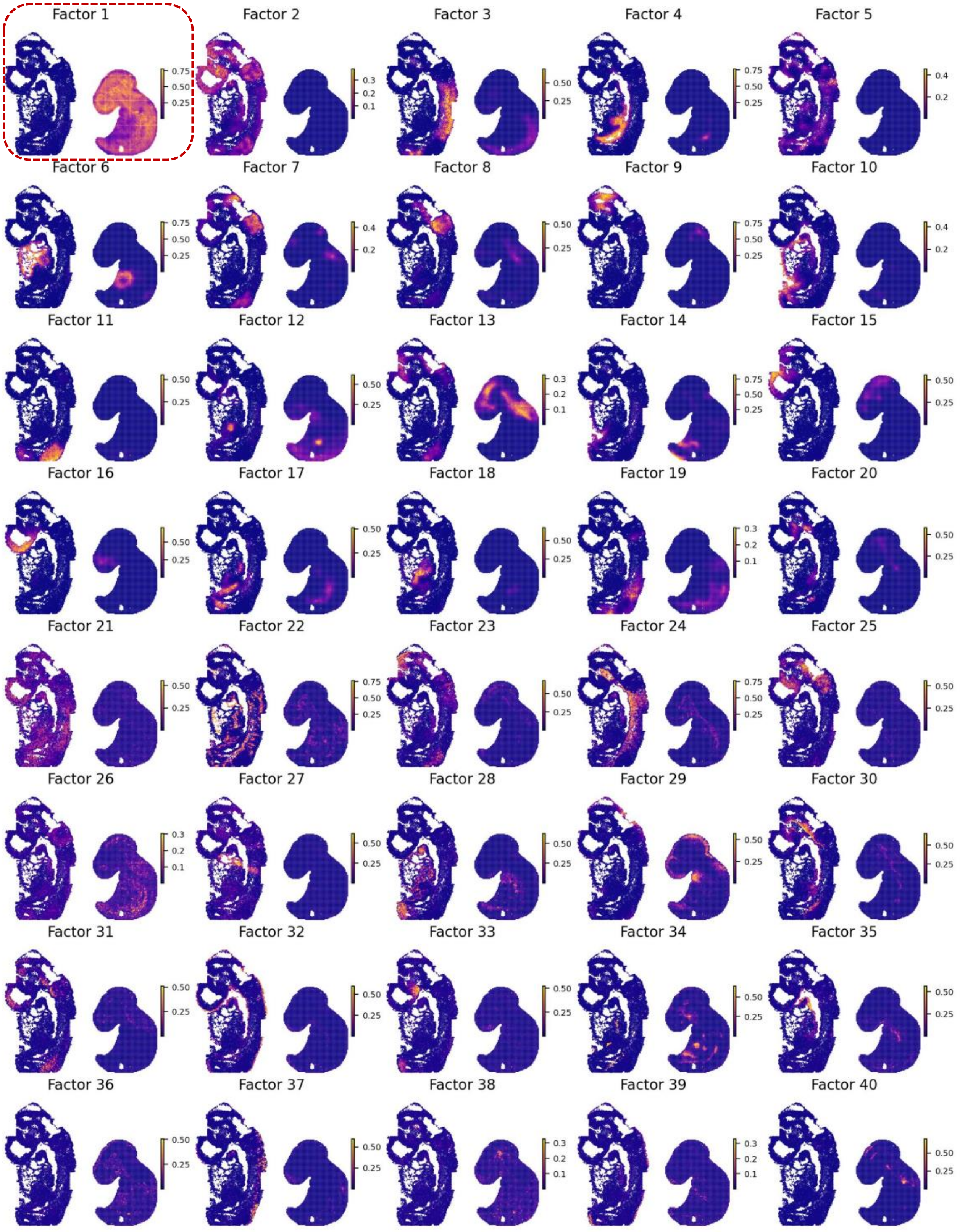


Supplementary Fig. 32: **NSFH's identified spatial factors in the joint analysis of the seqFISH and Stereo-seq mouse whole-embryo datasets.** The spatial distribution of each factor is visualized on the seqFISH and Stereo-seq slices, respectively. We marked NSFH's factor 1 which is confounded by strong technical effects between slices.

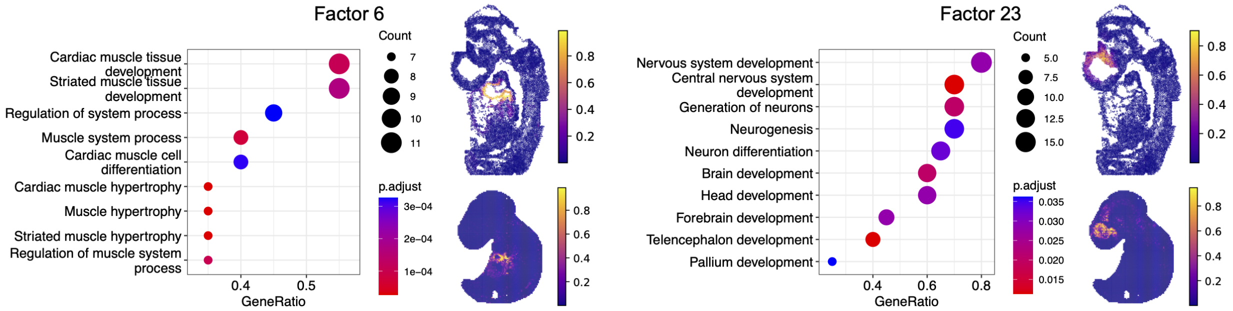


Supplementary Fig. 33: **Significant gene ontology terms and spatial distributions of INSPIRE's spatial factors.** We visualized gene ontology analysis results and spatial distributions of factors 6 (left panel) and 23 (right panel), respectively. Factor 6 is associated with the atria in embryonic hearts and is enriched in pathways related to cardiac muscle tissue development. Factor 23 corresponds to the forebrain in mouse embryos and is enriched in pathways including nervous system development, central nervous system development, and forebrain development.

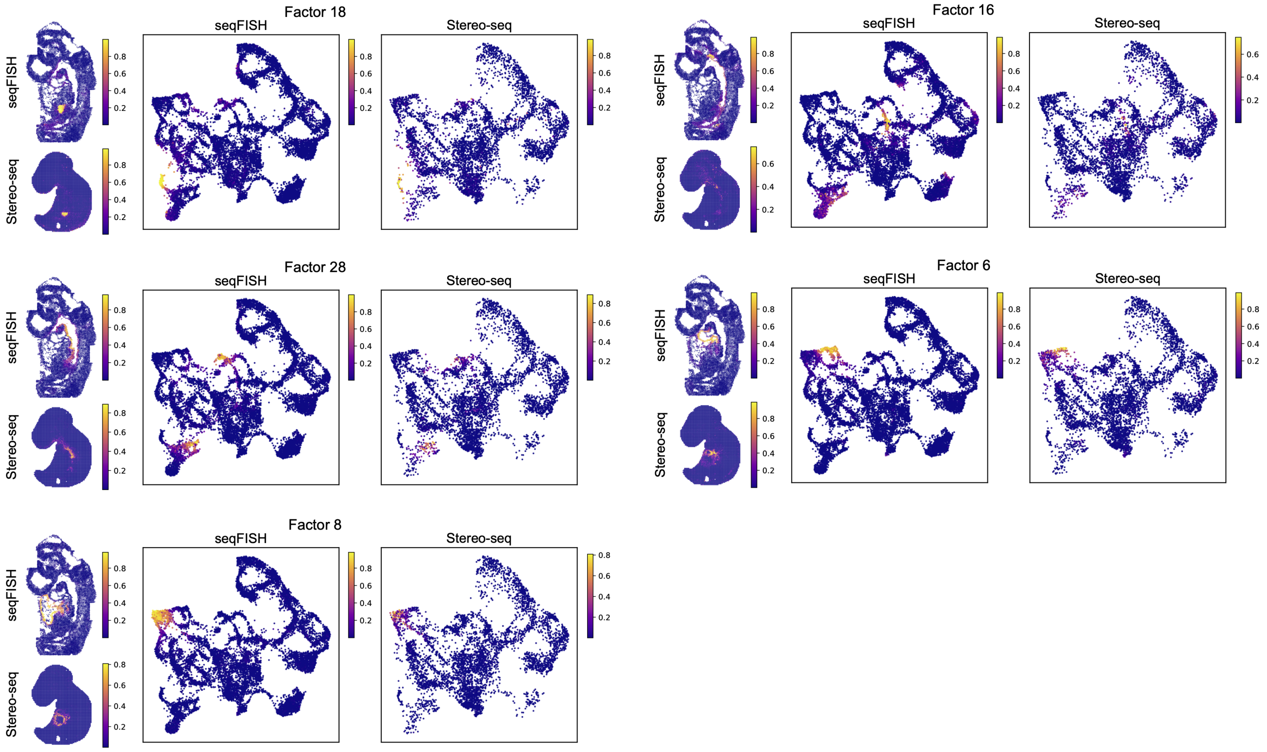


Supplementary Fig. 34: **Precise integrative analysis of the seqFISH and Stereo-seq mouse embryo datasets provided by INSPIRE.** For each factor inferred by INSPIRE, we visualized its spatial distributions across the two datasets, revealing the detailed spatial structures. Additionally, we visualized INSPIRE's aligned cell representations between the datasets, with cells colored according to factor weights. Here, we present five factors as an example to demonstrate that INSPIRE produced accurate cell embeddings that align well across datasets, even at the resolution of fine-grained spatial regions defined by the spatial factors.

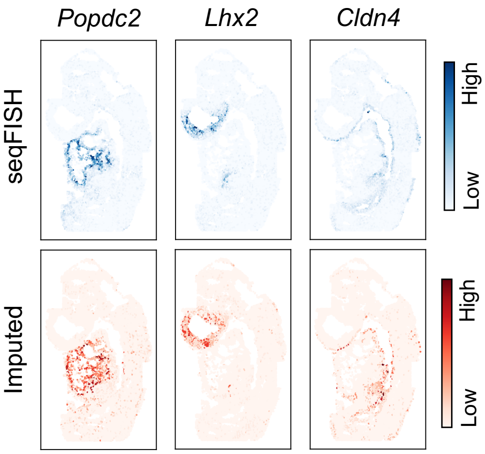


Supplementary Fig. 35: **INSPIRE's imputation of expression levels for genes *Popdc2*, *Lhx2* and *Cldn4* in the seqFISH dataset.** We manually excluded these genes from the seqFISH dataset and imputed their expression levels using the Stereo-seq dataset to demonstrate INSPIRE's gene imputation capability. Top panel shows the actual expression levels measured by seqFISH, while bottom panel presents the imputed expression levels for *Popdc2*, *Lhx2*, and *Cldn4*.

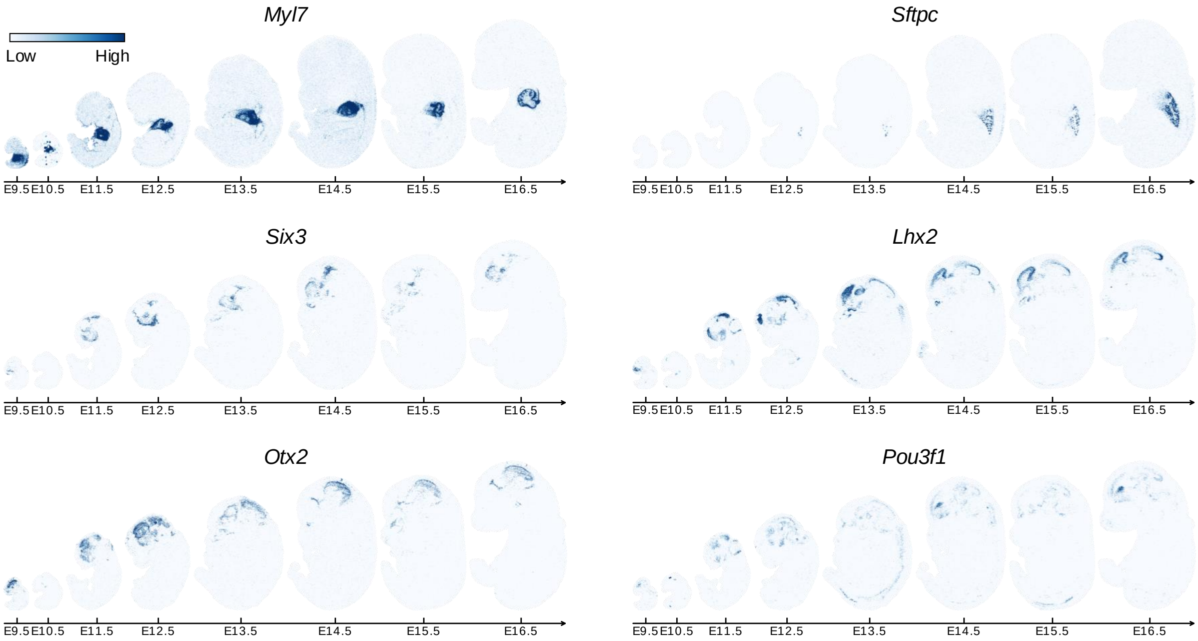


Supplementary Fig. 36: **Spatial expression patterns of marker genes for the major organs and tissues in the embryos across different developmental stages.** *Myl7* and *Sftpc* are marker genes for the heart and the lung, respectively. *Six3*, *Lhx2*, *Otx2* and *Pou3f1* are marker genes for the brain.

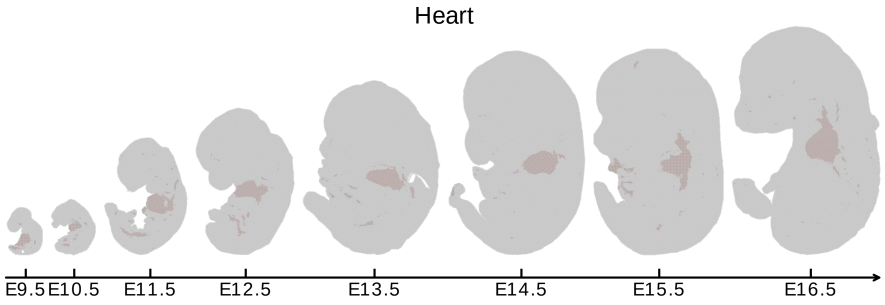


Supplementary Fig. 37: **Visualization of INSPIRE's identified region that represents the heart in the embryos across developmental stages.**

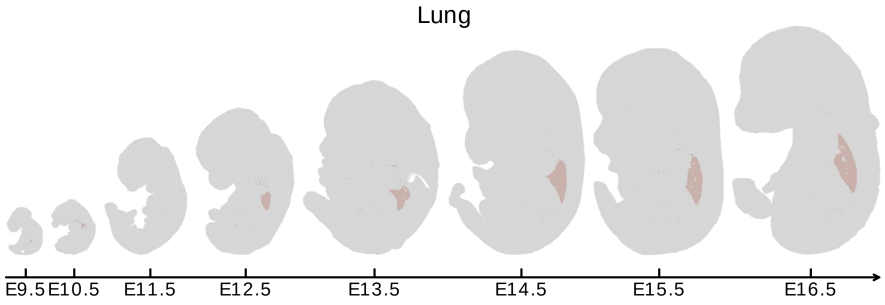


Supplementary Fig. 38: **Visualization of INSPIRE's identified region that represents the lung in the embryos across developmental stages.**

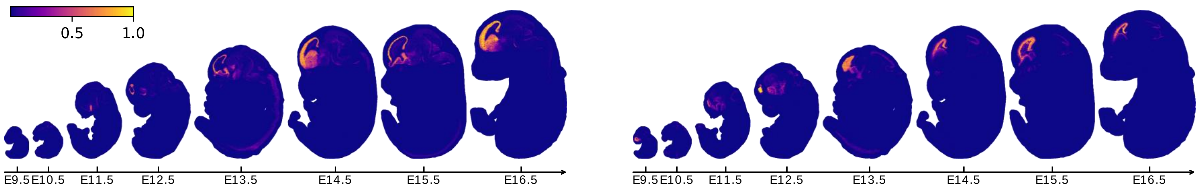


Supplementary Fig. 39: **Spatial distributions of two spatial factors detected by INSPIRE that locate in the embryonic forebrain.** These two factors emerged in the forebrain region by stage E11.5 and progressively developed into a layered structure in the subsequent developmental stages.

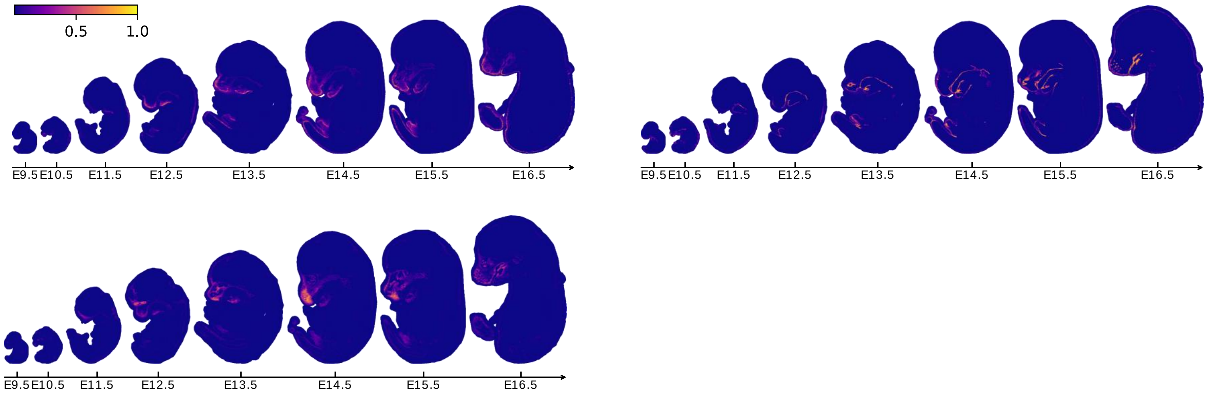


Supplementary Fig. 40: **Spatial distributions of three spatial factors identified by INSPIRE that characterize the spatial structure within embryonic mouth and jaw.** The spatial organization of these three factors became increasingly complex by stages E12.5 and E13.5.

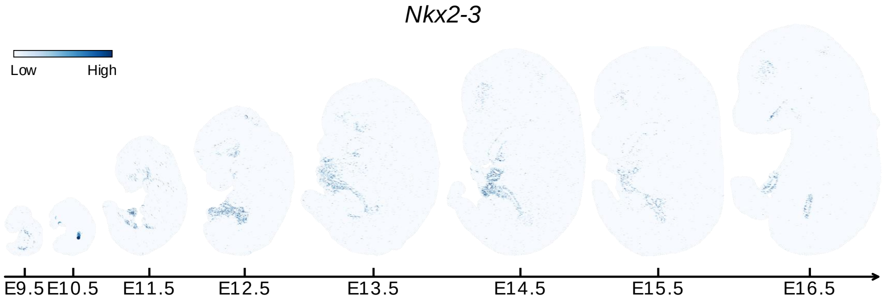


Supplementary Fig. 41: **Spatial expression pattern of gene *Nkx2-3* in the embryos across different developmental stages.**

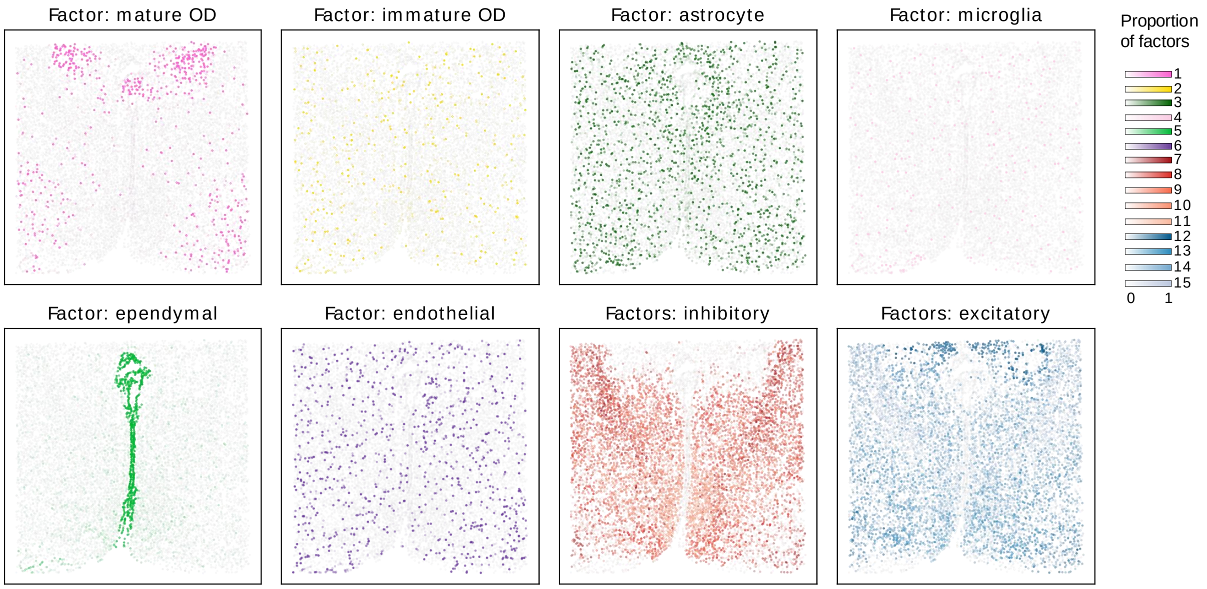


Supplementary Fig. 42: **Spatial distributions of spatial factors visualized on INSPIRE's registered MERFISH slices.** The reliability of INSPIRE's spatial slice registration can be validated based on this result. For example, we have already shown that factor 5 corresponded to ependymal cells. Here, we are able to demonstrate that the spatial distributions of ependymal cells aligned well across the two MERFISH slices, indicating the perfect slice registration.

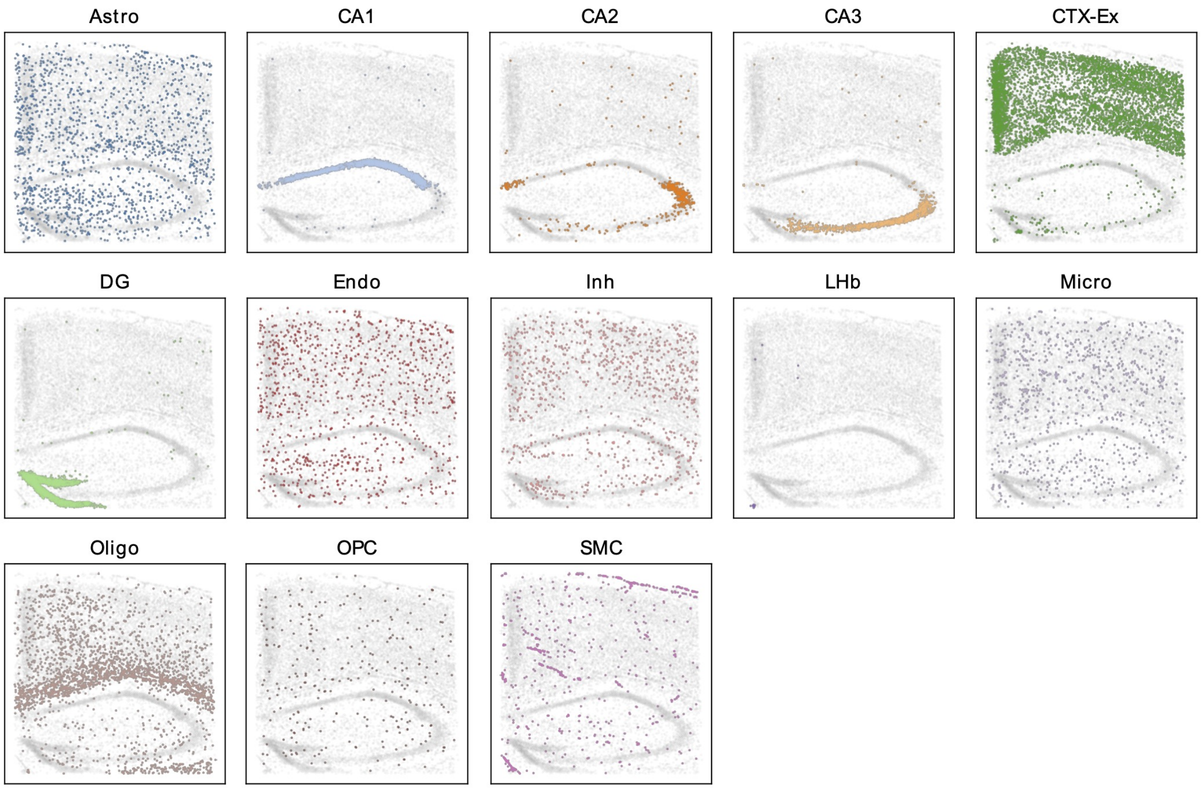


Supplementary Fig. 43: **Spatial distributions of the annotated cell types visualized on INSPIRE's registered STARmap PLUS slices.** The cell type annotations were provided by the original study. Distributions of cell types matched perfectly across the two STARmap PLUS slices, indicating the reliability of INSPIRE's spatial slice registration result.

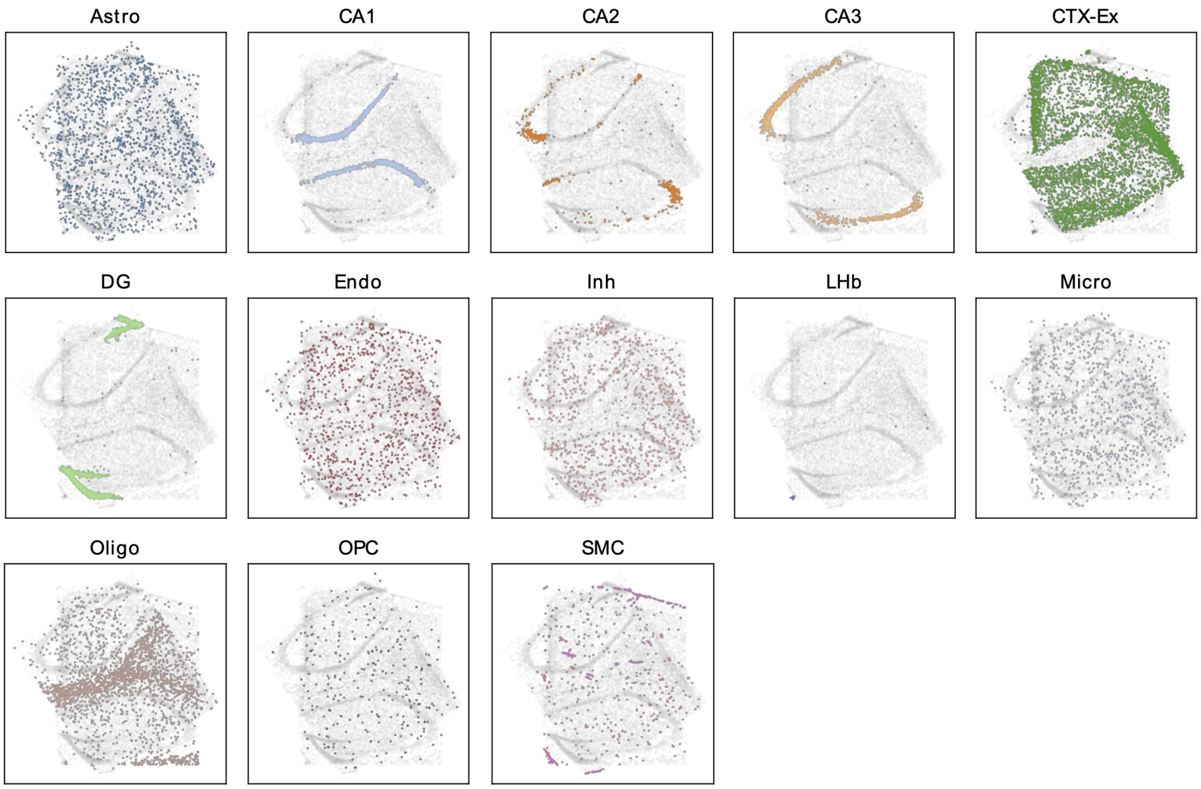


Supplementary Fig. 44: **Spatial distributions of the annotated cell types visualized on PASTE's registered STARmap PLUS slices.**

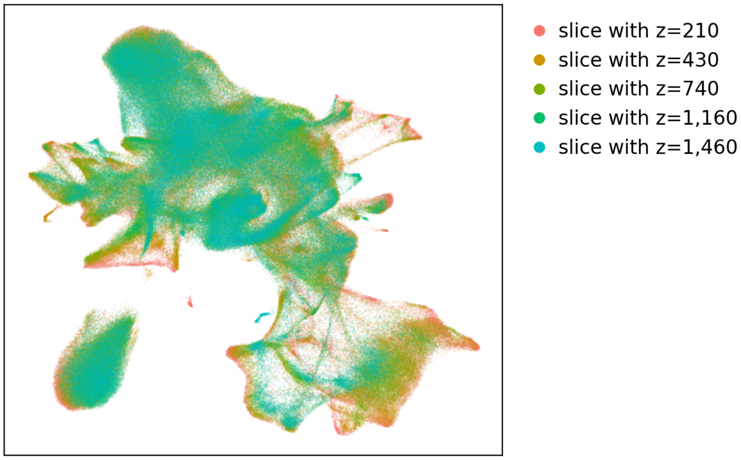


Supplementary Fig. 45: **INSPIRE's learned spot representations in the Stereo-seq data analysis.** The UMAP plot is colored by Stereo-seq slice indices.

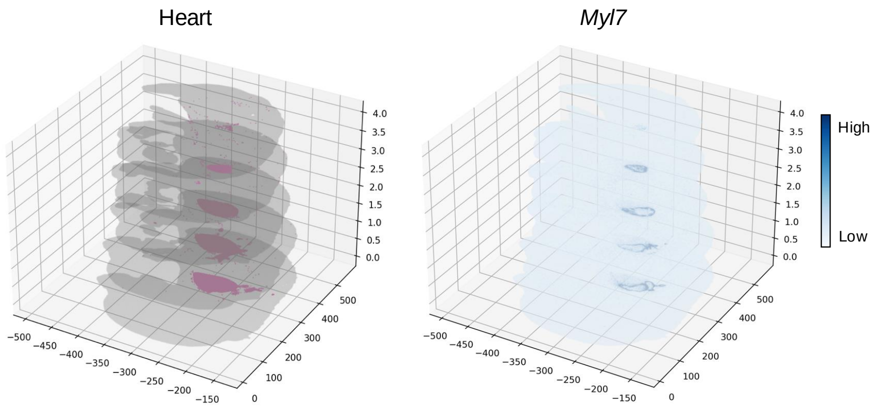


Supplementary Fig. 46: **Visualization of heart structure and distribution of *Myl7* expression on the reconstructed 3D model of the embryo.**

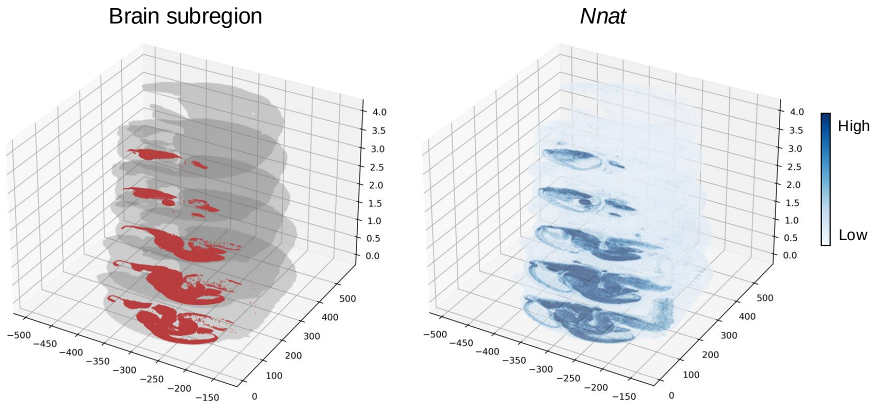


Supplementary Fig. 47: **Visualization of the 3D structure of a brain subregion and the distribution of *Nnat* expression on the reconstructed 3D embryo.**

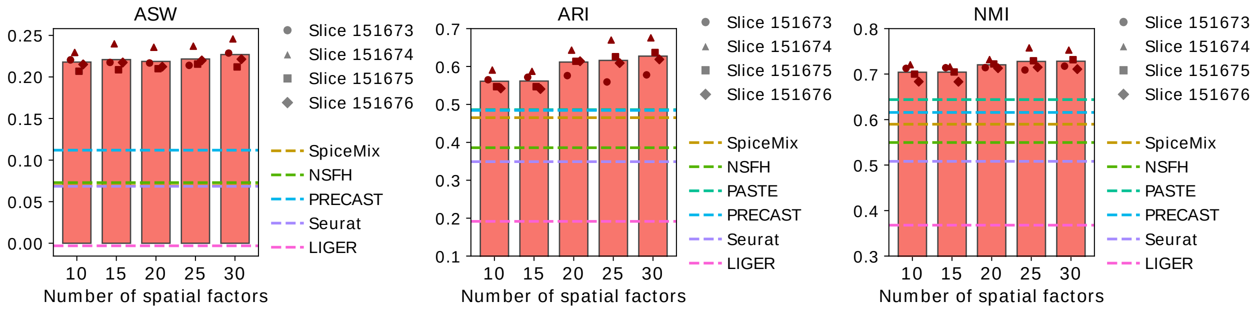


Supplementary Fig. 48: **Impact of the number of spatial factors in NMF on the quality of spot representations in the human DLPFC example.** We ran INSPIRE with number of spatial factors varying from 10 to 30, and evaluated the quality of the learned spot representations using the ASW, ARI, and NMI metrics. Additionally, for each metric, we compared the scores of INSPIRE to those of other methods.

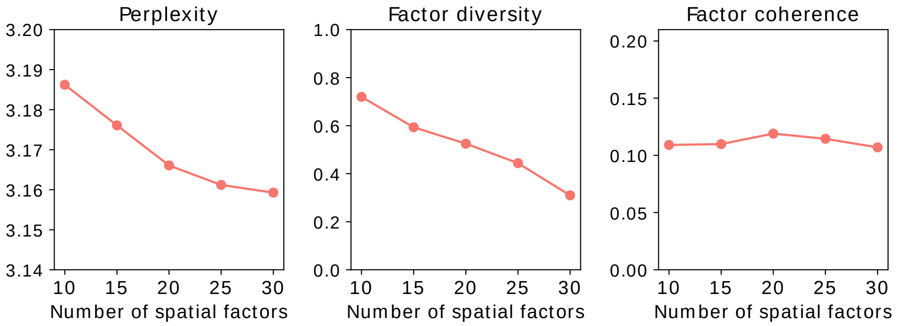


Supplementary Fig. 49: **Impact of the number of spatial factors in NMF on perplexity, factor diversity, and factor coherence scores in the human DLPFC example.** Perplexity measures model fitting accuracy, with lower scores indicating better accuracy. Factor diversity and factor coherence metrics evaluate the diversity and interpretability of spatial factors, respectively. Higher factor diversity and factor coherence scores indicate higher quality of spatial factors.

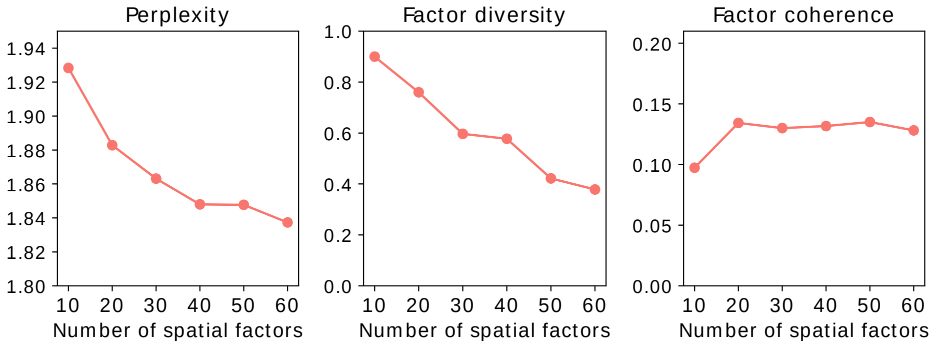


Supplementary Fig. 50: **Impact of the number of spatial factors in NMF on perplexity, factor diversity, and factor coherence scores in the analysis of mouse brain slices with different tissue views.**

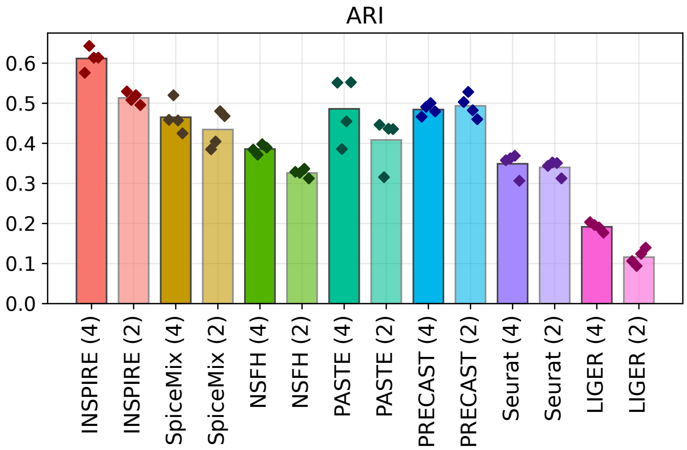


Supplementary Fig. 51: **Comparison of ARI scores between INSPIRE and other methods for four-slice and two-slice analyses on the human DLPFC dataset.** The performance of INSPIRE is denoted as "INSPIRE (4)" for four-slice analysis and "INSPIRE (2)" for two-slice analysis, with analogous labels assigned to the other methods.

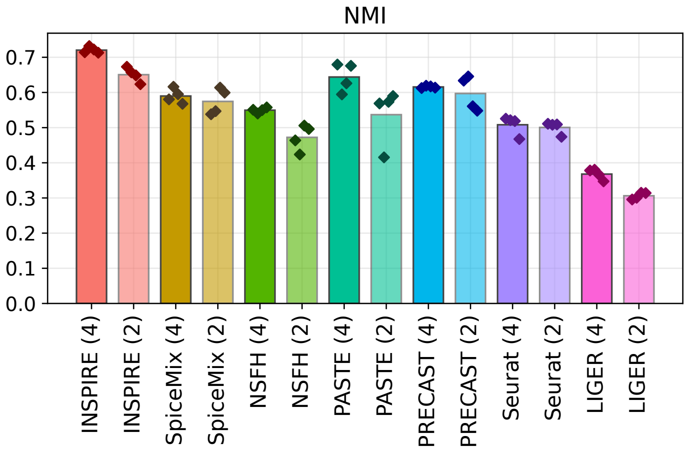


Supplementary Fig. 52: **Comparison of NMI scores between INSPIRE and other methods for four-slice and two-slice analyses on the human DLPFC dataset.**

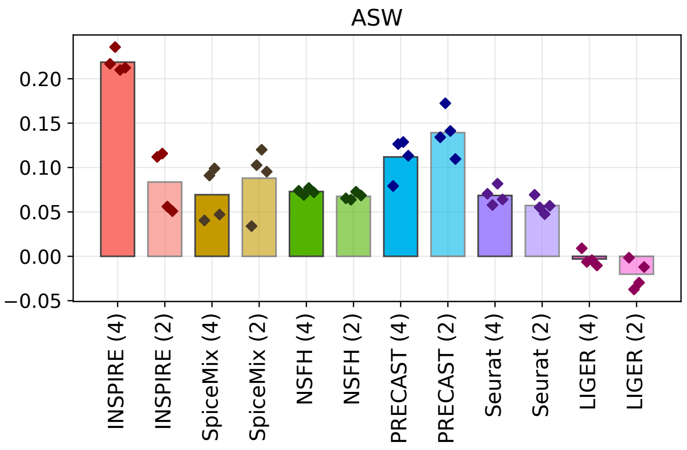


Supplementary Fig. 53: **Comparison of ASW scores between INSPIRE and other methods for four-slice and two-slice analyses on the human DLPFC dataset.**

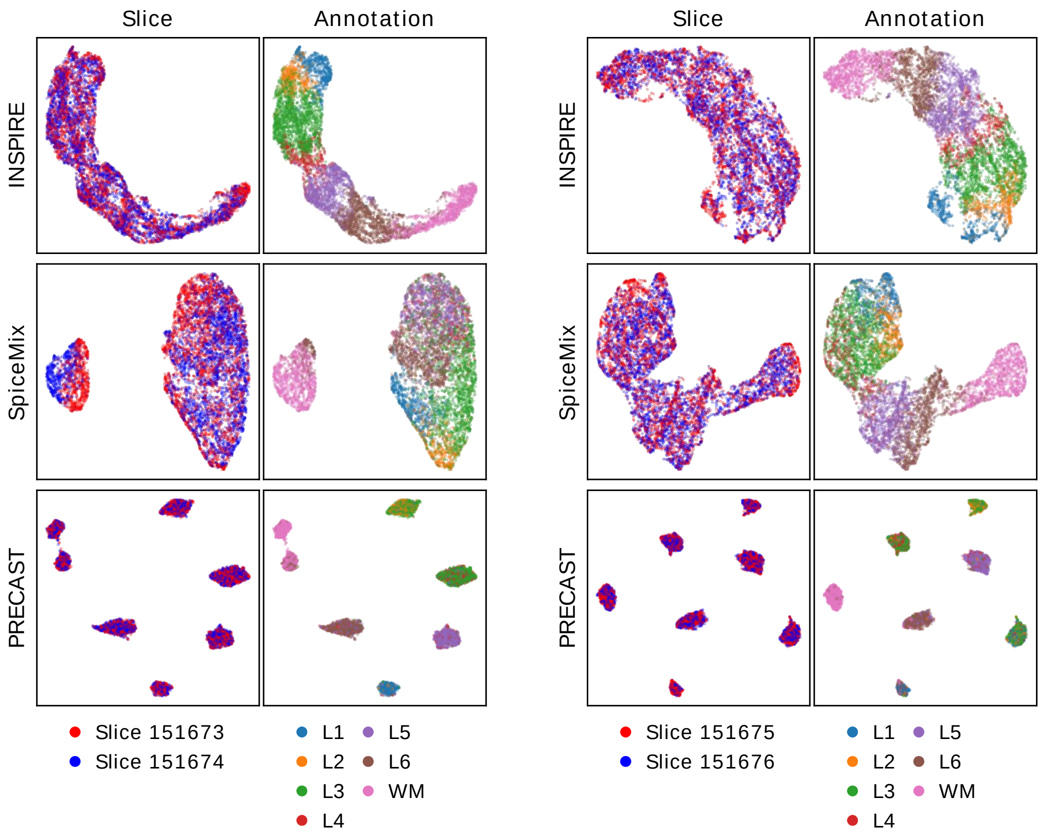


Supplementary Fig. 54: **Comparison of spot representations between INSPIRE and other methods for the two-slice analyses based on the human DLPFC dataset.** Spot representations were visualized using UMAP plots colored by slice index and manual annotation. The left panel displays the integrative analysis results of slices 151673 and 151674, while the right panel shows the integrative analysis results of slices 151675 and 151676.

### Supplementary Tables

| Example | # slices | # spots | # hvgs | Time (h:m:s) | GNN |
|---|---|---|---|---|---|
| Human DLPFC slices (Visium) | 4 | $14,243$ | $1,875$ | $23:44$ | GAT |
| Mouse brain slices with different views (Visium) | 3 | $8,820$ | $3,035$ | $15:49$ | GAT |
| Mouse brain slices across technologies (Slide-seq V2, MERFISH) | 2 | $74,816$ | 936 | $26:21$ | LGCN |
| Mouse embryo slices across technologies (seqFISH, Stereo-seq) | 2 | $20,065$ | 344 | $3:39$ | LGCN |
| Mouse embryo, developmental stages (Stereo-seq) | 8 | $514,415$ | $2,366$ | $1:19:27$ | LGCN |
| Mouse hypothalamus, spatial registration (MERFISH) | 2 | $11,045$ | 155 | $2:07$ | LGCN |
| Mouse hippocampus, spatial registration (STARmap PLUS) | 2 | $16,388$ | $2,766$ | $29:35$ | LGCN |
| Mouse embryo, 3D reconstruction (Stereo-seq) | 5 | $567,381$ | $2,966$ | $1:17:00$ | LGCN |

Supplementary Table 1: **Number of slices, number of spots, number of highly variable genes, INSPIRE's utilized GNN type and training time for different experiments.** For GNN type, GAT refers to networks with graph attention layers, while LGCN refers to networks with lightweight graph convolutional layers.

| Method | Spatial information modeling | Multiple dataset analysis | Unwanted variation removal | Gene program inference | Spatial registration | Mini-batch training |
|---|---|---|---|---|---|---|
| Seurat | No | Yes | Yes | No | No | No |
| LIGER | No | Yes | Yes | No | No | No |
| PASTE | Yes | Yes | No | No | Yes | No |
| PRECAST | Yes | Yes | Yes | No | No | No |
| SpiceMix | Yes | No | No | Yes | No | Yes |
| NSFH | Yes | No | No | Yes | No | Yes |
| INSPIRE | Yes | Yes | Yes | Yes | Yes | Yes |

Supplementary Table 2: **Summary of INSPIRE and compared state-of-the-art methods.** Unwanted variation removal refers to the ability of methods to produce integrated representations of cells or spatial spots across datasets, enabling effective downstream analyses such as spatial trajectory inference. Gene program inference refers to the ability of methods to uncover the organization of biological processes and the distribution of fine-grained cell types in tissues, along with their associated gene programs.

### Supplementary Notes

#### Supplementary Note 1: Examining the impact of varying spatial factor numbers on spot representation performance

In the human DLPFC example, we manually excluded the NMF component from INSPIRE, referring to this version as “INSPIRE (w/o NMF)”. Compared to INSPIRE (w/o NMF), INSPIRE’s spot representation consistently showed higher scores for all three metrics, including ASW, ARI, and NMI (Supplementary Fig. 4). This finding suggests that the NMF component in INSPIRE contributes to the improved accuracy in the latent space.

We then explored how varying the number of spatial factors in INSPIRE influences the quality of spot representations. We tested with spatial factor numbers set to 10, 15, 20, 25, and 30. For each setting, we measured ASW, ARI, and NMI scores to assess spot representation performance. As shown in Supplementary Fig. 48, these scores remained reasonably stable across different spatial factor numbers, with a slight increase in all three metrics as the number of spatial factors increased.

#### Supplementary Note 2: Choice of the number of spatial factors in INSPIRE

The number of spatial factors is a required input for INSPIRE. To determine the appropriate number of spatial factors for specific ST datasets, we can calculate three metrics: perplexity, factor diversity, and factor coherence. Perplexity is defined as:

$$\text{Perplexity} = \exp\left(-\frac{\log(p(\text{model}_K))}{\sum_{s=1}^{s}\sum_{i=1}^{N_s}\sum_{g=1}^{G} y_{i,g}^{s}}\right),$$

where $p(\text{model}_K)$ represents the likelihood of the datasets under INSPIRE’s model with a given number of spatial factors $K$, and $y_{i,g}^{s}$ is the expression level of gene $g$ in spot or cell $i$ from slice $s$. A lower perplexity indicates a better fit of the model to the datasets.

The other two metrics, factor diversity and factor coherence, evaluate the quality of the learned spatial factors and are detailed in the Methods section. In brief, factor diversity measures the percentage of unique genes associated with each factor, with a higher diversity score indicating more varied factors. Factor coherence assesses the interpretability of factors by evaluating the co-expression of genes associated with the same factor across cells or spots. A higher coherence score indicates better interpretability of spatial factors.

When analyzing ST datasets, we can fit a series of INSPIRE models with varying numbers of spatial factors $K$. By examining the relationship between $K$ and the three metric scores, we can derive empirical insights into selecting the appropriate number of spatial factors. Specifically, we explored these relationships using human DLPFC slices [1] and mouse brain slices with different views [3, 4, 5]. In the human DLPFC example, $K$ was varied from 10 to 30 in increments of 5, while in the mouse brain example, $K$ was varied from 10 to 60 in increments of 10. In both cases, the factor coherence score remained relatively stable across different values of $K$. However, as $K$ increased, both perplexity and factor diversity scores decreased (Supplementary Figs. 49 and 50). This trend suggests that while a higher number of spatial factors improves model fit to the ST datasets, it also leads to less diverse spatial factors. For the human DLPFC slices, which represent only a subregion of the brain, the factor diversity score dropped below 50% when $K$ reached 25. In contrast, for the mouse brain data, which encompasses multiple complementary views of the brain, the factor diversity score remained above 50% even when $K$ increased to 40. This indicates that the appropriate number of spatial factors depends on the scale of the ST slices: larger tissue regions with more complex spatial architectures require a higher number of spatial factors.

Based on empirical observations, we recommend using $K = 20$ for analyzing a subregion of an organ, $K = 40$ for a complex organ, and $K = 60$ for a whole organism. Alternatively, the INSPIRE model can be run with different values of $K$, and we recommend selecting the largest $K$ such that factor diversity exceeds a specified threshold. By default, we suggest a threshold of 50%.

#### Supplementary Note 3: Additional benchmarking results

In a benchmarking study on the human DLPFC dataset, we compared INSPIRE with other state-of-the-art methods for joint analysis of the four DLPFC slices. The results showed that INSPIRE consistently achieved superior data integration, with the highest ARI, NMI, and ASW scores. For a more comprehensive benchmarking, we also performed joint analyses on two slices at a time, including an integrative analysis of slices 151673 and 151674, as well as an integrative analysis of slices 151675 and 151676. INSPIRE's four-slice analysis consistently outperformed its two-slice analysis, demonstrating its ability to effectively leverage information across multiple slices (Supplementary Figs. 51, 52, and 53). Moreover, INSPIRE's two-slice analysis achieved superior spatial domain identification compared to all other methods, as measured by ARI and

NMI. However, PRECAST achieved the highest ASW score in the two-slice analysis, likely due to its use of a Gaussian mixture model to cluster latent representations of spatial spots. This approach enforces a pre-specified number of clusters in the latent space, encouraging spots with similar gene expression profiles to cluster closely, resulting in high ASW scores. However, a limitation of this method is that it tends to divide biologically meaningful continuous trajectories into disjoint clusters, potentially confounding biological insights (Supplementary Fig. 54).

[3] Mouse brain sagittal anterior section by Visium from 10X Genomics. https://www.10xgenomics.com/datasets/mouse-brain-serial-section-2-sagittal-anterior-1-standard-1-0-0.

[4] Mouse brain sagittal posterior section by Visium from 10X Genomics. https://www.10xgenomics.com/datasets/mouse-brain-serial-section-2-sagittal-posterior-1-standard-1-0-0.

[5] Mouse brain coronal section by Visium from 10X Genomics. https://www.10xgenomics.com/datasets/mouse-brain-section-coronal-1-standard-1-1-0.
